# Podocyte exopher-formation as a novel pathomechanism in membranous nephropathy

**DOI:** 10.1101/2024.04.04.588146

**Authors:** Karen Lahme, Wiebke Sachs, Sarah Froembling, Michael Brehler, Desiree Loreth, Kristin Surmann, Simone Gaffling, Uta Wedekind, Vincent Böttcher-Dierks, Marie R. Adler, Pablo J. Sáez, Christian Conze, Roland Thünauer, Sinah Skuza, Karen Neitzel, Stephanie Zieliniski, Johannes Brand, Stefan Bonn, Stephan Michalik, Uwe Völker, Marina Zimmermann, Thorsten Wiech, Tobias N. Meyer, Lars Fester, Catherine Meyer-Schwesinger

## Abstract

**Background:** Membranous nephropathy (MN) is caused by autoantibody binding to podocyte foot process antigens such as THSD7A and PLA_2_R1. The mechanisms of the glomerular antigen/autoantibody deposition and clearance are unknown.

**Methods:** We explore the origin and significance of glomerular accumulations in (1) diagnostic and follow-up biospecimens from THSD7A^+^ and PLA_2_R1^+^-MN patients compared to nephrotic non-MN patients, and (2) in experimental models of THSD7A^+^-MN.

**Results:** We discovered podocyte exophers as correlates of histological antigen/autoantibody aggregates found in the glomerular urinary space of MN patients. Exopher vesicle formation represents a novel form of toxic protein aggregate removal in *Caenorhabditis elegans* neurons. In MN patients, podocytes released exophers to the urine. Enrichment of exophers from MN patient urines established them as a glomerular exit route for antigens and bound autoantibody. Exophers also carried disease-associated proteins such as complement and provided a molecular imprint of podocyte injury pathways. In experimental THSD7A^+^-MN, exophers were formed from podocyte processes and cell body. Their formation involved the translocation of antigen/autoantibody from the subepithelial to the urinary side of podocyte plasma membranes. Urinary exopher-release correlated with lower albuminuria and lower glomerular antigen/autoantibody burden. In MN patients the prospective monitoring of urinary exopher abundance and of exopher-bound autoantibodies was additive in the assessment of immunologic MN activity.

**Conclusions:** Exopher-formation and release is a novel pathomechanism in MN to remove antigen/autoantibody aggregates from the podocyte. Tracking exopher-release will add a non-invasive diagnostic tool with prognostic potential to clinical diagnostics and follow-up of MN patients.

## INTRODUCTION

Idiopathic membranous nephropathy (MN) is an autoimmune disease of kidney podocytes. Podocytes are essential cells of the filtration barrier to blood. In MN, circulating autoantibodies are directed against podocyte foot process proteins, such as the M-type phospholipase A2 receptor (PLA_2_R1)^1^ in 80% and thrombospondin type-1 domain-containing 7 A (THSD7A)^2^ in 5% of patients. Based on clinical correlative observations, the causes of autoimmunity are likely to be multiple in MN^3,4,5,6,7^, as mirrored by the multitude of recently discovered potential MN autoantigens (reviewed in^7–9^). In the current concept of MN, the circulating autoantibodies mostly of the huIgG4 type cause the disease^10,11^ by complement-dependent^12,13^ as well as independent mechanisms^14–16^. The ensuing autoantibody/antigen reaction leads to their pathognomonic glomerular deposition as immune complexes, lastly leading to glomerular filtration barrier alterations and nephrotic range proteinuria^17^. The underlying mechanisms and clinical significance of glomerular aggregate formation in MN are unknown.

In the current study we discover exophers as the pathobiological correlate of glomerular urinary space aggregates in human and experimental MN. Exophers represent a novel type of extracellular vesicle (EV) recently discovered in *Caenorhabditis elegans* neurons^18^ and body wall muscles^19^. Exophers extrude from juxtanuclear plasma membrane areas^20^ as long nanotubules which can reach up to 4 µm in diameter at the distal end^21^. In *C. elegans*, exopher-formation depends on cellular stressors^17,24^ and mechanistically involves the delivery of cargo to aggresome-like organelles^20^. Exopher-like processes have morphologically been reported in mammalian systems and are thought to represent a protective strategy to remove proteotoxic protein aggregates^21,27^ and dysfunctional organelles such as mitochondria^22,24^. Phagocytosis of released exophers by neighboring cells is considered neuroprotective^18^. This export of proteotoxic protein aggregates may be conserved in flies^28,29^ and mammalian neurons^30^. Exophers as such have not been identified in humans yet.

Here we hypothesize that understanding the mechanisms that govern glomerular antigen/autoantibody deposition and clearance in MN will open new non-invasive diagnostic avenues with prognostic potential, as the podocytes’ ability to deal with the disease initiating autoantibodies is included. In comparative studies to nephrotic non-MN patients, we set out to establish glomerular aggregates as disease-specific exophers in MN. Using experimental podocyte and mouse models of THSD7A^+^-MN we investigate the dynamics and disease modulating effects of exopher-formation and release. The podocytes’ ability to form and release exophers to the urine is assessed in retrospective and prospective patient cohorts, and its additive clinical value in disease monitoring is shown.

## RESULTS

### Exophers are the pathobiological correlate of glomerular urinary space aggregates in MN

Our analyses of diagnostic biopsies and diagnostic/consecutive urine samples from a THSD7A^+^-MN patient (***Fig. 1***) and from three PLA_2_R1^+^-MN patients (***Fig. S1***) revealed human exopher-formation as an antigen-overarching principle in MN, compared to nephrotic non-MN patients (two minimal change disease, two primary FSGS, one tubulo-toxic kidney injury, and one IgA nephritis patient). The blood and urine parameters of the patients are summarized in ***Tables S1***, ***S2***.

**Figure 1:**
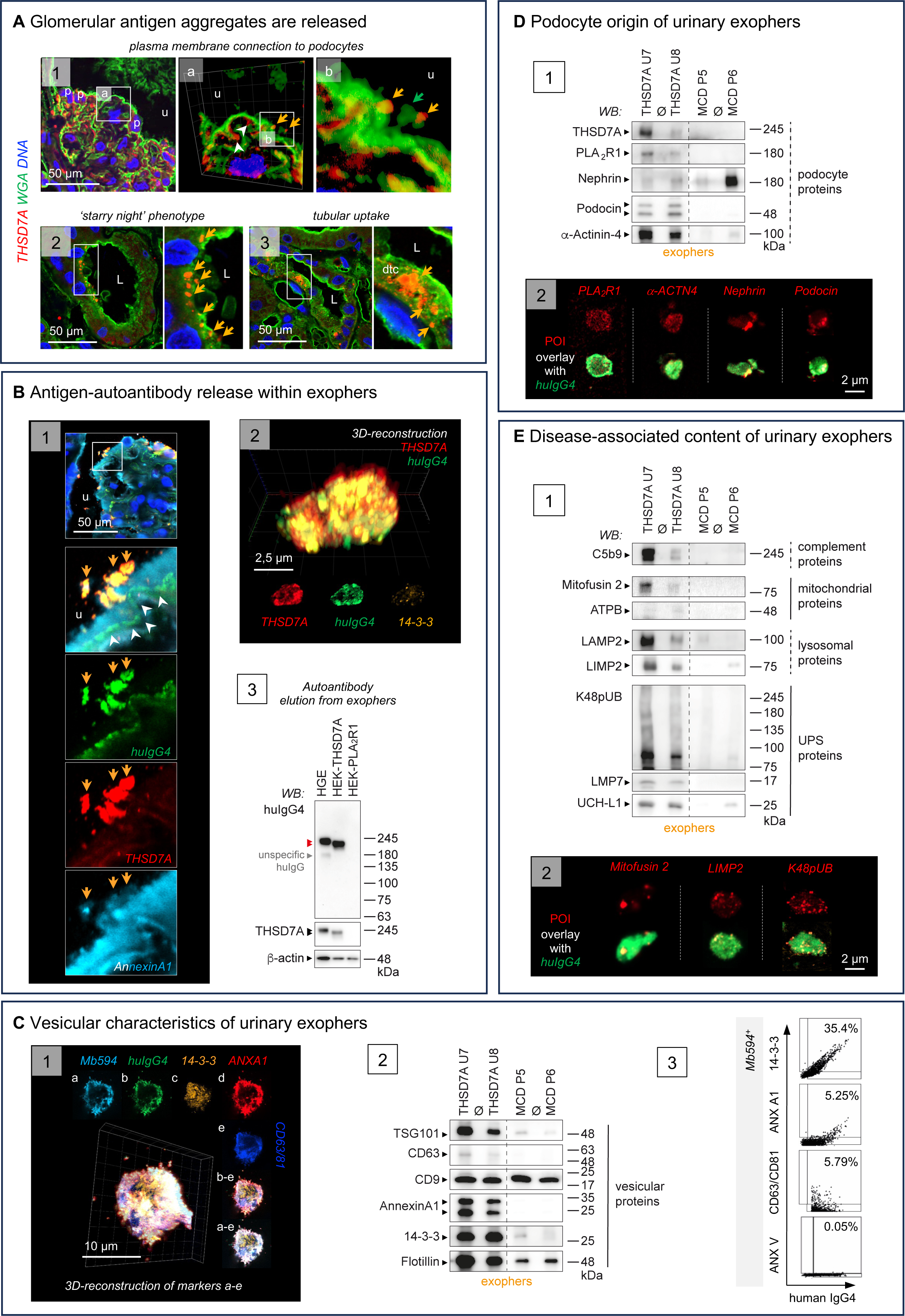
Glomerular urinary space aggregates are exophers in a THSD7A^+^-MN patient. The diagnostic biopsy and urines (U1-U8) collected from the THSD7A^+^-MN patient **P1** were evaluated for the presence of exophers. **A**) High-resolution confocal micrographs to THSD7A (red) and wheat germ agglutinin (green, WGA, binds to N-acetyl-D-glucosamine and sialic acid of the glycocalyx and demarcates plasma membrane), DNA (blue). *Panel 1:* Glomerular THSD7A aggregates localize to the subepithelial space (white arrowheads) and to the urinary (u) space (orange arrows). Urinary space THSD7A aggregates exhibit a WGA^+^ membrane connection (green arrow) to podocytes (p). *Panel 2:* THSD7A^+^ aggregates (orange arrows) are found within the tubular lumen (L) adherent to the brush border or *Panel 3* taken up by tubular cells, dtc = distal tubular cell. **B**) Glomerular and urinary THSD7A aggregates are huIgG4^+^. *Panel 1:* Confocal analyses demonstrate that THSD7A (red) colocalizes with huIgG4 (green) in urinary space aggregates (orange arrows). White arrowheads highlight huIgG4 in the subepithelial space. *Panel 2:* 3D-reconstructed confocal *z*-stack of a urinary exopher (from U1) stained for THSD7A (red), huIgG4 (green) and the vesicle marker 14-3-3 (orange). Lower row exhibits individual channels in one plane. *Panel 3:* Antibodies eluted from U2 exophers exhibit specific reactivity to THSD7A in human glomerular extract (HGE) and in HEK cells transduced with human THSD7A but not with human PLA_2_R1. **C**) Urinary exophers express vesicle markers. *Panel 1:* 3D-reconstructed confocal *z*-stack of an exopher from U2 demonstrates expression of the vesicle markers 14-3-3 (orange), Annexin A1 (red), CD63/CD81 (blue). MemBrite594 (light blue) stains the plasma membrane, huIgG4 (green) the bound autoantibody. *a-e:* individual channels in one plane. *Panel 2:* Immunoblot to vesicle markers in exophers isolated from U7 and U8 in comparison to two nephrotic minimal change disease (MCD) patients **P5** and **P6**. *Panel 3:* Image stream quantification of vesicle marker distribution on MemBrite594^+^huIgG4^+^ exophers in U1 discerns 14-3-3 as an abundant marker. Note the lack of Annexin V signal (marker for apoptotic bodies). Plot also shown in Fig. 5B *panel 1* Urinary exophers contain podocyte proteins. *Panel 1:* Immunoblot detection of podocyte-specific proteins in exophers from U7 and U8 in comparison to the two nephrotic MCD patients. *Panel 2:* Confocal analyses demonstrate expression of podocyte-specific proteins (protein of interest (POI) in red) within huIgG4^+^ (green) exophers of U2. **E**) Urinary exophers contain disease associated proteins. *Panel 1:* Immunoblot detection of disease associated proteins belonging to the complement cascade (C5b9, membrane attack complex), mitochondria (membrane proteins Mitofusin 2 and ATP synthase ATPB), lysosomes (membrane proteins LAMP2 and LIMP2), and the ubiquitin proteasome system (UPS: K48-polyubiquitinated proteins, the proteolytic β-subunit LMP7, the deubiquitinating enzyme UCH-L1) in exophers isolated from U7 and U8 in comparison to the two nephrotic MCD patients. *Panel 2:* Confocal analyses demonstrate expression of Mitofusin 2, LIMP2, or K48pUB (all proteins of interest (POI) in red) within huIgG4^+^ (green) exophers of U2.

We discovered THSD7A- or PLA_2_R1-antigen aggregates within the glomerular urinary space that had a membrane connection to podocytes by confocal microscopy of the diagnostic MN-patient biopsies (***Fig. 1A*** *panel 1, **Fig. S1A***). After urinary release, these antigen aggregates were found in adherence to the tubular brush border (***Fig. 1A*** *panel 2, **Fig. S1B***), reminiscent of the ‘starry night’ phenotype of burst *C. elegans* exophers^18^. Additionally, these antigen aggregates localized to the cytoplasm of parietal epithelial and tubular cells, demonstrating cellular reuptake of aggregates released from the glomerulus (***Fig. 1A*** *panel 3, **Figs. S1C*** and ***S2A***). Antigen aggregates were strongly positive for huIgG4, identifying them as immune complexes within the glomerular urinary space and tubular lumen (***Fig. 1B*** and ***Fig. S1D*** *panels 1*, ***Fig. S3***). Strikingly, antigens and huIgG4 colocalized stronger in aggregates within the glomerular urinary space than within the podocyte subepithelial space, as observed by confocal microscopy. Demonstrating the urinary release of these MN-antigen^+^/huIgG4^+^ aggregates, a subset of large (∼5-8 µm) extracellular vesicles (EVs) was discerned among urinary EVs of MN patients by confocal microscopy, in which the MN-antigen and huIgG4 strongly colocalized. These EVs also showed a granular positivity for the EV marker 14-3-3 (***Fig. 1B*** and ***Fig. S1D*** *panels 2*). The mean size of huIgG4^+^-EVs ranged from ∼580 nm in the THSD7A^+^-MN patient (***Fig. S2B***) and ∼450 nm in a PLA_2_R1^+^-MN patient when measured by nanoparticle tracking analyses (NTA, ***Fig. S1E*** *panel 1*), discerning them as a very large urinary EV subset. Characterization of the EV-bound huIgG4 antibodies demonstrated that they exhibited a specific reactivity to human THSD7A or PLA_2_R1 protein upon elution, establishing them as the pathogenic MN-autoantibodies that were bound to the urinary EVs (***Fig. 1B*** and ***Fig. S1D*** *panels 3*). Demonstrating the vesicular origin of these MN-antigen^+^/autoantibody^+^ EVs, immunofluorescence, immunoblotting, and image stream analyses showed the expression of EV marker proteins, such as 14-3-3, Annexin A1, CD63/CD81 (***Fig. 1C*** and ***Fig. S1E***). To this end, 14-3-3 represented the most discerning marker in huIgG4^+^-EVs. This is of interest, as 14-3-3 protein was recently shown to be required for exopher-formation in *C. elegans* neurons^20^. In comparison to nephrotic non-MN patients, huIgG4^+^-EVs isolated from MN-patient urines contained abundant podocyte proteins such as THSD7A, PLA_2_R1, Nephrin, Podocin, and α-Actinin-4 by immunoblot or by immunofluorescence (***Fig. 1D*** and ***Fig. S1F***) showing their podocyte origin. Besides podocyte proteins, disease-associated proteins such as complement (C1q, C5b9), mitochondrial (Mitofusin 2, ATPB), lysosomal (LIMP2, LAMP2), and ubiquitin proteasome system (UPS) proteins (K48-polyubiquitinated proteins as proteasome substrates, UCH-L1 and the immunoproteasome subunit LMP7) were abundant by immunoblot or by immunofluorescence in huIgG4^+^-EVs from MN-patient urines (***Fig. 1E*** and ***Fig. S4***). Showing the glomerular origin of these discovered urinary huIgG4^+^-EVs, confocal analyses localized selected disease-associated proteins to huIgG4^+^ aggregates within the glomerular urinary space (***Figs. S5*** and ***S6***) as well as within the tubular lumen (***Figs. S7*** and ***S8***) within the diagnostic biopsies of THSD7A^+^- and PLA_2_R1^+^-MN patients. Together, these analyses show the formation of exophers in MN glomeruli that contain the MN antigen with bound autoantibody, 14-3-3, and disease-associated proteins, which are released by the podocyte and passage through the nephron for cellular reuptake or urinary removal.

The urinary exophers identified in MN-patients were unique in respect to their appearance and content compared to urinary huIgG4^+^-EVs from nephrotic non-MN patients as summarized in ***Table S3***. HuIgG4^+^-EVs from nephrotic non-MN patients were mostly devoid or low abundant of podocyte proteins, except for huIgG4^+^-EVs from MCD patients which were abundant in Nephrin. Additionally, huIgG4^+^-EVs from nephrotic non-MN patients had a lower content of disease-associated proteins such as complement, proteasome substrates, mitochondria, and lysosomes compared to MN-patient exophers (***Fig 1*** and ***Fig***. ***S1***). Further, they were smaller and had a less aggregate-like huIgG4 expression pattern by immunofluorescence compared to MN exophers (***Figs***. ***S9*** and ***S10*** *panels 1, 2*). Interestingly, the urinary abundance of huIgG4^+^/14-3-3^+^-EVs strongly differed between patient groups. Primary FSGS and tubulo-toxic kidney injury patients exhibited a low abundance (∼0.6-1.2%), whereas MCD and IgAN patients exhibited a high abundance of huIgG4^+^/14-3-3^+^-EVs ranging from ∼25% in MCD and ∼10% in the IgAN patient, irrespectively of the extent of proteinuria (***Figs***. ***S9*** and ***S10*** *panels 3*). No specific reactivity to glomerular/podocyte/HEK cell proteins could be identified from the immunoglobulins eluted from isolated huIgG4^+^-EVs of non-MN patients (***Figs***. ***S9*** and ***S10*** *panels 4*).

In summary, these findings identify glomerular urinary space aggregates in THSD7A^+^-and PLA_2_R1^+^-MN patients as exopher vesicles formed by podocytes that contain the MN-antigen and bound autoantibody together with disease-associated proteins. These exophers are accessible to non-invasive molecular analyses through their urinary release as huIgG4^+^/14-3-3^+^-EVs. In the following we turned to experimental models of MN, to establish the 1) subcellular origin and dynamics of podocyte exopher-formation and release in response to MN-autoantibody binding and 2) to determine the clinical effect of exopher-formation and release on MN.

### Exophers form at podocyte processes and cell body in experimental THSD7A^+^-MN

In mice, THSD7A^+^-MN was induced by intravenous injection of rabbit (rb)THSD7A-antibodies (abs)^31^. Scanning electron microscopy of day 1, 3, and 7 kidneys (***Fig 2A***, ***Fig. S11***) revealed that podocytes formed long membrane extensions with distal bulges that extended into the urinary space in response to rbTHSD7A-ab exposure. These structures were similar to exophers by morphology^18^ and increased in length and abundance over time. The membrane extensions originated from the urinary side of foot and major processes as well as from the cell body (***Fig. 2A*** *panel 1*, ***Figs. S11*** and ***S12A***). Additionally, large protrusions with rough surfaces without a membrane extension were focally observed at the urinary side of podocyte processes and cell body (***Figs. S11*** and ***S12A***). The immunogold detection of the bound rbTHSD7A-abs on day 7 abundantly localized rbIgG to the distal ends of the long membrane extensions as large, gold-labeled mulberry-like structures within the glomerular urinary space by transmission EM (***Fig. 2A*** *panel 2* and *3, **Fig. S12B***-***D***). Additionally, extensive gold-labeled aggregates were observed that directly protruded from the urinary side plasma membranes of podocyte foot and major processes. In comparison to the abundant gold-labeling of urinary space aggregates, the ‘expected’ subepithelial deposition of rbTHSD7A-abs was discreet on day 7 (***Fig. 2A*** *panel 3, **Figs. S12C*** and ***S13A***). In contrast, subepithelial deposition of rbTHSD7A-abs was abundant on day 1, whereas gold-labeling of podocyte processes within the glomerular urinary space was scarce (***Fig. S13B***). No intracellular gold-particles were observed at the evaluated time points. Immunogold labeling of THSD7A as the corresponding antigen also discerned gold-labeled THSD7A within and at the distal end of podocyte-derived membrane extensions that reached into the urinary space (***Fig. 2A*** *panel 4*). Further, gold labeled THSD7A was present within aggregates protruding from podocyte foot processes (***Fig. S12E***, ***F***). This pattern was comparable to the rbIgG immunogold aggregate pattern described above. Like in MN-patients, THSD7A^+^-MN mice released glomerular urinary space aggregates as exophers to the urine. In line, vesicular structures with gold-labeled rbTHSD7A-abs were observed contacting proximal tubular brush borders (***Fig. S14A***). By confocal microscopy THSD7A^+^/rbIgG^+^ complexes were seen within proximal tubular and parietal epithelial cells (***Fig. S14B***) showing their reuptake after having been released from podocytes within the glomerulus.

**Figure 2:**
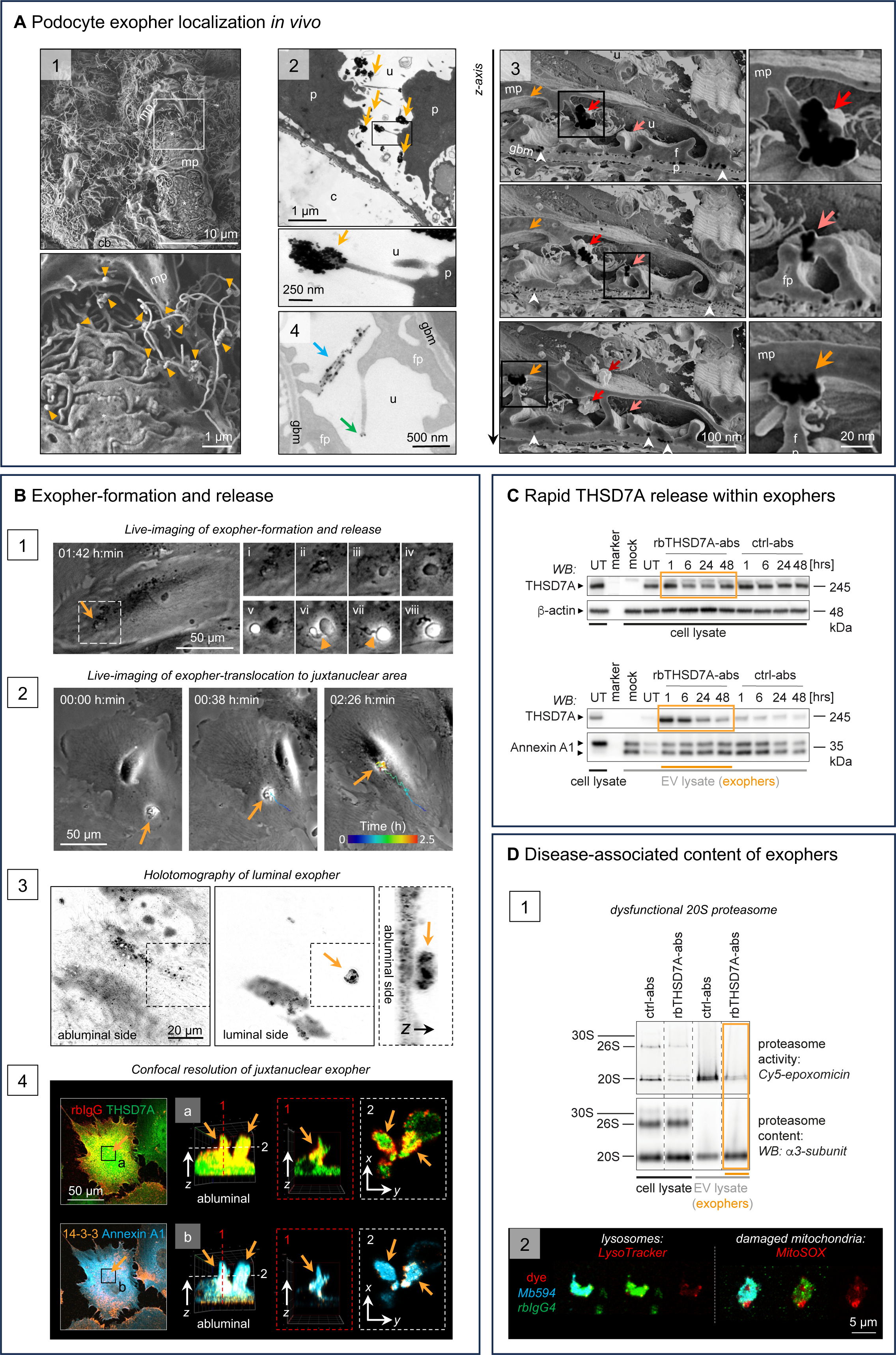
Ultrastructure and dynamics of exopher-formation in experimental MN. **A)** THSD7A^+^-MN was induced in BALB/c mice by injection of rabbit (rb)THSD7A-antibodies (abs) or control-abs. Exopher-formation is visualized by ultra-structural techniques on day 7 in THSD7A^+^-MN mice. *Panel 1*: Scanning EM (SEM) depicting long membrane extensions with distal bulges (orange arrowheads) which originate from major (mp) and foot processes (fp) as well as from the cell body (cb) of podocytes; asterisks = area of fp effacement. *Panel 2*: Transmission EM (TEM) of immunogold-labeled rbTHSD7A-abs demonstrates gold aggregates within the urinary space connected to the podocyte cell body by long membrane extensions (orange arrows); u = urinary space, c = capillary lumen, gbm = glomerular basement membrane, p = podocyte. Close-up: exopher. *Panel 3*: 3D-reconstruction of 48 consecutive TEM micrographs demonstrates the budding-off of gold-labeled structures from podocyte foot and major processes; colored arrows highlight one individual gold structure; gold appears black in the plane of cutting; white arrowheads demarcate subepithelial gold. *Panel 4*: TEM of immunogold-labeled THSD7A depicts the antigen along (blue arrow) and at the distal end (green arrow) of membrane extensions. **B**) Live-cell imaging of human podocytes overexpressing human THSD7A that were exposed to rbTHSD7A-abs in culture. *Panel 1:* Phase contrast live-cell imaging of a forming exopher (see ***Movie S1***). The inset panels correspond to the emboxed area. Images at different time points are shown (i= 01:42 h:min, ii= 01:54, iii= 02:15, iv= 02:22, v= 02:28, vi= 02:32, vii= 02:33, viii= 02:34). Orange arrowhead highlights exopher stalk. *Panel 2:* Phase contrast snapshots at indicated times demonstrate exopher translocation from the cell periphery to the juxtanuclear cell area. The exophers trajectory is shown color coded over 2 h 26 min (see ***Movie S2***). *Panel 3:* Holotomographic images shown with an inverted LUT in grayscale visualize an exopher with a high refractive index protruding from the luminal podocyte side into the medium. The dashed square shows the abluminal (left micrograph) and luminal (middle micrograph) side of the podocyte area, of which the 3D-reconstructed *z-*stack is shown in the right micrograph. Note the inconspicuous abluminal side. *Panel 4:* Confocal resolution of rbIgG (red), THSD7A (green), 14-3-3 (orange), and Annexin A1 (light blue) expression in a podocyte after *in vitro* exposure to rbTHSD7A-abs. For clarity, color overlays are shown pair wise. *Panels a* and *b* represent 3D-reconstructions of the *z-*stack showing exophers that arise from the juxtanuclear plasma membrane area. The micrographs in a dashed box represent single planes in the *z*-axis (red dashed box) or *x/y-*axis (white dashed box) of the 3D-reconstructed image. Note the abundance of Annexin A1 and of 14-3-3 aggregates, as well as rbIgG along the outer aspect of THSD7A in the protruding exophers. **C**) Exposure to rbTHSD7A-abs induces a rapid decrease of THSD7A within the podocyte cell lysate and a concomitant increase of THSD7A in released EVs; UT = untreated THSD7A overexpressing podocytes, mock = control cells without THSD7A expression. **D**) Content of EVs enriched from the medium after 6 h of exposure to ctrl-abs or rbTHSD7A-abs. *Panel 1:* Native PAGE resolution of proteasome complex activity in the cell lysate and the corresponding EV fraction using a pan proteasomal cy5-epoxomicin activity-based probe. Proteasome content is assessed by probing for the α3 proteasome subunit. Note the high abundance of inactive 20S proteasome in exophers. *Panel 2*: Immunofluorescent resolution of lysosomes using LysoTracker or of dysfunctional mitochondria using MitoSOX in exophers (rbIgG^+^Mb594^+^ EVs) released to the medium within 6 hours of rbTHSD7A-ab exposure.

### Exopher-formation encompasses rapid antigen crosslinking, translocation, and release in podocytes

Human podocytes that overexpress THSD7A in culture are specifically targeted by patient^11^ and rbTHSD7A-abs (***Figs. S15*** and ***S16A***). Morphologically, *in vitro* exposure of podocytes to patient THSD7A-abs rapidly induced the formation of honeycomb-like autoantibody patches at the luminal cell surface, which predominated at the cell periphery (***Fig. S15***), indicating antigen crosslinking by the autoantibody. In addition, live-cell imaging discerned areas of plasma membrane alterations at the cell periphery. These areas were the starting point of exopher-formation after exposure to rbTHSD7A-abs (***Fig. 2B*** *panel 1, **Movie S1***). As such, we could observe the translocation of peripheral altered plasma membrane areas towards juxtanuclear cell areas. These translocation processes were accompanied by the formation of stalked, large EVs, which were subsequently released to the medium (***Fig. 2B*** *panels 1* and *2, **Movies S1*** and ***S2***). The abluminal side of podocytes showed no alterations in holotomographic and confocal 3D reconstructions (***Fig. 2B*** *panels 3, 4*). From the luminal side of podocytes, protruding EVs could be seen, which had a high refractive index as observed using holotomography (***Fig. 2B*** *panel 3*). Confocal resolution demonstrated that within these protruding EVs, rbTHSD7A-abs localized to the outer aspect of THSD7A. Further, the EV markers Annexin A1 and 14-3-3 were highly abundant, 14-3-3 in an aggregate pattern (***Fig. 2B*** *panel 4*). Protruding EVs were seen at peripheral and juxtanuclear plasma membrane areas. Together, this identifies the protrusions as forming exophers in cultured podocytes.

The majority of exophers were rapidly released from podocytes within 6 hours of rbTHSD7A-ab exposure and were rbIgG^+^ by image stream (***Fig. S16B***). Released exophers were large (mean ∼470 nm, max ∼4,5 µm) by NTA measurement (***Fig. S16C***). Molecular analyses demonstrated that already within 1 hour of *in vitro* exposure to rbTHSD7A-abs, podocytes released substantial amounts of THSD7A within exophers to the medium, which matched the loss of THSD7A within the corresponding podocyte lysates (***Fig. 2C***). Exophers did not only reduce the podocyte membrane burden of THSD7A and bound autoantibody but also substantiated a cellular exit route for disease-associated proteotoxic material. As such, they were abundant of 20S proteasome complexes with impaired proteolytic activity (***Fig. 2D*** *panel 1*). Additionally, selected exophers exhibited an uptake of LysoTracker and MitoSOX dyes (***Fig. 2D*** *panel 2*), demonstrating the release of lysosomes and damaged mitochondria, respectively.

Together these data show that autoantibody binding to THSD7A initiates crosslinking of the antigen at the podocytes’ peripheral membranes. Altered plasma membrane areas translocate to juxtanuclear membrane areas to be rapidly released as stalked, large exophers. As part of the proteotoxic disease process, exophers contain THSD7A with the bound autoantibody, lysosomes, dysfunctional proteasomes, and damaged mitochondria.

### Urinary exophers are derived from podocytes in experimental THSD7A^+^-MN

Within the urine of THSD7A^+^-MN mice the mean size of EVs was ∼470 nm and as such significantly larger than the mean of ∼390 nm in ctrl-abs injected mice (***Fig. 3A*** *panel 1*). Similarly to MN patients, a subset of urinary EVs were identified as exophers in THSD7A^+^-MN mice using the markers 14-3-3 and rbIgG by image stream (***Fig. 3A*** *panel 2*) and by immunofluorescence (***Fig. 3A*** *panel 3*). We proceeded to establish the podocyte restricted nature and dynamics of urinary exopher release in THSD7A^+^-MN using *mT/mG* reporter mice, which express membrane bound GFP (mGFP) in podocytes and membrane bound tdTomato (mTomato) in all other cells (***Fig. 3B*** *panel 1* and *2*). Using this setup, the urinary release of podocyte (GFP^+^) EVs and of non-podocyte (tdTomato^+^) EVs could be differentially tracked. As exemplarily shown for day 10 urines, a subfraction of podocyte (GFP^+^) EVs was rbIgG^+^ and Annexin V^-^ in THSD7A^+^-MN mice (***Fig. 3B*** *panels 3* and *4*), the latter as expected for exophers^18^. Time course analyses demonstrated that podocytes increasingly released EVs to the urine especially after day 5 following exposure to rbTHSD7A-abs in comparison to ctrl-abs (***Fig. 3C*** *panels 1* and *2*). Starting day 3, a small rbIgG^+^ EV fraction within the podocyte EVs was discernible, which significantly differed in abundance between THSD7A^+^-MN and control urines after day 7 (***Fig. 3C*** *panel 1*). The release of non-podocyte-EVs was not affected by rbTHSD7A-abs or ctrl-abs (***Fig. 3C*** *panels 3* and *4*). Together, these data substantiate that urinary exopher-release is a reaction specific to the podocyte in experimental MN.

**Figure 3:**
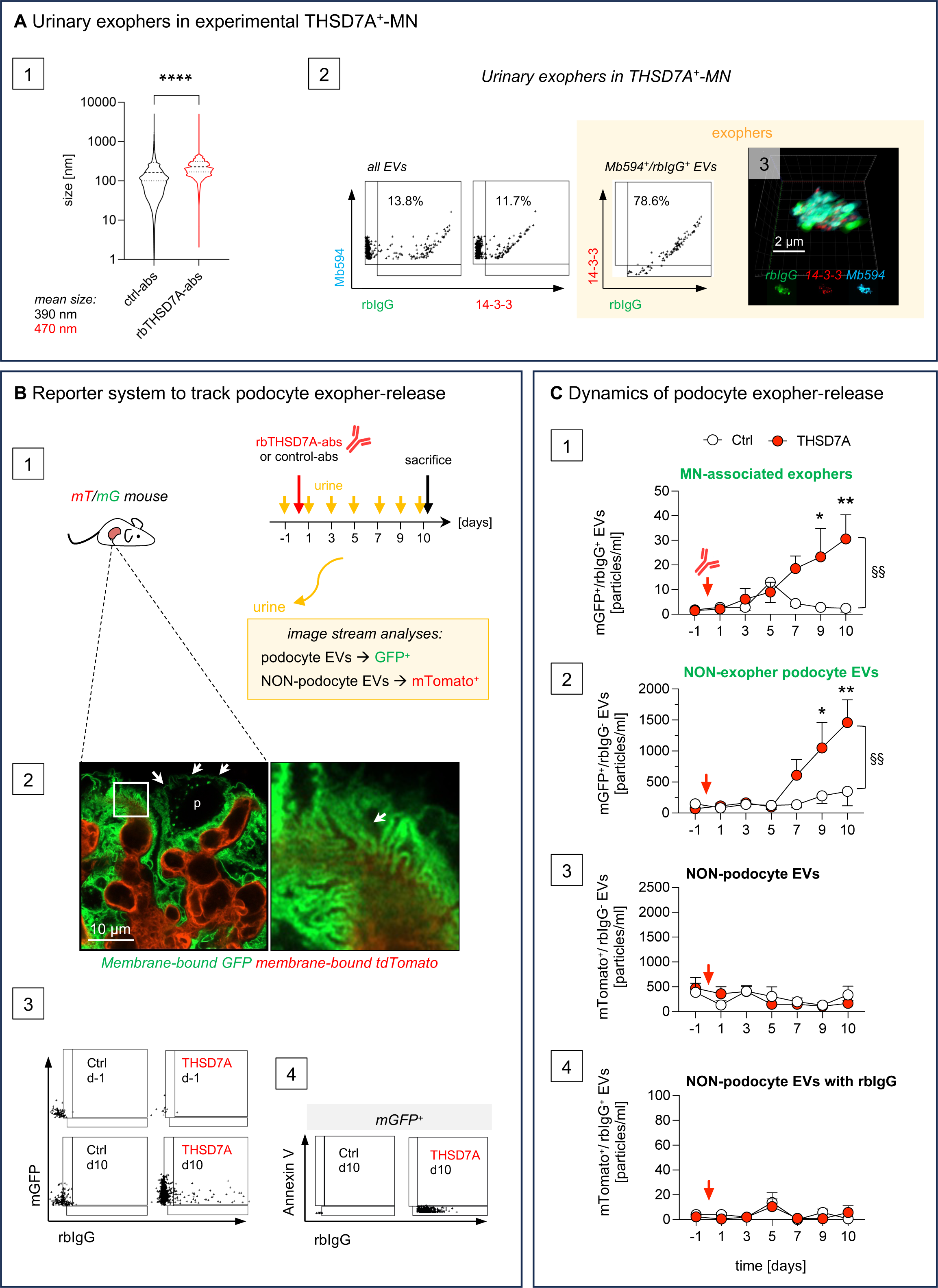
Dynamics of urinary exopher-release in experimental THSD7A^+^-MN. THSD7A^+^-MN was induced in mice by injection of rbTHSD7A-abs or ctrl-abs and urinary EVs were analyzed. **A**) *Panel 1*: Size distribution of urinary EVs assessed by NTA. Violin plots indicate median, 25% and 75% percentile; \*\*\*\**p* <0.0001, unpaired t-test, pooled data of day 7 and 9 urine measurements of *N* = 3 mice per group. Mean size of EVs is indicated. The presence of exophers was assessed *panel 2* by image stream and panel 3 by confocal microscopy using MemBrite594 (light blue) to stain the plasma membrane, rbIgG^+^ (green) to detect bound autoantibodies, and 14-3-3 (red) as vesicular marker that is abundant in exophers. Micrograph depicts 3D-reconstructed confocal *z*-stack of an exopher, individual channels of one plane are shown in the bottom row micrographs. **B**) *Panel 1* illustrates experimental setup in *mT/mG* reporter mice, which express membrane bound GFP (mGFP) in podocytes and membrane bound tdTomato (mTomato) in all other cells. Urinary EVs were isolated from 250 µl urine and 10^9^ total EVs were analyzed by image stream. EVs released from podocytes were discerned by their GFP fluorescence, EVs released from non-podocyte cells by tdTomato fluorescence by image stream. *Panel 2*: Kidney of a naïve *mT/mG* mouse was cleared with SCALE VIEW and the intrinsic fluorescence of the reporter proteins was assessed. Note the clear demarcation of the podocyte (p) plasma membrane with mGFP at the cell body and foot processes (arrows). *Panel 3*: Representative day (d) −1 and day 10 image stream plots of GFP^+^/rbIgG^+^ urinary EVs, which represent the MN-associated exophers. *Panel 4*: Representative day 9 image stream plots of % Annexin V^+^ EVs from the mGFP^+^/rbIgG^+^ EV population (MN-associated exophers) to demarcate apoptotic bodies. **C**) Graphs depict quantification of the 4 different EV subtypes released to the urine in the course of disease development, mean ± SEM, *N* = 11 control-abs, *N* = 14 rbTHSD7A-abs, pooled data from 2 independent experiments; \**p ≤* 0.05, \*\**p ≤* 0.01 to ctrl mice, ^§§^*p ≤* 0.01 between ctrl-ab and rbTHSD7A-ab time course, 2-Way ANOVA with Tukey’s multiple comparison test. Red arrow indicates time of rbTHSD7A-ab injection.

### Balanced podocyte exopher-formation and release is protective in experimental THSD7A^+^-MN

We assessed whether modulation of exopher-formation and urinary release affected clinical and morphological THSD7A^+^-MN. For this purpose, the extent of albumin loss to the urine was monitored in parallel to the release of exophers in the development of THSD7A^+^-MN. With beginning albuminuria 1 day after MN induction, urinary release of exophers was discernible. Following an initial increase, exopher-release decreased in the course of disease development, whereas albuminuria continued to worsen (***Fig. 4A*** *panel 1*). Acceleration of exopher-release through ‘short-term’ proteotoxic stress (induced by proteasome inhibition on day 7) was mirrored by a momentary decrease of albuminuria relative to vehicle treated THSD7A^+^-MN mice (***Fig. 4A*** *panel 2*) suggesting a protective effect. Podocyte exophers were the only urinary EVs whose release was increased by proteotoxic stress (***Fig. S17A, B***). Strikingly, exposure to ‘long-term’ proteotoxic stress (achieved by starting proteasome inhibition prior to THSD7A^+^-MN induction and by maintaining proteotoxic stress throughout the development of MN) exacerbated urinary exopher-release especially between days 5 to 7 relative to vehicle-THSD7A^+^-MN mice (***Fig. 4A*** *panel 3*). Thereafter, urinary exopher-release diminished relative to vehicle-THSD7A^+^-MN, despite the persistence of circulating serum autoantibodies (***Fig. S18***). This late decrease in exopher-release was mirrored by a strong exacerbation of albuminuria relative to vehicle controls. Morphologically, proteasome-inhibited THSD7A^+^-MN mice displayed larger glomerular aggregates in the urinary space than vehicle treated THSD7A^+^-MN mice, especially in the setting of prolonged proteotoxic stress when visualized by super resolution microscopy of rbIgG aggregates in optically cleared kidney sections from *mT/mG* mice (***Fig. 4A*** *panel 4* and *5*). Identifying them as exophers, these rbIgG^+^ aggregates were in connection with the podocyte cell body through long mGFP^+^ membrane extensions (***Fig. 4A*** *panel 4*). These data suggest that especially in the setting of prolonged proteotoxic stress the decrease in urinary exopher abundance at the end of the observation time resulted from impaired exopher-release rather than from impaired exopher-formation. We quantified the amount of THSD7A and rbIgG protein deposition within a defined number of glomeruli at the end of the observation period by immunoblotting (***Fig. S17C***) to assess whether exopher-release affected the extent of glomerular antigen / autoantibody burden. Indeed, correlative analyses demonstrated a significant inverse dependency between the extent of urinary exopher-release and the glomerular deposition of THSD7A protein (***Fig. 4B*** *panel 1*) or of rbIgG in vehicle and ‘short-term’ proteotoxic stress exposed mice (***Fig. 4B*** *panel 2*). Additionally, urinary exopher abundance exhibited an inverse correlation with albuminuria (***Fig. 4B*** *panel 3*), demonstrating that exopher-release is protective in MN by reducing the glomerular burden of antigen/autoantibody deposition.

**Figure 4:**
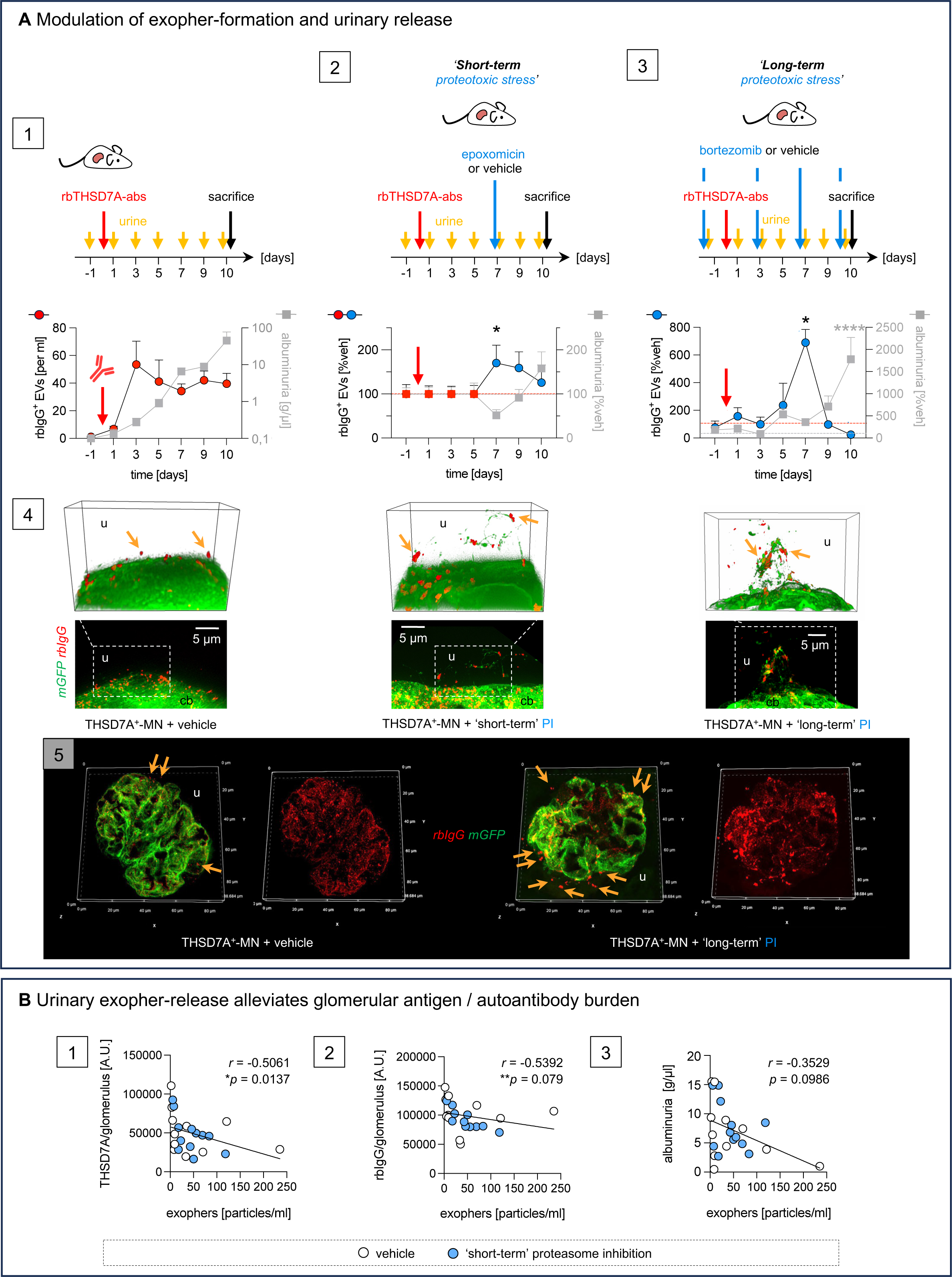
Modulation of exopher-release in experimental THSD7A^+^-MN. **A)***Panel 1*: THSD7A^+^-MN was induced in *mT/mG* reporter and BALB/c mice on day 0 by administration of rbTHSD7A-abs (red arrow), urine was collected in the time course, on day 10 kidney slices and glomeruli were isolated. Abundance of rbIgG^+^-EVs was assessed from 10^9^ total urinary EVs by image stream. Albuminuria was measured in parallel. Graph depicts the time course of urinary exopher release (rbIgG^+^-EVs; particles/ml; left *y* axis, red round symbols) and albuminuria (right *y* axis, grey square symbols; g/µl) in THSD7A^+^-MN. *Panel 2*: Additionally, short-term proteotoxic stress was achieved by administration of a proteasomal inhibitor (epoxomicin, blue arrow) or vehicle (DMSO) every day after day 7 for 4 consecutive days. Graph depicts changes in urinary exopher-release (left *y* axis, colored round symbols) and albuminuria (right *y* axis, grey square symbols) in short-term proteasome inhibited mice relative to vehicle-THSD7A^+^-MN mice (dashed lines at 100%). Pooled values from 3 independent experiments; *N* > 9 per group, mean ± SEM; \**p* < 0.05, 2-way ANOVA with Tukey’s multiple comparison test. *Panel 3*: Long-term proteotoxic stress was achieved by administration of the proteasomal inhibitor bortezomib or vehicle prior to MN induction on day −1 followed by 2 times per week for 10 days. Graph depicts changes in urinary exopher-release (left *y* axis, blue round symbols) and albuminuria (right *y* axis, grey square symbols) in long-term proteasome inhibited relative to vehicle THSD7A^+^-MN mice (dashed lines at 100%); values from 1 experiment; N > 4 per group, mean ± SEM, \*\*\*\**p* < 0.0001, 2-way ANOVA with Tukey’s multiple comparison test. *Panel 4* and *5:* Kidney slices of *mT/mG* reporter mice that express membrane (m) bound GFP (mGFP, green) in podocytes were optically cleared and stained for bound rbTHSD7A-abs (red). In *panel 4* 3D-reconstructed z-stacks of the podocyte cell body (cb) plasma membrane is shown, in *panel 5* a 3D-reconstructed overview of a glomerulus; u = glomerular urinary space, orange arrows highlight individual exophers. **B**) Spearman’s correlations of total glomerular *panel 1* THSD7A or *panel 2* rbIgG protein abundance (as detailed in ***Fig. S17C***) and *panel 3* of albuminuria (g/µl) to the respective urinary exopher abundance (particles/ml) at time of sacrifice.

### Non-invasive exopher monitoring in a patient with relapsing THSD7A^+^-MN

Urinary exopher-release and as such the podocytes’ ability to remove bound autoantibodies was prospectively monitored for 2½ years in a 62-year-old male who was diagnosed with THSD7A^+^-MN in May 2021. The patient initially presented with heavy nephrotic syndrome and normal renal function. His diagnostic histopathology was significant for a THSD7A^+^-MN (stage I Ehrenreich and Churg), with segmental thickened GBM (***Fig. 5A*** *panel 1*), a pseudolinear positivity for huIgG (***Fig. 5A*** *panel 2*), a fine glomerular positivity for huIgG4 at the filtration barrier (***Fig. 5A*** *panel 3*), and THSD7A aggregates. THSD7A aggregates were discreet in the subepithelial space and prominent in the urinary space, some within parietal epithelial cells (***Fig. 5A*** *panel 4*). Ultra-structurally, foot process effacement and GBM thickening were seen, with some long membrane extensions derived from foot and major processes (***Fig. 5A*** *panel 5*). The clinical course of the patient (***Fig. 5B*** *graph, **Table S1***, *patient P1*) showed a relapsing nephrotic syndrome complicated by cerebral infarctions (09/2021 and 10/2022), and a progressive decline of renal function (eGFR 90 ml/min at diagnosis to 33 ml/min 12/2023). The patient received several courses of rituximab as well as a cyclophosphamide pulse therapy, under which he developed a symptomatic COVID-19 infection. Besides the diagnostic urine (U1), 7 additional urines (U2-U8) were collected and analyzed for exopher abundance (***Fig. 5B*** *panel 1*), for the presence of exopher-bound autoantibodies (***Fig. 5B*** *panel 2*), and for the presence of a molecular imprint of podocyte injury pathways (***Fig. 5B*** *panel 3*). Albeit the presence of high sTHSD7A-ab titers of 1:160, urinary exopher-release progressively decreased from ∼35% at diagnosis to ∼3% in U3. From 10/2022 on, sTHSD7A-abs were mostly negative, nonetheless autoantibodies were persistently eluted from urinary exophers. Four months after the last rituximab administration, the patient presented with increasing edemas and proteinuria (28/06/2023). Upon immunological checkup, a low but recovering CD19^+^ B-cell count was noted (2%; 14/µl), sTHSD7A-abs were still not detectable. Nonetheless, analyses of U7 showed an increase in urinary exopher abundance (4%) which contained bound autoantibodies demonstrating immunologic activity. Rituximab administration ameliorated the patients’ proteinuria and edema. U8 collected 4½ months later still in the setting of negative sTHSD7A-abs, exhibited a low exopher abundance (∼0.2%) with readily detectable exopher-bound autoantibodies. Molecular disease patterning by proteomic analyses of exophers from U1-U8 (***Fig. 5B*** *panel 3*) not only substantiated the vesicular/plasma membrane origin of exophers, but also the enrichment of disease associated pathways such as cell stress response (heat shock proteins, chaperones), classical complement and humoral immune system, as well as proteolytic pathways (proteasome and lysosome) shown within our biochemical analyses. Even though a molecular imprint of the podocyte injury state was visible over all investigated time points, a certain dynamic in the proteome profile was present.

**Figure 5:**
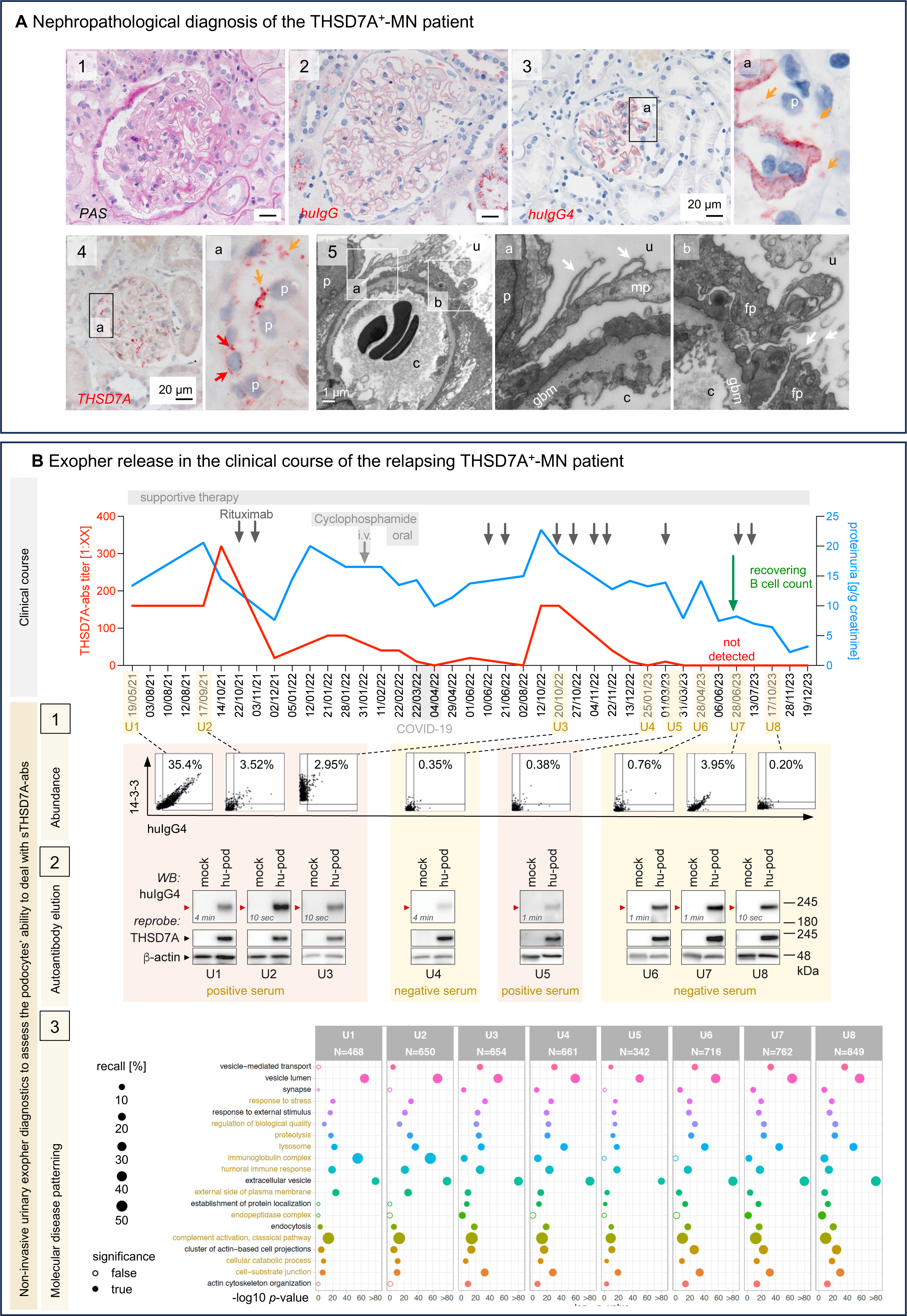
Non-invasive exopher monitoring in relapsing THSD7A^+^-MN. **A)** Micrographs from the nephropathological diagnosis (19/05/2021) of the patient **P1** encompassing light microscopy with *Panel 1*: PAS staining for general morphology; immunohistochemistry to *Panel 2*: huIgG, *Panel 3*: huIgG4, and *Panel 4*: THSD7A. Orange arrows highlight aggregates within the urinary space, red arrows aggregates within parietal epithelial cells. *Panel 5*: Transmission EM; white arrows highlight membrane extensions from major (*panel 5a*) and foot processes (*panel 5b*), p = podocyte, gbm = glomerular basement membrane, fp = foot process, mp = major process, u = urinary space, c = capillary space. **B**) Dynamics and characteristics of urinary exopher-release over 2 ½ years of relapsing nephrotic syndrome. *Graph* depicts on the left *y*-axis the serum THSD7A autoantibody titer (red curve) and on the right *y*-axis urinary protein to creatinine ratio (blue curve). The main clinical interventions, as well as the COVID-19 infection are indicated. *Panel 1*: Image stream plots demonstrate the relative content of huIgG4^+^ exophers in the urine in % exophers from 10^9^ total EVs. *Panel 2:* Reactivity of the huIgG4 eluted from 10^9^ exophers to THSD7A protein; hu-pod = human podocytes with human THSD7A overexpression, mock = human podocytes without THSD7A overexpression. Membranes were reprobed for THSD7A and β-actin to control for THSD7A expression/height and loading. The exposure times for the huIgG4 signal are indicated, all blots originate from one membrane exposed at the same time. Note the clear detection of THSD7A-autoantibodies bound to released exophers in all urines, albeit a concomitant negative sTHSD7A-ab titer. *Panel 3*: Proteomic molecular disease patterning of exophers isolated from U1-U8. For pathway analysis, only proteins identified with ≥ 2 peptides with a Q-value < 0.001 were used. Graph displays the *p*-values corresponding to the significance of pathway enrichment of selected functions. *P*-values below 1E-80 were capped; recall rate = proteins identified and assigned to category / number of proteins that are annotated to the category. Pathways reflecting exopher-specific processes are highlighted in ocher. Detailed information on the pathway proteins is provided in ***Table S4***. A total list of identified proteins can be found on PRIDE.

### Glomerular aggregate morphometrics and non-invasive exopher monitoring in PLA_2_R1^+^-MN

Patients with THSD7A^+^-MN are rare, therefore we developed a computational image algorithm to pattern subepithelial and urinary space PLA_2_R1 aggregates throughout the glomeruli, the latter as the biological correlate of podocyte exopher-formation. The ‘density score’ of this algorithm included the size, amount, neighborhood, and intensity of PLA_2_R1 aggregates throughout the glomeruli not differentiating between urinary space and subepithelial aggregates (***Fig. S19***). A low density score related to the predominance of urinary space aggregates and a disrupted subepithelial PLA_2_R1 deposition pattern, whereas a high density score reflected the occurrence of dense subepithelial PLA_2_R1 depositions and scarce/large urinary space aggregates (***Fig. 6A*** *panel 1*). PLA_2_R1 density differed in diagnostic biopsies of a well-characterized retrospective PLA_2_R1^+^-MN patient cohort^32^ in which 18 patients were stratified to a remission group (R; amelioration of serum creatinine, eGFR, proteinuria, albuminuria, and/or serum albumin) within a mean follow-up period of 42,83 ± 5,52 months, and 17 patients to a non-remission (NR) group, within a mean follow-up period of 43,24 ± 9,2 months. In the remission group, PLA_2_R1 density and sPLA_2_R1-abs were low at diagnosis. The non-remission group exhibited a high PLA_2_R1 density as well as high sPLA_2_R1 titers at diagnosis (***Fig. 6A*** *panel 2*). These data suggest that the amount of antigen deposited within the subepithelial space and/or the ability to clear the subepithelial space from the PLA_2_R1 deposits differed at diagnosis between both patient groups.

**Figure 6:**
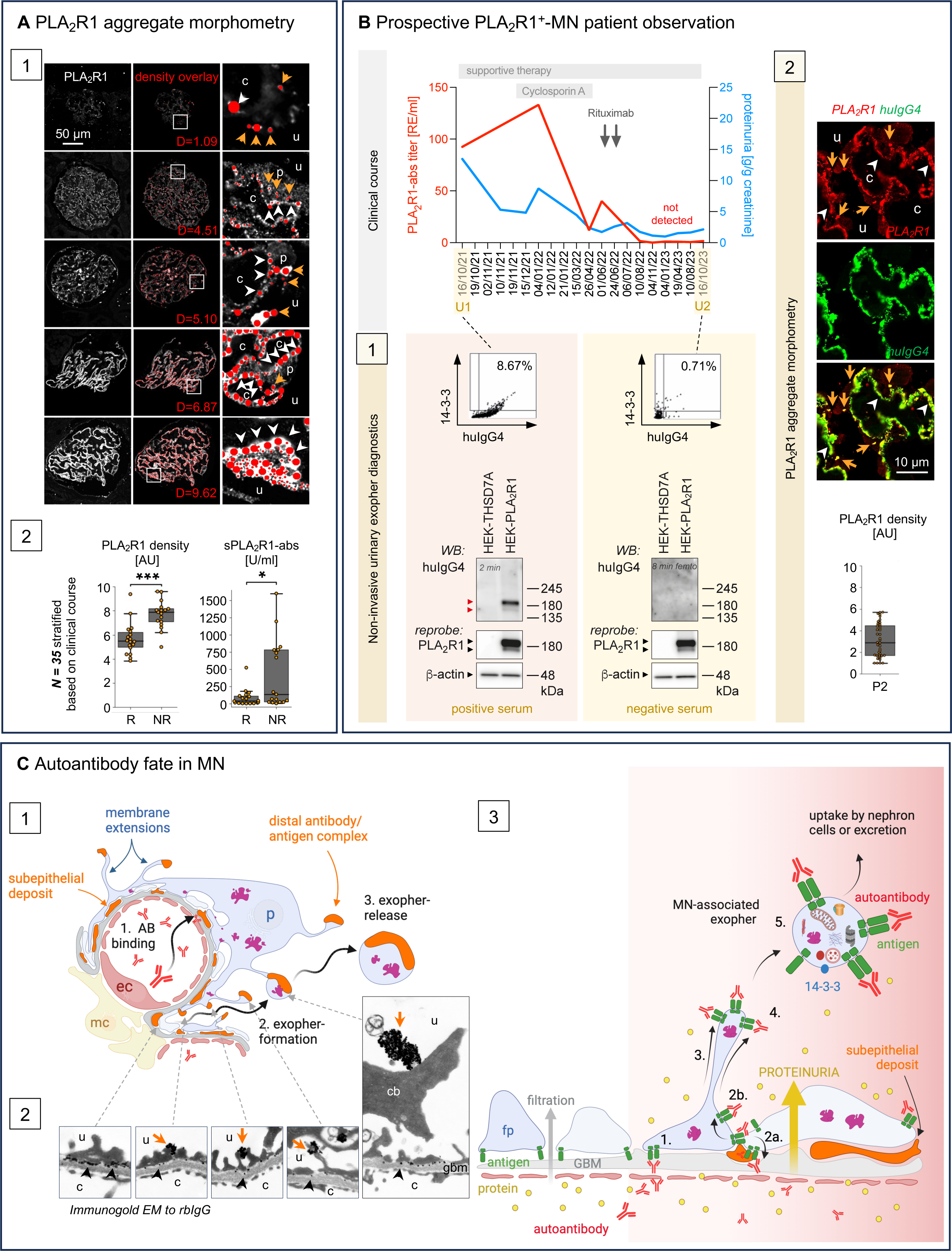
Glomerular aggregate morphometrics and non-invasive exopher monitoring in PLA_2_R1^+^-MN. **A)** Confocal images of PLA_2_R1 stainings (white) from PLA_2_R1^+^-MN patients were subjected to computational image analyses of the glomerular PLA_2_R1 aggregate pattern. *Panel 1*: Scoring range of the PLA_2_R1 aggregate density algorithm (D = density; arbitrary units); red areas in overlay are shown, orange arrows = urinary space PLA_2_R1 aggregates, white arrowheads subepithelial PLA_2_R1 deposition, P = podocyte nucleus, u = urinary space, c = capillary space. *Panel 2:* PLA_2_R1 aggregate density was quantified in diagnostic biopsies of a retrospective PLA_2_R1^+^-MN patient cohort^32^, assembled based on clinical outcome. *N* = 18 patients were stratified into the remission group R and *N* = 17 into the non-remission group NR. Boxplots show the median, first and third quartiles, and whiskers 1.5 times interquartile range of glomerular PLA_2_R1 density (A.U. = arbitrary units) or of the serum PLA_2_R1 autoantibody (sPLA_2_R1-ab) titers in units/ml at diagnosis, \**p* ≤ 0.05, **\*\*\****p* ≤ 0.001, Mann Whitney U-Test. **B**) Prospective analysis of a 22-yr-old female PLA_2_R1^+^-MN patient **P2** diagnosed 16/10/2021. Graph depicts clinical course of the serum PLA_2_R1 autoantibody titer (red curve; left *y*-axis) and of the urinary protein to creatinine ratio (blue curve; right *y*-axis). Diagnostic (U1) and follow-up (U2) urines were collected. The main clinical interventions are indicated. *Panel 1:* Image stream quantification of %huIgG4^+^/14-3-3^+^ exophers present within 10^9^ Mb594^+^ urinary EVs and immunoblot assessment of the specific reactivity of exopher-eluted huIgG4 to huPLA_2_R1 or huTHSD7A protein in HEK cell lysates; reprobes control for PLA_2_R1 expression/height and loading; exposure times for the huIgG4 signal are indicated. Note the clear detection of PLA_2_R1 autoantibodies in U1 and their absence in U2. *Panel 2*: Confocal analyses of PLA_2_R1 (red) and huIgG4 (green) expression pattern in the diagnostic biopsy. Boxplot shows quantification of PLA_2_R1 density, the median, first and third quartiles, and whiskers 1.5 times interquartile range are indicated; each datapoint represents one glomerular section. Note the presence of small exophers in the urinary space (orange arrows) and a disrupted subepithelial PLA_2_R1 pattern (white arrowheads). **C**) Scheme summarizing the current concept of autoantibody fate using THSD7A as model antigen. *Panel 1:* The glomerular autoantibody burden depends on 1. the amount of circulating autoantibody that binds subepithelially to THSD7A and the ability of podocytes to dispose of these autoantibodies by 2. exopher-formation and 3. exopher-release to the urine. *Panels 2*, *3*: Exopher-formation involves autoantibody-mediated THSD7A crosslinking in the subepithelial space (**1.**), deposition of THSD7A/autoantibody complexes in the subepithelial space (**2a**., classical concept of deposit formation; black arrowheads in the immunogold EM micrographs to rbIgG) and/or translocation towards the apical foot process membrane (**2b**., new concept; orange arrows in the immunogold EM micrographs to rbIgG). Apically displaced THSD7A/autoantibody complexes at foot, major processes and podocyte cell body extend to the urinary space through exopher-formation (**3.**). Exophers contain THSD7A and the extracellularly bound autoantibodies in addition to 14-3-3 as discerning EV marker, disease-associated proteins, dysfunctional mitochondria, and degradative machinery (**4.**). Exophers are released to the urine and can be taken up by other nephron cells (**5.**). Scheme created with Biorender; ec = endothelial cell, mc = mesangial cell, p = podocytes, GBM = glomerular basement membrane, fp = foot process, c = capillary lumen, u = urinary space, cb = cell body.

Exopher-release was prospectively analyzed in a 20-year-old female nephrotic patient diagnosed 10/2021 with PLA_2_R1^+^-MN (stage I-III Ehrenreich and Churg) based on the clinical and nephropathological workup (***Fig 6B***, ***Table S1***, *patient P2*). At diagnosis sPLA_2_R1-ab titer was moderate (92 RE/ml). Urinary exopher diagnostics identified ∼8% exophers, and exopher-bound PLA_2_R1 autoantibodies were readily detectable (***Fig 6B*** *panel 1*). Morphologically, PLA_2_R1 density was low in the diagnostic biopsy, which related to a disrupted subepithelial PLA_2_R1 deposition and small glomerular urinary-space PLA_2_R1 aggregates (***Fig 6B*** *panel 2*). Together, these findings indicated a low glomerular aggregate burden in the setting of moderate sPLA_2_R1-ab titer, with dynamic exopher-formation and release at diagnosis. Within one year the patient went into clinical remission under a supportive treatment regimen, and cyclosporin A. A relapse 01/06/2022 was treated with rituximab. Serum PLA_2_R1 titers were first negative 10/08/2022 and remained negative thereafter, suggesting immunologic remission. In the last follow-up urine U2 (16/10/2023) exopher abundance was low (∼0.7%) and no exopher-bound PLA_2_R1 autoantibodies were detectable, substantiating immunologic remission. Based on these patient observations we propose that simultaneously tracking the fate of pathogenic autoantibodies (serum levels, glomerular binding, and release) enables a fine-tuned monitoring of immunologic MN disease activity, especially in the setting of negative serum autoantibody titers.

In conclusion, our cumulative investigations in patient and experimental MN demonstrate that a balanced process of podocyte exopher-formation and release represents a protective antigen-overarching pathophysiologic sequel in MN. As summarized in ***Fig 6C***, we propose that simultaneously tracking the fate of pathogenic autoantibodies (serum levels, glomerular binding, and release, ***Fig. 6C*** *panel 1*) enables a fine-tuned monitoring of immunologic MN disease activity. Mechanistically, exopher-formation includes the displacement of antigen/autoantibodies from the subepithelial space to the apical membranes of foot and major processes and cell body (***Fig. 6C*** *panel 2*). Exopher-release to the urine ultimately leads to a reduction of the glomerular subepithelial autoantibody and antigen burden (***Fig. 6C*** *panel 3*).

## DISCUSSION

Podocytes continue to emerge as primary targets to autoimmunity not only in MN^33^. Our knowledge about how autoantibodies affect podocytes is limited and the mechanisms of the glomerular antigen/autoantibody deposition and clearance are unknown. Here we discover that exophers represent the biological correlate of aggregates within the glomerular urinary space in MN in patients and in experimental MN models. In patients, MN-associated (autoantibody^+^/antigen^+^) EVs fulfilled important features of exophers. As such, *morphological criteria* (extrusion from apical membranes especially from the juxtanuclear area as stalked large vesicles; expression of a characteristic set of EV markers)^18,20,21^, *dynamic criteria* (enhanced release by proteotoxic stress, reuptake by cells along the nephron)^18,27,34^, and *molecular criteria* (abundance of cell-specific and disease-associated proteins, presence of organelles)^18,35,22^ establish them as a hitherto unrecognized podocyte-derived urinary EV form. They are distinct from migrasomes^36^, a large EV-type forming at tips and intersections of retraction fibers of migrating cells^37^. They also differ in size and vesicular marker composition from EVs described by Hara et al., which originate from tip vesiculation of podocyte microvilli^38^. Whether the membrane extensions identified in THSD7A^+^-MN mice reflect the well-known feature of microvillous changes of injured podocytes^39^ is unclear. However, both processes, i) exopher-formation in neurons and cardiomyocytes^22^, as well as ii) microvillous podocyte transformation are considered protective mechanisms, the latter suggested to be associated with complete remission of MCD or FSGS in patients^40^. The protective cellular effects of exopher-formation and release include the removal of toxic components when proteostasis and organelle function are impaired^18^. Podocytes face both challenges in experimental MN^32,41–43^. Additionally, protection can be attributed to clearance of the subepithelial space from immune complexes and complement by their membrane translocation to the urinary podocyte side and release within exophers. As such, the course of albuminuria and the extent of glomerular antigen/autoantibody deposition inversely correlated with exopher-release in experimental THSD7A^+^-MN. Exopher-release might reduce the effects on GBM remodeling, and abolish the disease exacerbating effect of complement activation^14^ thus enabling podocyte recovery.

One could speculate that an aspect of MN progression and therapy resistance could originate from high circulating autoantibody levels that constantly target podocytes, which cannot maintain the necessary level of exopher-formation for removal. As such, urinary exopher-release was low in the relapsing THSD7A^+^-MN patient in the setting of high sTHSD7A-abs, and the histological pattern of scarce urinary space aggregates and dense subepithelial deposition predominated in the non-remitting PLA_2_R1^+^-MN patient group. Of note, extensive exopher-formation might also have disease aggravating aspects i) by transfer of toxic protein aggregates to surrounding cells, promoting pathology^44^, ii) by exopher phagocytosis through surrounding cells as a starting point for (cross-)presentation and immune activation leading to secondary autoantibody formation against intracellular proteins^32,45^ or to epitope-spreading of primary MN autoantibodies^46^, iii) by initiating lipid biosynthesis^27^, which when dysregulated perpetuates podocyte injury^47^, and iv) by substantial loss of plasma membrane required for exopher-formation, necessitating membrane repair. The flux balance between serum autoantibody titer, exopher-formation and release is certainly crucial for the ultimate outcome in MN.

We identified the ubiquitously expressed 14-3-3 chaperone-like protein^48^ as an exopher-identifying protein. 14-3-3 abundance was high and ‘aggregate-like’ in patient and experimental exophers, and 14-3-3 aggregates were observed at juxtanuclear areas of exopher-formation in podocytes, corroborating the recently published involvement of 14-3-3 in neuronal exopher-formation^20^. In podocytes, 14-3-3 is a known synaptopodin interacting protein^49^, playing a role in the downstream function of cyclosporin A^49^.

To this end, MN-associated exophers represent a unique form of huIgG4^+^/14-3-3^+^ EVs in respect to their appearance and molecular composition. It is, however, unclear whether the huIgG4^+^/14-3-3^+^ EVs abundantly present in nephrotic urines from IgAN and MCD patients represent exophers or not, as they differ in size, appearance, and molecular composition. In IgAN the binding of human antibodies to aberrantly glycosylated and mesangially deposited IgA drives disease^50^, and a binding of antibodies to podocytes has not been described. Nonetheless, the isolated huIgG4^+^/14-3-3^+^ EVs contained 14-3-3, mitochondrial, lysosomal and UPS proteins as well as podocyte proteins. Also, the abundant huIgG4^+^/14-3-3^+^ EVs present in MCD urines are intriguing. They contained Nephrin but compared to MN-associated exophers no strict exopher-defining proteotoxic/organellar proteins, and a different abundance of EV markers. However, the aggregate-like pattern of huIgG4 and 14-3-3 expression on a subset of MCD EVs, might relate to EV-bound anti-Nephrin antibodies recently identified in the serum of a subset of MCD patients^33^. But as the IgG subtype of anti-Nephrin antibodies is unclear, and the detection of anti-Nephrin antibodies remains technically challenging, further investigations are underway. Of note, huIgG4 present within nephrotic urines adheres to EV surfaces to some extent^51^. Therefore, larger cohorts will be needed to define an adequate cut-off for exopher abundance.

Following the identification of the first MN antigen in 2009^1^, it took several years, a substantial amount of patients and independent observations to establish the indisputable clinical power of serum autoantibody measurement for diagnosis, evaluation of prognosis, and for clinical monitoring of MN^7^. Identification of exopher-formation as the pathobiology behind glomerular aggregate formation adds a clinically relevant information to patient monitoring. The two prospective MN patient observations suggest that tracking the fate of autoantibodies (1. serum levels, 2. glomerular binding, 3. urinary exopher-release and composition) allows fine-tuned monitoring of MN disease activity, especially in the setting of negative serum autoantibody titers. In the PLA_2_R1^+^-MN patient, the low abundance of exophers and the absence of exopher-bound autoantibodies in the follow-up urine substantiated the immunologic remission reflected by the absence of sPLA_2_R1-abs. In the THSD7A^+^-MN patient, however, detection of ∼4% exophers with bound autoantibodies in the setting of negative sTHSD7A-abs substantiated persistent immunologic activity as the cause of nephrotic syndrome aggravation. Another interesting aspect was the low urinary exopher-release (∼0.35%) in the setting of abundant sTHSD7A-abs in March 2023 which, besides scarring, could indicate impaired exopher-formation/release by podocytes as a possible trigger of disease persistence or even aggravation similar to the ‘long-term’ proteasome-inhibited mice.

Therapeutic modulation of podocyte exopher-formation/release could represent an interesting approach i.e. by proteasome modulation or by targeting renal cell exopher uptake^22^ and the ensuing immune reaction. Existing therapeutic principles in MN might already modulate exopher-formation/release, i.e. cyclosporin A through interference with 14-3-3/synaptopodin signaling^49^ and rituximab through off-target effects on podocyte lipid biology^52^, both aspects described to modify exopher-formation in *C. elegans*. As exophers can be non-invasively enriched in substantial amounts, they open a new avenue of diagnostic approaches in nephrology, reaching from simple quantification and autoantibody detection (potentially beyond MN) to deep molecular disease patterning. Exopher-enrichment from nephrotic urines is technically elaborate, therefore the development of simple urine tests is necessary to enable routine clinical use. In the future this might, however, represent a sensitive and non-invasive method to monitor MN autoantibodies by clinicians and even patients themselves, allowing for an early detection of immunologic activity. As exophers enable the molecular assessment of podocyte injury patterns/extent, future investigations in larger prospective patient cohorts will prove their clinical and prognostic value.

## MATERIALS AND METHODS

Please refer to the supplemental appendix for detailed materials and methods.

## ACKNOWLEDGEMENTS

We are grateful to the staff of the FACS Sorting Core Unit and the UKE imaging Facility (UMIF) under the DFG Research Infrastructure Portal: RI_00489 for excellent technical assistance. We thank the Nikon Center of Excellence at LIV for support. This work was funded by the Deutsche Forschungsgemeinschaft (DFG, German Research Foundation, CRC 1192 project B3, B6 and ME 2108/10-1 to CMS; CRC 1328 (Project A20) to P.J.S.) and by the Human Frontier Science Program (HFSP) RGP0032/2022. The LSM980 Airyscan 2 was funded by the DFG (INST 152/952-1 FUGG to CMS). UW was financially supported by the Else Kröner-Fresenius-Stiftung iPRIME Scholarship (2021_EKPK.10), SS, and KN were financially supported by the Integrated Research Training Groupe of the CRC 1192. UKE, Hamburg. We are grateful to Andrea Hilpert and Anita Hecht in Erlangen for excellent technical assistance in preparing the murine immunogold EM samples. We thank Christian Hentschker, and Manuela Gesell Salazar, for excellent support in the proteome analyses. We thank Johanna Herwig for help in the human podocyte *in vitro* assays.

## AUTHOR CONTRIBUTIONS

KL established and performed all the urinary murine and patient vesicle work. WS and KL performed the animal work comprising characterization of *mT/mG* and BALB/c THSD7A^+^-MN. WS performed the THSD7A glomerular quantification. SF performed the hu-podocyte EV experiments, UW, KN, and SK the THSD7A abundance in hu-podocytes. SF, MRA, PJS performed the phase contrast time-lapse and holotomography experiments. LF performed the immunogold SEM and TEMs of murine glomeruli and patient urinary exophers, SG performed the 3D-reconstruction of TEMs. KS and UV performed proteomic analyses of urinary exophers. DL performed histologic procedures, and generated samples, and micrographs for morphometrics. MB, MZ and SB developed the computational imaging algorithms for rbTHSD7A-abs and PLA_2_R1 morphometry. SZ, JB performed excellent technical assistance. TW provided the images of original patient pathological diagnosis. VB-D imaged exophers in patient biopsies. CC and RT performed the 3D-reconstruction of *mT/mG* cleared kidney slice. TNM collected and provided the urine samples and performed the clinical correlations. CMS conceived the project, designed the experiments, performed stainings and confocal microscopy, analyzed data, supervised the project, and wrote the paper.

## DISCLAIMER

SG did not receive any financial compensation for the 3-dimensional TEM reconstructions.

