## Supplemental appendix for "Podocyte exopher-formation as a novel pathomechanism in membranous nephropathy"

### TABLE OF CONTENT

| | | |
|---|---|---|
| *Authors & Institutions* | | *P2* |
| *Suppl. Methods* | | *P3-13* |
| *Figure S1.* | Exophers are found in nephrotic $PLA_2R1^+$-MN patients P2-P4. | *P14-15* |
| *Figure S2.* | Exophers are found in a THSD7A+-MN patient P1. | *P15* |
| *Figure S3.* | Antigen/autoantibody complexes are shed to the urine and found in adherence to the tubular apical membrane in membranous nephropathy patients P1 and P2. | *P16* |
| *Figure S4.* | Urinary exophers enriched from the diagnostic urines of $PLA_2R1^+$-MN patients P2-P4 contain disease associated proteins. | *P17* |
| *Figure S5.* | Glomerular huIgG4$^+$ aggregates within the urinary space contain disease-associated proteins in the diagnostic biopsy of the THSD7A$^+$-MN patient P1. | *P18* |
| *Figure S6.* | Glomerular huIgG4$^+$ aggregates within the urinary space contain disease-associated proteins in the diagnostic biopsy of the $PLA_2R1^+$-MN patient P2. | *P19* |
| *Figure S7.* | HuIgG4$^+$ aggregates within the tubular lumen contain disease-associated proteins in the diagnostic biopsy of the THSD7A$^+$-MN patient P1. | *P20* |
| *Figure S8.* | HuIgG4$^+$ aggregates within the tubular lumen contain disease-associated proteins in the diagnostic biopsy of the $PLA_2R1^+$-MN patient P2. | *P21* |
| *Figure S9.* | Characteristics of the huIgG4-enriched urinary EV fractions from MCD patients P5, P6 and primary FSGS patients P9, P10 in diagnostic urines. | *P22* |
| *Figure S10.* | Characteristics of the huIgG4-enriched urinary EV fractions from nephrotic patients with IgA nephritis P7 or tubulo-toxic kidney injury P8 in diagnostic urines. | *P23* |
| *Figure S11.* | THSD7A autoantibody binding induces the formation of long membrane extensions with distal bulges from podocytes in experimental THSD7A$^+$-MN. | *P24* |
| *Figure S12.* | THSD7A autoantibody binding induces exopher-formation in podocytes experimental THSD7A$^+$-MN. | *P25* |
| *Figure S13.* | Immunogold EM localization of injected rbTHSD7A-abs day 1 and day 7. | *P26* |
| *Figure S14.* | Urinary exophers-release and uptake in experimental THSD7A$^+$-MN. | *P27* |
| *Figure S15.* | *In vitro* exposure to human THSD7A-abs results in crosslinking of THSD7A. | *P28* |
| *Figure S16.* | *In vitro* exposure of human podocytes to rbTHSD7A-abs induces release of large rbIgG$^+$ EVs. | *P29* |
| *Figure S17.* | Short-term proteotoxic stress induces specific exopher-release to the urine. | *P30* |
| *Figure S18.* | 'Long-term' proteotoxic stress decreases urinary exopher-release despite persistence of rbTHSD7A-abs in the serum of a THSD7A$^+$-MN mouse. | *P31* |
| *Figure S19.* | Computational image analyses of $PLA_2R1$ aggregate density. | *P32* |
| *Figure S20.* | General workflow of EV sample preparation from patient urine as detailed in the method section. | *P33* |
| *Table S1.* | Clinical parameters of the THSD7A$^+$-MN patient P1 and of $PLA_2R1^+$-MN patients P2-P4 at time of analyses. | *P33* |
| *Table S2.* | Clinical parameters of the nephrotic non-MN patients P5-P10 at time of analyses. | *P34* |
| *Table S3.* | HuIgG4$^+$-EV characteristics in the diagnostic urines (U1) of nephrotic patients | *P34* |
| *Table S4.* | Proteomics (attached excel list) | |
| *Table S5.* | Antibodies and dyes used in the study. | *P35-36* |
| *Suppl. References* | | *P36-37* |

### AUTHOR / INSTITUTE INFORMATION

**Institute of Cellular and Integrative Physiology, Center for Experimental Medicine, University Hospital Hamburg-Eppendorf and Hamburg Center of Kidney Health, Hamburg, Germany**
*Karen Lahme,*
*Wiebke Sachs*,
*Sarah Froembling*,
*Desiree Loreth,*
*Uta Wedekind*,
*Vincent Böttcher-Dierks*,
*Sinah Skuza*,
*Karen Neitzel*,
*Stephanie Zielinski*,
*Johannes Brand*,
*Catherine Meyer-Schwesinger*,

**Institute of Medical Systems Biology, Center for Biomedical AI (bAIome), Center for Molecular Neurobiology (ZMNH), University Medical Center Hamburg-Eppendorf, Hamburg, Germany**
*Michael Brehler,*
*Stefan Bonn*,
*Marina Zimmermann*,

**Interfaculty Institute for Genetics and Functional Genomics, University Medicine Greifswald, 17475 Greifswald, Germany**
*Kristin Surmann,*
*Stephan Michalik,*
*Uwe Völker*,

**Chimaera GmbH, Erlangen, Germany**
*Simone Gaffling,*

**Cell Communication and Migration Laboratory, Institute of Biochemistry and Molecular Cell Biology, Center for Experimental Medicine, University Medical Center Hamburg-Eppendorf, 20246 Hamburg, Germany**
*Marie R. Adler*,
*Pablo J. Sáez*,

**Technology Platform Microscopy and Image Analysis (TP MIA), Leibniz Institute of Virology, Hamburg, Germany**
*Christian Conze,*
*Roland Thünauer,*

**Institute of Pathology, Nephropathology Section, University Medical Center Hamburg-Eppendorf, Germany**
*Thorsten Wiech*,

**Nephrology, Asklepios Klinikum Barmbek, and Faculty of Medicine, Semmelweis University Budapest, Asklepios Campus Hamburg, Hamburg, Germany**
*Tobias N. Meyer,*

**Institute of Neuroanatomy, Medical Faculty, University of Bonn, Bonn, Germany and Institute of Anatomy and Cell Biology, Friedrich-Alexander-Universität Erlangen-Nürnberg, 91054 Erlangen, Germany**
*Lars Fester,*

### SUPPLEMENTARY METHODS

The presented research complies with all relevant ethical regulations. Written consent by the patients and ethics approval was obtained for the urine analyses and correlations to diagnostic biopsy samples (#2023-101080-BO-ff). For the retrospective $PLA_2R1$-MN patient analyses approval was obtained by the Hamburg GN Registry board: Dr. Wiech, Dr. Hoxha, Dr. T.B. Huber. For the generation of human glomerular extract, the healthy (tumor-free) part of the kidney not needed for pathological diagnosis was used. Patients provided written consent for the use of samples for research. Because patients were anonymized, information on age and gender are not available. Similar samples cannot be accessed by external users. Animal studies were approved by the Behörde für Justiz und Verbraucherschutz, Amt für Verbraucherschutz, Lebensmittelsicherheit und Veterinärwesen, Hamburg, Germany.

#### Antibodies and dyes

Primary antibodies and dyes used for this study are listed in ***Table S5***.

#### Cell culture

*Human podocytes:* Human immortalized podocytes (a kind gift of Moin Saleem, University of Bristol, Bristol, Great Britain) were cultured under permissive conditions [32°C, 5% $CO_2$, RPMI 1640 Thermo Fisher, #61870-010 supplemented with 10% fetal calf serum, 1X insulin, transferrin selenium (ITS, PAN Biotech, #P07-03200), 1 µg/ml Puromycin (Sigma, #P7255)] in uncoated, vented tissue culture flasks (Sarstedt) as described[1]. Stable overexpression of human THSD7A with a C-terminal myc-Flag tag was achieved by lentiviral transduction. Mock cells were transduced with an empty vector[2]. For differentiation, podocytes were cultured for 10 days under non-permissive conditions [37°C, 5% $CO_2$, RPMI 1640 supplemented with 10% FCS, 1X ITS]. Cell density was kept below 80-90% to allow process development. DNA was extracted from cell pellets and controlled for the absence of mycoplasma infections on a 2-3 monthly basis according to the manufacturer's instructions (Minerva Biolabs #11-8100), cellular passages below 60 were used for experiments. To collect extracellular vesicles, podocytes were transferred to culture in an exosome-depleted medium (Thermo Fisher, #A2720803) 16 hours prior to commencement of antibody treatment. *In vitro* antibody treatment was performed with human or rabbit anti-THSD7A-specific antibodies or unspecific control huIgG or rbIgG, all 1:200 from a 4 mg/mL stock solution) for 0, 1, 6, 24, and 48 hours. Podocytes were harvested by scratching in PBS on ice. Cell pellets were collected and lysed in T-PER (Thermo Fisher, #78510) with SigmaFast EDTA-free protease inhibitor cocktail (Sigma, #S8830). Extracellular vesicles (EVs) were collected from 240.000 cells in 3 ml exosome-depleted medium.

*HEK293T cells:* HEK293T cells (Sigma, # 12022001) were cultured in DMEM supplied with 10% FCS and 1X penicillin/streptomycin (Thermo Fisher, #15070063) at 37°C and 5% $CO_2$. Transient transfection of HEK293T cells with full length human (hu)THSD7A (Origene, #RC213616) or $huPLA_2R1$[3] constructs was performed using JetPEI (VWR, #101000053) according to the manufacturer's instruction. In brief, 5 µg of respective plasmid DNA was mixed with 10 µl of JetPEI and 10 mM sodium chloride in a final volume of 500 µl and after incubation for 25 minutes at room temperature gently added to < 80% confluent HEK293T cells that were seeded at the same time day of transfection and allowed to attach to the bottom of a 6 cm cell culture plate (Sarstedt, #83.3901). Cells were allowed to express the protein for 48 hours at 37°C and 5% $CO_2$. Cells were harvested by scraping ice cold PBS (Thermo Fisher, #14190-169). The resulting cell suspension was transferred to a 1.5 ml Eppendorf tube, centrifuged at 4°C at 900 x *g* for 10 minutes and the supernatant was discarded. To remove remaining medium residues, the pellet was resuspended in PBS, centrifuged again at 4°C at 900 x *g* for 10 minutes, supernatant was discarded, and the pellet was lysed in T-PER (Thermo Fisher, #78510) after addition of Proteinase Inhibitor Cocktail (Sigma, #S8820) and PhosStop (Sigma, #4906845001). After protein quantification by the use of ROTI®Quant universal (Carl Roth, #0120.1) according to the manufacturer's instruction, 10 µg of total protein was prepared for immunoblotting under non-reducing conditions (loading buffer as described below but without DTT) for detection of autoantibodies to huTHSD7A or $huPLA_2R1$ in murine sera or in the eluted antibody fraction from enriched patient exophers.

**Phase contrast live cell imaging**

Podocytes were seeded in 35 mm diameter bottom glass fluorodishes (World Precision Instrument, WPI) coated with Poly-L-Lysine. Imaging was performed with a Leica DMi8 M / C / A inverted microscope using a HC PL Fluotar 20x/0,4 CORR PH1 objective (Leica Microsystems). Podocytes were imaged for 4 hours; treatment was started 30 minutes prior to commencement of imaging with addition of rbTHSD7A-abs or ctrl-abs to the medium. The images were recorded with an ORCA-Flash4.0 Digital camera (Hamamatsu Photonics), and Meta-Morph® Version 7.10.5.476 software (Molecular Devices). Pictures were acquired every 1 min for a total duration of 4 hours. Movies were reconstituted, processed, and analyzed using Fiji (ImageJ 1.53t) software. The tracking was performed using the TrackMate v7 plugin from Fiji[4,5] after processing the movies as previously described[6]. During the acquisition, podocytes were kept at 37°C, and 5% $CO_2$ inside a humid chamber on the objective stage.

**Holotomography**

Refractive index detection and holotomographic images were acquired in a Nanolive 3D Cell Explorer-fluo (Nanolive SA, Tolochenaz, Switzerland), equipped with a 60x/0.8 objective, and STEVE software (Full v1.6.3496), as previously described[7]. Podocytes were imaged 45 minutes after treatment with rbTHSD7A-abs or ctrl-abs as described in the previous sections. For the acquisition, the cells were kept at 37°C, and 5% $CO_2$ inside a humid chamber on the objective stage. The holotomographic images were processed by using the post-processing plugins in STEVE. According to the manufacturers' instructions we performed the following image processing: first a histogram a log10 adjust was made, followed by a background subtraction (automatic estimation). No edge preserving filter was applied. The 3D images of the refractive index were made later in Fiji (ImageJ) by using the plugin 3D project.

**Animal experiments**

Male BALB/c mice with a pure genetic background were purchased from Charles River, *Gt(ROSA)26Sortm4(ACTB-tdTomato,-EGFP)Luo/J x hNPHS2Cre mice* (*mT/mG*)[8] with a BALB/c background were provided from the Huber laboratory (III Medical Clinic, UKE, Hamburg-Eppendorf) and were predominantly analyzed at 16-22 weeks of age. *mT/mG* reporter mice express membrane-bound GFP in podocytes and all other renal cells express membrane-bound tdTomato. Mice had free access to water and standard animal chow (Altromin 1328 P), were synchronized to a 12:12 hour light-dark cycle and were housed in a pathogen-free animal facility at the University Medical Center Hamburg-Eppendorf. We concentrated on THSD7A turnover in this study, which contrasting $PLA_2R1$, is constitutively expressed in rodents[9], and reliable mouse models of $THSD7A^+$-MN have been developed[10,11]. In the passive model of $THSD7A^+$-MN[10], mice are exposed to rabbit anti-THSD7A antibodies (rbTHSD7A-abs) by intravenous injection. These rbTHSD7A-abs bind to the large extracellular region of THSD7A as a type 1 transmembrane protein with a single transmembrane domain, and a short intracellular domain[12]. Similarly to the patient autoantibodies to THSD7A, rbTHSD7A-abs exhibit a binding affinity to THSD7A that is sensitive to the conformation of the protein[10,13]. THSD7A localizes to the most basal aspect of the podocyte foot process domain near the slit diaphragm underneath Nephrin[2]. Accordingly, autoantibody binding rapidly occurs at the subepithelial aspect of FPs, preferentially at the slit diaphragm[2] and mice develop proteinuria after 1-3 days, which reaches nephrotic ranges after 7 days. In this model, glomerular accumulations of THSD7A and rbTHSD7A-abs are observed in a comparable pattern to patients with $THSD7A^+$-MN[10], albeit to a lesser degree.

Here, $THSD7A^+$-MN was induced by intravenous injection of 180 µl (1,5 mg/ 30 g mouse) rabbit anti-human/mouse THSD7A antibodies or unspecific rabbit IgG as a control in male BALB/c or *mT/mG* mice aged 12-22 weeks[10]. For 'short-term' inhibitor studies mice were treated on four consecutive days with either the proteasomal inhibitor epoxomicin (0.5 mg/kg bodyweight in 125 µl, Enzo, #BML-PI127-0100) or vehicle (25% DMSO in 125 µl, Carl Roth, #4720.2) by intra-peritoneal injection. For 'long-term' proteasome inhibitor studies, treatment with the proteasomal inhibitor bortezomib (0.5 µg/g bodyweight in 125 µl DMSO; PS-341, UbpBio, #F1200) or vehicle (25% DMSO in 125 µl, Carl Roth, #4720.2) was commenced one day prior to THSD7A-MN induction by intra-peritoneal injection 2 times/ week for 11 days. Animal euthanasia was performed following subcutaneous buprenorphine hydrochloride (0,1 mg/kg KG, TEMGESIC, Indivior Europe Limited, #ind00979) administration for analgesia 30 minutes prior to cervical neck dislocation under 3.5% isoflurane (Baxter, #HDG9623) inhalation narcosis. During all procedures, body temperature was kept at 37 $\pm$ 0.5°C. All experimental procedures were performed according to the institutional guidelines.

**Sample collection**

For sample collection, mouse urine was collected in metabolic cages over a period of 4 hours. Isolated kidney packages were thoroughly perfused through the renal arteries with 5 ml PBS/Dynabeads$^{TM}$ M-450 Tosylactivated (Invitrogen, Thermo Fisher, #4013) per kidney to remove blood components such as unbound immunoglobulins from the renal circulation. Successful perfusion was controlled by the color of the kidney. Samples were stored at -80°C prior to further analysis, fixed in 4% PFA for histological evaluations, or processed for glomerular isolation.

**Mouse serum analysis**

Mouse blood was centrifuged for 10 min at 1500 x *g*, 4°C to separate blood cells from serum. In serum circulating specific rabbit THSD7A antibodies were detected by immunoblotting. For this, huTHSD7A and huPLA$_2$R1 protein produced in HEK cells was loaded non reduced on a SDS-PAGE. Mouse serum (300 µl) was mixed 1:2 with SuperBlock$^{TM}$ (Thermo Scientific #37515), incubated as primary antibody on the WB membrane over night at 4°C and detected with anti-mouse HRP secondary antibody.

**Urinary albumin to creatinine ratio**

Urine albumin content was quantified using a commercially available ELISA system (Bethyl, Montgomery, USA, #A90-134A) according to the manufacturer's instructions. Briefly, 96-well plates were coated 1:100 with goat anti-mouse albumin in binding buffer (0.05 M carbonate-bicarbonate pH 9.6) over night at 4°C. After washes in 50 mM Tris, 0.14 M NaCl, 0.05% Tween-20 pH 8.0, the plates were blocked for 30 min at room temperature with 50 mM Tris, 0.14 M NaCl, and 1% BSA pH 8.0 and rewashed. Diluted urine was incubated for 60 min at RT. Following washes, the secondary antibody (1:40,000; horseradish peroxidase goat anti-mouse albumin) was applied for 60 min at room temperature. After washes, enzyme substrate supplied by the kit was added and the color development was stopped after 5 min with 2 M phosphoric acid. Extinction was measured at 450 nm in an ELISA plate reader (BioTek, #EL 808). The urinary albumin concentration was calculated according to the formula for absorption = $(A - D)/1 + (x/C)^B + D$, where A and D are values from the standard curve. Regression values for the standard curve were calculated to assess the accuracy of the measured values. Standard curves with $r$ values > 0.9950 were used. Urinary albumin values were standardized against urine creatinine values of the same individuals determined by Jaffé (Hengler analytic, #114444) or to the urine volume and plotted as [g albumin / g creatinine] or as [g albumin / µl urine].

**Glomeruli isolation**

*Murine glomeruli* were isolated using Dynabead perfusion as described[14]. In brief, kidneys were perfused with 5 ml magnetic bead solution per kidney (magnetic bead solution: of 50 µl Dynabeads™; M-450 Tosylactivated Invitrogen ThermoFisher #14013 with 200 µl SPHERO Polystyrene Magnetic Particles, Spherotech, #PM-40-10 diluted in 50 ml HBSS) and digested with collagenase (in HBSS, 1.2 mg/ml collagenase 1A, Sigma, #C9891, 100 U/ml DNaseI, Sigma, #10104159001) for 15 min, 37°C, 1.300 rpm. The solution was strained through a 100 µm cell strainer (Sarstedt, #83.3945.100), which was rinsed with HBSS, this was done 3 times. The solution was centrifuged for 5 min, 600 x *g*, 4°C. The pellet was resuspended in HBSS, and the glomeruli were separated from tubuli with a magnetic particle concentrator. The number of isolated glomeruli and contaminating tubuli was quantified under the phase contrast microscope prior to storage of glomeruli at -80°C.

*Human glomeruli* were used as positive controls for immunoblots. Healthy parts of kidneys from patients who underwent nephrectomy were used for the preparation of an extract of human glomeruli. Cortex was separated from the medulla, the cortical tissue was weighed and pulpified with a scalpel. Approximately 0.5 g tissue was portioned into each reaction tube and digestion solution (1.2 mg/ml Collagenase 1 A, 100 U/ml DNase I) was added. Digestion was performed for 15 min, 37°C, 1.300 rpm in a ThermoMixer (Eppendorf). The tissue homogenate was sieved through a 212 µm sieve placed on a beaker and pushed through with a small Erlenmeyer flask using low pressure. In between the tissue was rinsed multiple times with PBS containing 0.05% BSA. To finally separate glomeruli from tubular remnants, 53 µm sieves were used whereby intact glomeruli remained on the sieve. By using a syringe filled with PBS containing 0.05% BSA, a small cannula, and high pressure the glomeruli were flushed into a clean beaker. To increase the purity grade of the glomerular isolate, the last step was performed two times, using clean 53 µm sieves each time. The collected suspension was centrifuged for 10 min at 800 x *g*, 4°C. The supernatant was decanted, and the pellet was solved in PBS containing 0.05% BSA. The number and purity of isolated glomeruli was determined under a phase contrast microscope.

**Morphological analyses**

*Cleared kidney slices:* Kidney pieces from *mT/mG* mice were immersion fixed with 4% PFA o/n at 4°C and sliced 300 µm thick with a Vibratome (VT1000S, Leica Biosystems Nussloch GmbH, Germany), washed with PBS and optically cleared for 48 hours using SCALEVIEW A2 (FUJIFILM Wako Chemicals, #193-18455). For immunofluorescence of exophers, cleared slices were incubated with cy5 anti-rabbit IgG (1:100 for 3 days). After extensive washing over 48 hours, 3D image stacks were acquired using a Nikon A1 R confocal laser scanning microscope with A 60x CFI Plan Apo Lambda immersion oil objective (NA 1.42). Additionally, a Visitron Spinning Disk Microscope equipped with a 100x CFI Plan Apo Lambda (NA 1.45) objective and a Yokogawa CSU W-1 Spinning Disk unit with 50 µm pinholes and a Photometrics Prime 95B (back-illuminated sCMOS camera) was employed. The mGFP and cy5 tags were visualized using solid-state lasers of 488, 640, and 647 nm and image stacks were recorded with 0.130 and 0.150 µm Z-steps, respectively. The 3D image deconvolution was carried out with Nikon's NIS-elements AR 5.42.02 using a theoretical PSF and employing an automatic deconvolution algorithm. 3D image data sets were subsequently further processed using Nikon Denoise.ai and Clarify.ai tools. Visualization of the 3D volume views of whole glomeruli as well as 2D MIPs from the podocyte cell body area were carried out using NIS-elements. 3D reconstruction of the image stacks in blend mode from the podocyte cell body area was rendered by utilizing Imaris 8.2.1. Brightness, contrast, and gamma were adjusted for optimal representation.

*Light microscopy:* For light microscopic evaluation, 1.5 µm thick paraffin sections were stained with periodic-acid-Schiff reagent (Sigma, #1.08033.0500) according to the manufacturer's instructions for the assessment of general renal morphology in the THSD7A$^+$-MN patient. Nuclei were counterstained with hematoxylin (Serva, #24420.02).

*Immunohistochemical analyses:* 1 µM paraffin sections of the diagnostic renal biopsy of the THSD7A$^+$-MN patient were deparaffinized and rehydrated. Antigen retrieval was obtained for THSD7A by boiling in DAKO antigen retrieval buffer, pH 9 (15 min at 98°C; DAKO, #S2367) and subsequent cooling and for huIgG4 and huIgG by protease XXIV (5 *µ*g/ml; Sigma, #8038) for 15 minutes at 37°C and stopping of the reaction in 100% EtOH. Nonspecific binding was blocked with 5% horse serum (Vector, #VEC-S-2000) with 0.05% Triton X-100 (Sigma, #T8787) in PBS for 30 min at RT prior to incubation at 4°C o/n with rabbit anti-THSD7A, mouse anti-huIgG4, or goat anti-huIgG in blocking buffer. Staining was visualized with the ZytochemPlus AP Polymer kit (Zytomed Systems, Berlin, Germany, #POLAP-100) according to the manufacturer's instructions. Nuclei were counterstained with hematoxylin (Serva, #24420.02) and sections were mounted with gum arabic (Roth, #4159.3). Stainings were evaluated with an Axioskop using the Axiovision software (all Zeiss, Oberkochen, Germany).

*Immunofluorescent analyses paraffin sections:* Patient biopsy and mouse kidney paraffin sections were deparaffinized and rehydrated. Antigen retrieval was obtained by i) boiling in retrieval buffer, pH9 (#S2367, DAKO), pH6.1 (#S2369, DAKO) or 0.05% citraconic acid (#82604, Sigma) for 30 min at 98°C, followed by cooling or ii) protease XXIV (Sigma, #8038, 5 µg/ml) digestion 15 min at 37°C followed by a wash in 100% EtOH. Unspecific binding was blocked with 5% normal horse serum (#VEC-S-2000) in

PBS 0.05% TritonX-100 (Sigma, #T8787) for 30 min at RT. Primary antibodies were incubated in blocking buffer o/n at 4°C. Following washes in PBS, fluorochrome-labeled donkey secondary antibodies (all 1:200, Jackson Immunoresearch Laboratories), biotinylated-WGA (followed by AF647-streptavidin) or rhodamine-coupled WGA, and Hoechst were applied where appropriate for 30 min at RT. After washes in PBS, sections were mounted in fluoromount (Southern-Biotech, #0100-01).

*Immunofluorescence of urinary exophers:* Urinary EVs were stained using the image stream antibodies and according to the method described in the paragraph image stream. For additional analyses of the exopher content, the staining panel was expanded to include the proteins of interest (see ***Table S5*** for antibodies and dyes used). Stained and washed EVs were airdried on poly-L-lysine (1:1000 in sterile water) coated slides and mounted with fluoromount.

For immunofluorescence of cultured podocytes, cells were extensively washed after live-imaging, fixed with 4% PFA for 8 min at RT. Following PBS washes, unspecific binding was blocked with 5% normal horse serum (#VEC-S-2000) in PBS 0.05% TritonX-100 (Sigma, #T8787) for 30 min at RT. Primary antibodies were applied o/n in blocking buffer at 4°C. The following day, samples were PBS washed and incubated with secondary antibodies (all 1:200) in blocking buffer for 30 min. Samples were PBS washed and mounted with fluoromount.

Immunofluorescence was resolved by conventional and high-resolution confocal microscopy either with a LSM800 with Airyscan 1 or with a LSM980 with Airyscan 2 microscope using ZEN 3.0 software (all ZEISS).

**Computational image analysis**

35 patient biopsies of a retrospective $PLA_2R1^+$-MN patient cohort were stained for $PLA_2R1$. Each glomerulus present within the biopsy cylinder was imaged using an LSM800 in confocal modus at a 2873 x 2873 resolution, Plan-Apochromat 63x/ 1.40 Oil DIC M27 objective, pinhole 1AU. The algorithm was implemented in Python 3 (mainly packages Scikit-image and OpenCV) using a Difference of Gaussian blob detection[15]. Algorithm inputs were images saved in Carl Zeiss Image Data file format. Using the grayscale $PLA_2R1$ channel, images were first thresholded with a Multi-Otsu approach[16] separating high signal areas from background noise. The binary image was further processed removing very small noisy pixels and applying the morphological operation of closing. Based on the now cleared binary image a glomerular mask was extracted by calculating the convex hull around the binary signal and therefore roughly following the Bowman's capsule / the glomerulus outline. In the masked area signal clusters (areas of overlapping or touching signal blobs) were separated from each other using a watershed transformation[17]. In the last step, the binary output of the watershed separation is used as input for the Difference of Gaussian blob detection. Based on the detected blobs the $PLA_2R1$ density for each blob is calculated. $PLA_2R1$ density is defined as the sum of the number of neighboring blobs (excluding the current blob) within a radius of 50 pixels around each blob divided by total number of blobs in the image. The division normalizes the image to a range from 0 (all blobs have 0 neighbors) to the total number of blobs (all blobs have all blobs as neighbors). The $PLA_2R1$ aggregated density per patient is calculated as the median of all $PLA_2R1$ densities of all images of that patient. All code will be made available via GitHub. Please refer to ***Fig. S19*** for further details.

**Ultrastructural analyses**

*Pre-embedding Immunogold EM:* Experimental mice were *in vivo* fixed for immunogold labeling modified according to Somogyi and Takagi (1982)[18]. In brief mice were transcardially perfused with 0.1 M phosphate buffer (PB) pH 7.4 at 4°C. One liter fixative consisted of 500 ml 0.2 M PB pH 7.4, 150 ml saturated picric acid (Bouin's solution: 0.044 M picric acid, 5% (V/V) acidic acid, and 10% (V/V) formaldehyde in aqua dest.), 346.8 ml paraformaldehyde solution (40 g PFA in aqua dest.) and 3.2 ml of 25% glutaraldehyde at a final pH 7.4. The final concentrations were 4% paraformaldehyde, 0.08% glutaraldehyde, and 0.15% picric acid. Kidneys were removed after perfusion procedure and postfixed in 100 ml 0.1 M PB including 4% paraformaldehyde and 0.15% picric acid for 24 hrs at 4°C. Vibratome sections (LEICA VT 1000S) of fixed tissue were made with a thickness of 60 µm of areas of interest and the slices were placed into cryoprotection solution containing 0.1 M PB (pH 7.4), 30% sucrose, 1% polyvinylpyrrolidone, and 30% ethylenglycol. For pre-embedding immunogold staining, cryoprotection solution was removed from the 60 µm thin slices by washing twice in 0.1 M sodium phosphate buffer pH 7.4 (PBS) for 20 minutes, followed by washing of the slices in 0.1 M PBS with 10% sucrose for 30 minutes and 0.1 M PBS with 20% sucrose for further 30 minutes. Tissue was transferred to a drop of PBS with 20% sucrose and frozen briefly with nitrogen and thawed to room temperature. Subsequently, the tissue was washed three times in PBS pH 7.4 for 10 minutes. Blocking reagent solution (BRS) were prepared with 100 ml PBS, 800 mg BSA (Sigma-Aldrich A7638-5G), 250 µl 40% cold water fish skin gelatine (AURION CWFSG 900.033) and 162.5 mg sodium acid. For detection of bound rbTHSD7A-abs, 12 nm Colloidal Gold-AffiniPure Donkey anti-rabbit IgG (H+L) (Jackson ImmunoResearch) was used. For the detection of THSD7A antigen, indirect immunostaining was performed, using the affinity-purified, pathogenicity-inducing patient THSD7A autoantibody[19] as primary antibody to omit the risk of cross-reactivity to the injected and bound rabbit THSD7A-abs and to the intrinsic mouse IgG within the sample. The bound patient THSD7A autoantibody was then detected by 12 nm Colloidal Gold-AffiniPure Goat anti-human IgG (H+L) (Jackson ImmunoResearch). To increase the specificity of primary antibody reaction, blocking reagent solution (BRS) supplemented with 2% normal goat serum (NGS, Sigma) was used for one hour for blocking reaction. The primary antibody was solved in BRS overnight at 4°C. After 24 hours the tissue was washed in PBS 3 times for 5 minutes each and incubated with 12 nm gold conjugated antibody solution (1:50) in BRS for 2 hours. Subsequently, the tissue was washed three times in PBS pH 7.4 every 5 minutes, followed by 1% glutaraldehyde solution in 0.1 M PB for 10 minutes. The tissue was solved in 1% OsO4 in aqua dest. at 4°C for 10 minutes followed by dehydration in an ethanol series. Subsequently, the probes were transferred twice into propylene oxide for 5 minutes and embedded in a mixture of epon and propylene oxide (4:1) overnight. Tissue samples were cut with an ultra-thin (Reichert Jung Ultracut) microtome and thin sections were inspected in a Zeiss electron microscope (TEM Leo 910) and TEM Jeol 1400plus. Pictures were taken assisted by ImageSPViewer (Tröndle) and image viewer Jeol.

*3D reconstruction:* 3D reconstruction of 13-48 consecutive EM micrographs was performed using the HiD® Histodigital reconstruction approach[20]. First, the orientation of the micrographs was homogenized using rigid image registration, such that all images were aligned. Since the thin sections are deformed during cutting and further processing, they are affected by non-linear deformations which prevent a coherent 3D impression even after slice alignment. Therefore, in a second step, an intensity-based non-rigid registration routine is applied to reverse these deformations such that spatial coherence of the data set is achieved. Volume visualizations were created with the Chimaera SDK.

*Scanning electron microscopy (SEM):* For evaluation in SEM, bouin fixed 60 µm mouse kidney slices were incubated in series of ethanol/aqua dest. bath with increasing ethanol concentration (30%, 50%, 70%, 90%, 100%,) at ten minutes intervals followed in acetone/EtOH bath by increasing acetone concentration (50%, 70%, 90%, 100%) at ten minutes intervals. Then tissue probes were dried in critical point dryer (Bal-Tec CPD 030) and stored in Vacuum dryer. The dried tissue probes were placed with conducting silver on carbon plate on a metallic probe holder and dried again in vacuum chamber for 24 hours. The surface was sputtered by 5 nm thin platinum layer in a sputter coater (Bal-Tec SCD 500) and evaluated on SEM Jeol 7500F via YAG and LABE Backscatter electron detector (BSE).

#### Extracellular vesicle (EV) isolation

*EV isolation from human podocyte culture:* Medium was collected from cells and fixation solutions were added to the collected cell culture medium [1:100; solution 1: 660 mM $NaN_3$ (Sigma, #S2002), 200 mM EGTA (Carl Roth, #3054.2) in dd$H_2O$, solution 2: 100 mM PMSF (Merck-Millipore, #P7626) in 100% ethanol]. The medium was centrifuged for 10 min, 2.000 x *g*, 4°C, the supernatant was removed and centrifuged for 10 min at 2.000 x *g* and 4°C. The supernatant was ultra-centrifuged for 1.5 hours at 100.000 x *g* and 4°C, the supernatant was discarded, and the pellet resolved in filtered PBS.

*EV isolation from mouse urine:* Fixation solution (1:100 dilution [660 mM $NaN_3$, 200 mM EGTA in $ddH_2O$, solution 2: 100 mM PMSF in 100% ethanol]). Urine samples were centrifuged for 10 min at 5.000 x *g* and 4°C. To reduce the protein amount of the nephrotic urine the supernatant was dialyzed for 20 hours in 1 x PBS at 4°C on a magnetic stirrer, using a 100 kDa cut-off dialysis membrane Spectra/Por Biotech CE (Carl Roth, #8967.1). The dialyzed urine was ultra-centrifuged for 1.5 hours at 100.000 x *g* and 4°C. The supernatant was discarded, and the pellet solved in filtered PBS. Concentration was measured with MemBrite® Fix Cell Surface dye 594/615 by Image Stream.

*EV isolation from patient urine:* The general workflow for total urinary EV isolation and subsequent $huIgG4^+$-EV enrichment for the individual analyses (image stream, immunofluorescence, immunoblotting, NTA, autoantibody elution, proteomics) is summarized in ***Fig. S20.*** Nephrotic patients admitted to the clinic for a diagnostic kidney biopsy collected either a 24-hour urine or spot urine prior to the kidney biopsy. Urine was processed by addition of fixation solution (1:100; [660 mM $NaN_3$, 200 mM EGTA in $ddH_2O$, solution 2: 100 mM PMSF in 100% ethanol]) and centrifuged for 10 min at 2.500 x *g*, 4°C. The supernatant was filtered through 5 µm syringe filters (Millex Merck Millipore, #SLSV025LS) to remove cell debris. 50 ml aliquots were stored at -80°C prior to use. Patients were included to the study after serological and nephro-pathological diagnosis of following diseases: $THSD7A^+$-MN (Patient (**P**)**1**), $PLA_2R1^+$-MN (**P2**-**P4***)*, minimal change disease (**P5**, **P6**), IgA nephritis (**P7**), tubulo-toxic kidney injury (**P8**), and primary focal segmental glomerulosclerosis (pFSGS: **P9**, **P10**). Two patients (**P1** and **P2**) were prospectively followed, and urinary samples were collected within their routine clinical management. Stored urine samples were thawed over night at 4°C, thoroughly vortexed and centrifuged for 10 min at 2500 x *g*, 4°C to remove aggregates. To reduce the abundance of contaminating proteins from the highly proteinuric urine, the urine was ultra-filtrated by centrifugation at 3.000 x *g* and 4°C using 100 kDa cut-off filter tubes (Amicon Ultra-15 Centrifugal Filter Unit, Merck-Millipore, #UFC910024), until half of the initial volume was reached. The filtered supernatant was ultra-centrifuged for 1.5 hours, 100.000 x *g* and 4°C, the supernatant was discarded, and the EV pellet was solved in filtered PBS. Particles/ml were measured with MemBrite® Fix Cell Surface dye 594/615 by image stream. The resulting EV samples are called total EVs (input) in the following methods (= sample 1, ***Fig. S20***).

### Enrichment of $huIgG4^+$-EVs from total EVs

*Preparation of huIgG4-beads for EV pulldown:* Dynabeads™ M-450 Tosylactivated (Invitrogen ThermoFisher, #14013) were washed with 0.1M sodium phosphate buffer pH 8.0 on a DynaMag. Anti-human IgG4 antibody (Mouse Anti-Human IgG4 pFc'-UNLB, Southern-Biotech, #9190-01) was coupled to Dynabeads according to manufacturer's instructions. Antibody coupled beads were blocked 1:1 with 2% cold fish gelatine (Sigma, #G7041) and SuperBlock™ (Thermo Scientific, #37515) in filtered PBS for 1 hour at room temperature on a roller shaker. Beads were washed on a DynaMag twice with 0.1M sodium phosphate pH 8.0.

*$HuIgG4^+$-EV pulldown:* Depending on the planned assay, $10^5$ (immunoblotting), $10^9$ (proteomic analyses), $10^9$ (image stream, NTA), or $10^{10}$ (autoantibody elution) total EVs (= sample 1, ***Fig. S20***) were used as starting material. Total EVs were added to blocked huIgG4-beads, filled up to 250 µl with 2 mM EDTA in filtered PBS pH 7.4 and incubated overnight on a roller shaker at 4°C. EVs bound to huIgG4-beads were concentrated on a DynaMag. Enriched (bead bound) $huIgG4^+$-EVs (= sample 2 in ***Fig. S20***) were eluted from the beads for quantification depending on the planned analyses. In general, EV concentration (particles/ml) was measured by image stream using MemBrite® Fix Cell Surface dye 594/615. The supernatant containing the rest EV fraction ($huIgG4^+$-EV depleted) was removed and processed by ultra-centrifugation for 1.5 hours at 100.000 x *g* and 4°C to pellet the rest EVs (= sample 3 in ***Fig. S20***). The rest EV pellet was dissolved in sterile PBS, concentration (particles/ml) was measured by image stream as detailed above.

### Image Stream

Vesicle abundances and marker composition were analyzed by image stream. Following starting material was used: 1) podocyte EVs released into 3 ml of exosome-depleted medium, 2) total EVs isolated from 250 µl of mouse urine 3) $10^9$ EVs isolated from patient urine. To determine the particle/ml concentration of individual EV sample preparations, 3 µl were stained.

*EV staining for image stream: EVs* were stained in filtered PBS containing 8% exosome-depleted FBS (Thermo Fisher, #A2720803) in an end volume of 10 µl and stained for 45 min in the dark at room temperature. Stained EVs were washed in 500 µl 2% exosome-depleted FBS in filtered PBS using 300 kDa filter (PALL Nanosep Centrifugal Devices with Omega Membrane - 300K), centrifuged for 10 min at 5000 x *g* and resolved in wash buffer containing 2% exosome-depleted FCS in filtered PBS. Controls included for all analyses were buffer controls without EVs and antibodies, unstained EV samples, and buffer with antibodies only. All antibodies used for image stream (listed in ***Table S5***) were centrifuged for 10 min at 15.000 x *g* at 4°C before use. Antibodies used to stain murine samples were cy5 anti-rabbit (Jackson ImmunoResearch Laboratories), AF546 anti-14-3-3, CoraLite Plus 647 anti-AnnexinV (Proteintech), and 1x MemBrite ® Fix Cell Surface dye 594/615 (Biotium). For human EVs staining the antibodies used were, CoraLite ® 488 anti-human IgG4 (Proteintech), PacBlue anti-CD63 (BioLegend), PacBlue anti-CD81 (Biolegend), APC anti-AnnexinA1 (Novus Biologicals), CoraLite Plus 647 anti-AnnexinV (Proteintech) and 1x MemBrite® Fix Cell Surface dye 594/615 (Biotium).

*Image stream measurements:* EVs were analyzed on an ImageStreamX MkII (ISX Amnis/MilliporeSigma) with a 60x magnification and low flow rate with standard sheath fluid (Dulbecco's PBS pH 7.4). mGFP, FITC, and cy2 signal were collected in channel 2 (480-560 nm filter), mTomato and AF546 signal in channel 3 (560-595 nm filter), MemBrite ® Fix Cell Surface dye 594/615 signal in channel 4 (595-640nm filter), PacBlue signal in channel 7 (430-505 nm filter) and AF647, cy5 and APC signals in channel 11 (640-745 nm filter). Channels 1 and 9 were used as brightfield channels (430-480 nm and 570-595 nm filter) and channel 6 (745-800 nm filter) for site scatter (SSC) detection. Mouse EV gating: For *mT/mG* reporter mice all fluorescent EVs positive for mTomato (Ch03) and mGFP (Ch02) were used as total vesicle population. Gates only positive for mGFP (podocyte EVs), mTomato (all other EVs), and positive for both signals were set. mGFP$^+$ population or mTomato$^+$ population were analyzed for rbIgG (Ch11) and Annexin V (Ch11) signal (mGFP$^+$ population: Ch11 against Ch02). The same was done for the mTomato$^+$ population (Ch03). For BALB/c mice MemBrite® 594/615 positive EVs were used as total vesicle population. Ch04 MemBrite594/615 fluorescence plotted against SSC Ch06. All events that showed low SSC (<1000) but a fluorescent intensity for MemBrite594/615 were used as total EVs for further analysis. The MemBrite594/615$^+$ population was plotted for the marker rbIgG (cy5 Ch11) and 14-3-3 (AF546 Ch03). Human EV gating: Ch04 MemBrite594/615 fluorescence plotted against SSC Ch06. All events that showed low SSC (<1000) but a fluorescent intensity for MemBrite594/615 were used as total EVs for further analysis. The MemBrite594$^+$ population was plotted for the marker huIgG4 in combination with 14-3-3, Annexin A1, Annexin V, and CD63/81 with uniform gates. Data analysis was performed using Amnis IDEAS software (version 6.2).

#### Nano particle tracking (NTA) analysis

EV size distributions within EV fractions were assessed by NTA analyses. For NTA measurement of patient EV fractions (input, rest EVs, huIgG4$^+$-EVs), $10^9$ EVs per fraction were applied. For this purpose, huIgG4$^+$-EVs were freed from huIgG4-beads via thorough vortexing for 3 min prior to quantification by image stream. For NTA analyses of human podocyte EV size distribution, $10^5$ EVs were used.
Briefly, EVs were diluted at 1:1000 in 1.5 ml PBS and 500 µL were loaded into the sample chamber of an LM10 unit (Nanosight, Amesbury, UK). Measurement settings were set to screen gain = 2 and camera level = 15. Five videos of one minute duration with 25°C temperature control inside the chamber were recorded for each sample. Data analysis was performed with NTA 3.0 software (Nanosight). Software settings for analysis were as follows: detection threshold = 6, screen gain = 10. The maximum particle size detected from the instrument as true particles was 5 µm.

#### Immunoblot

Immunoblots were performed with human podocyte cell lysates, whole glomerular lysates, soluble *versus* insoluble mouse glomerular lysates, human podocyte EV lysates, and human patient urine EV lysates.

*Patient urinary huIgG4⁺-EVs*: HuIgG4⁺-EVs bound to the huIgG4-beads were separated from the beads by thorough vortexing for 3 min. Beads were removed by DynaMag and the supernatant containing the free huIgG4⁺-EVs was removed to determine the EV concentration *via* image stream. $10^4$ enriched huIgG4⁺-EVs were lysed in 10 µl PBS with 2 µl 5x loading buffer [250 mM Tris-HCl pH 6.8, 10% SDS, 0,5 M DTT, 50% glycerol und 0.25% bromophenol blue], incubated for 10 min at RT and loaded on an SDS-PAGE and processed for immunoblotting as detailed below. Of note, EV protein abundances determined by immunoblotting are suitable for qualitative observations ("protein *xy* is present"). Quantitative observations ("protein *xy* is more abundant in sample *A* than in *B*") are limited due to normalization issues: 1) normalization to absolute EV number is problematic due to the size variations of EV populations, which leads to different total protein abundances; 2) normalization to protein content is problematic, as BCA measurements are not reliable from the nephrotic urine despite all attempts to reduce albumin contamination; 3) normalization to home keeper is hampered by the lack of adequate and uniform EV home keepers within different EV fractions.

*Human podocyte culture EVs:* Total EVs isolated from 3 ml exosome-depleted medium were resuspended in 12 µl PBS. Subsequently, 3 µl of a 5-fold loading buffer was added, samples were heated for 10 min at 95°C, cooled and completely loaded for SDS-PAGE analyses.

*Murine glomeruli:* For fractionation into a soluble and insoluble pellet, glomeruli were subsequentially lysed: First, the soluble fraction was obtained by adding 30 µl of T-Per containing 1 mM sodium fluoride (Sigma, #S1504), 1 mM sodium orthovanadate (Sigma Aldrich, #S6508), 100 nM Calyculin A (Sigma, #208851), 1x SigmaFast EDTA-free protease inhibitor cocktail (Sigma, #S8830) per 1000 glomeruli. Glomeruli were mechanically shredded with a pestle within the lysis buffer, incubated for 30 min on ice with vortexing every 5 minutes. Lysed glomeruli were centrifuged for 30 min, 4°C at 16000 x *g* and the supernatant containing the soluble proteins was collected. For the insoluble fraction, the remaining pellet was lysed in 30 µl per 1000 glomeruli urea buffer [8 M Urea, 10 mM DTT in 50 mM Tris, pH 8,0], shredded with a pestle, incubated for 30 min on ice with vortexing every 5 minutes, centrifuged for 30 min at 16000 x *g*, 4°C and the supernatant with insoluble proteins was collected. 6 µl of the soluble and insoluble fractions (according to 200 glomeruli) were then denatured with SDS solubilization buffer [250 mM Tris-HCl pH 6.8, 10% SDS, 0,5 M DTT, 50% Glycerol und 0.25% Bromphenolblue] and loaded on an SDS-PAGE.

In general, samples were separated on a 4-15% MiniProtean TGX gel (BioRad, #4568083, 4568085, or 4568086) in a Tris-glycine migration buffer (0.25 M Tris, 1.92 M glycine, 1% SDS, pH 8.3). Protein transfer was performed in transfer buffer (0.192 M glycine, 25 mM Tris base, 20% EtOH in $H_2O$) in a TransBlot Turbo System (BioRad). After the transfer, all proteins were visualized by Ponceau staining. PVDF membranes (Merck-Millipore, #IPVH00010) were blocked (3% nonfat milk in TBS-T [10 mM Tris pH 7.4, 100 mM NaCl, 0.05% Tween20; all ThGeyer, #11784154, #11646918, #116597745]) prior to incubation with primary antibodies diluted in Superblock blocking reagent (Thermo Fisher, #37535) or non-fat milk in TBS-T. Binding was detected by incubation with HRP-coupled secondary antibodies (1:10.000, 3% nonfat milk in TBS-T). Before and after incubation of the membrane in secondary antibody, the membrane was washed three times for 10 minutes in TBS-T. Protein expression was visualized with Chemiluminescent Substrate West Pico Plus (Thermo Fisher, #34578) or Chemiluminescent Substrate Femto Super Signal Maximum Sensitivity Substrate (Thermo Fisher, #34095) according to manufacturer's instructions on Amersham ImageQuant 800 (Cytiva). Western blots were analyzed using software from ImageJ[21]. Normalization was performed to either ponceau, β-actin, or vesicular markers stained on the same membrane for densitometric quantification. If experiments from different membranes were pooled, an internal calculation of relative changes to control within one membrane was assessed. Bands of the same membrane are shown, fine dashed white lines indicate, where bands were not adjacent to another on the membrane.

### Autoantibody elution from urinary patient vesicles

Autoantibodies were eluted from the HuIgG4-bead-bound EVs (= sample 2, ***Fig. S20***) as follows. Beads with bound EVs were incubated in 50 µl 0.1M glycine pH 3.5 for 3 min and vortexed vigorously for 1 min. Similar volume of neutralization buffer (0.1M Tris pH 8) was added. The first supernatant containing the EVs and autoantibodies was collected on a DynaMag. To elute remaining EVs and autoantibodies, beads were reincubated with 0.1M glycine pH 3.5, vortexed and neutralized. The second supernatant was pooled with the first one. Beads were removed by DynaMag and the EV concentration was measured by image stream. To remove the EVs from the supernatant, the supernatant was ultra-centrifuged for 1.5 hours, 100.000 x *g* at 4°C. The supernatant containing the eluted autoantibodies (= sample 4, ***Fig. S20***) was stored for subsequent use as primary immunoblot antibody.

**Native proteasome activity**

Native proteasome activity assay was performed with human podocyte cell and EV lysates. 2.4 x $10^5$ cells and 1.3 x $10^6$ EVs (as determined by image stream) were lysed in TSDG buffer [10 mM Tris, 10 mM NaCl, 25 mM KCl, 1 mM MgCl2, 0.1 mM EDTA, 10 % glycerol, 1mM DTT, 2 mM ATP, pH 7.5]. Cells were lysed in 40 µl TSDG buffer, EVs in a total volume of 11.52 µl TSDG buffer by performing 7 freeze and thaw cycles on a mixture of pure ethanol and dry ice and a 42°C water bath. Cell lysates were centrifuged for 10 minutes at 16000 x *g* at 4°C. Cell lysate supernatant (5 µg) and total EV lysates were used for proteasome activity measurements with the pan-proteasomal activity-based probe (ABP) cy5-epoxomicin (kind gift from B. Florea, Bio-organic synthesis group, Leiden University, Leiden, the Netherlands). Lysates were incubated with 4 µM cy5-epoxomicin for 1h at 37°C in a total volume of 12 µl. Samples were mixed with 5 x native sample buffer [250 mM BisTris, 250 mM NaCl, 50% glycerol, pH 6.5] and directly loaded to a 3-12% native gel (Serva, #43250). Proteins were separated for 3 hours at 150 V in native running buffer [50 mM BisTris, 50 mM Tricin, 0.4 mM ATP, 2 mM MgCl2, 0.5 mM DTT]. Protein transfer and immunoblot were performed as described. 20S Proteasome loading was assessed by blotting for the $\alpha$3-subunit, which is present in all proteasome constitutions.

**Proteomics of HuIgG4$^+$-EVs**

The huIgG4-bead-bound EVs (= sample 2, ***Fig. S20***) were washed and freed from the beads as follows. 50 µl 0.15M sodium carbonate pH11 was added, incubated for 1 min, vortexed thoroughly for 2 minutes and neutralized with similar volume of 0.1M Tris pH8.0. On a DynaMag the supernatant containing the huIgG4$^+$-EVs was removed, and the beads were incubated again with 0.15 M sodium carbonate pH 11.0, vortexed and neutralized to free remaining EVs from Beads. Beads were removed by DynaMag and EV concentration was measured by image stream. HuIgG4$^+$-EVs were ultra-centrifuged for 1.5 hours at 100.000 x *g*, 4°C. Supernatant was discarded and EVs were prepared for proteomics. To do so, EVs were lysed in 20 mM HEPES, pH8, with 1% (v/v) sodium dodecyl sulfate (SDS) followed by 1 min incubation at 95°C. Extraction was enhanced by one freezing step in liquid nitrogen and thawing at room temperature while shaking for 15 min at 1400 rpm (Eppendorf). After a 5 min incubation of samples in an ultrasonication bath, samples were centrifuged for 60 min at 17,000 x *g* and room temperature. The supernatant containing soluble proteins was subjected to protein concentration determination using a BCA assay with bovine serum albumin as standard (Micro BCA Protein Assay Kit, Thermo Fisher Scientific).

Each 1.5 µl sample referring to 3*$10^5$ EVs, were supplemented with 550 ng $^{15}$N-labelled protein extract of *Bacillus subtilis* as normalization standard and filled up with HEPES 20 mM, pH 8 to 10 µl. Then, disulfide bounds were reduced for 30 min with dithiothreitol (2.5 mM final concentration) and free SH-groups at cysteine were subsequently alkylated using iodoacetamide (10 mM final concentration) for further 30 min in the dark. Then, 8 µl of 20 µg/µl SeraMag bead mix were added to each sample in order to prepare protein extracts for tryptic digestion (protein : trypsin ratio 25 : 1) using the SP3 bead-based protocol[22]. After stopping the trypsin reaction with 0.5% (v/v) trifluoric acid and separation peptides from the beads using a magnetic rack and centrifugation, peptides were investigated by nanoLC-MS/MS using an Orbitrap Exploris™ mass spectrometer (Thermo Fisher Scientific) coupled to an Ultimate 3000 RSLC (Thermo Fisher Scientific). Peptide separation of 1 µl peptide solution (referring to peptides of 1.23*$10^4$ EVs) was accomplished with a solvent gradient from 0 to 100% acetonitrile in 0.1% acetic acid and mass spectrometry data were acquired in data independent mode. Settings including gradient and mass ranges are provided in ***Table S4***.

Data analysis was accomplished using Spectronaut 18.6 (Biognosys). First, a spectral library was created from a sample set containing the mentioned fractions of exophers of the four urine samples by comparing against a Uniprot database limited to human entries (2022). The following parameter settings were employed: tryptic digestion, up to two missed-cleavages allowed, carbamidomethylation at cysteine as fixed and oxidation at methionine as variable condition. The library was then used to compare spectra with each pellet and supernatant fraction per urine sample. In addition, an existing *B. subtilis* $^{15}N$ spectral library was enabled. Detailed search settings are provided as ***Table S4.***

Statistical analysis was performed using R version 4.3.0[23]. The Spectronaut ion report was loaded into R for further processing. The ions were median normalized with respect to the sample-specific *B. subtilis* spike-in signal per sample. Only the normalized ions with a Q-value below 0.001 were utilized for the iBAQ protein intensity calculation. For this calculation, only proteins with 2 or more peptides identified per sample with a Q-value below 0.001 were considered. The resulting iBAQ values were then filtered to retain only unique protein groups.

The final list of proteins per sample, derived from this process, was used for the gProfiler analysis. This analysis was conducted using the gprofiler2[24]package (v0.2.2) using g:SCS multiple testing correction method the significance threshold was set to 0.05. Plots were genrated using tidyverse (v2.0.0)[25], ggtext (v0.1.2)[26], glue (v1.7.0)[27], ggh4x (v0.2.6)[28] packages. The final data with identified proteins and related pathways from Gene Ontology[29,30] can be found as ***Table S4***. Raw data and the spectral library have been deposited to the ProteomeXchange Consortium via the PRIDE[31] partner repository with the dataset identifier PXD049008. The reviewer account details for the PRIDE repository are:

**Username:**
**Password:** pQXHqNCm.

#### Statistical analysis

Results were expressed as mean ± SEM, and significance was set at $*p < 0.05$. The means in a time course were compared using the two-way ANOVA using Prism 8.2.1 for Mac OS X, GraphPad Software, San Diego, California USA, www.graphpad.com. Animal experiments were compared using the one-way ANOVA Kruskal-Wallis test followed by the Dunn's post-test for multiple comparisons using Prism 8.2.1 for Mac OS X, GraphPad Software, San Diego, California USA, www.graphpad.com. Animal experiments with only two groups were compared using the Mann-Whitney U test. Replicates used were biological replicates, which were measured using different samples from distinct experiments. All mice were littermates and were blindly assigned to the experimental groups.

### *Figure S1*

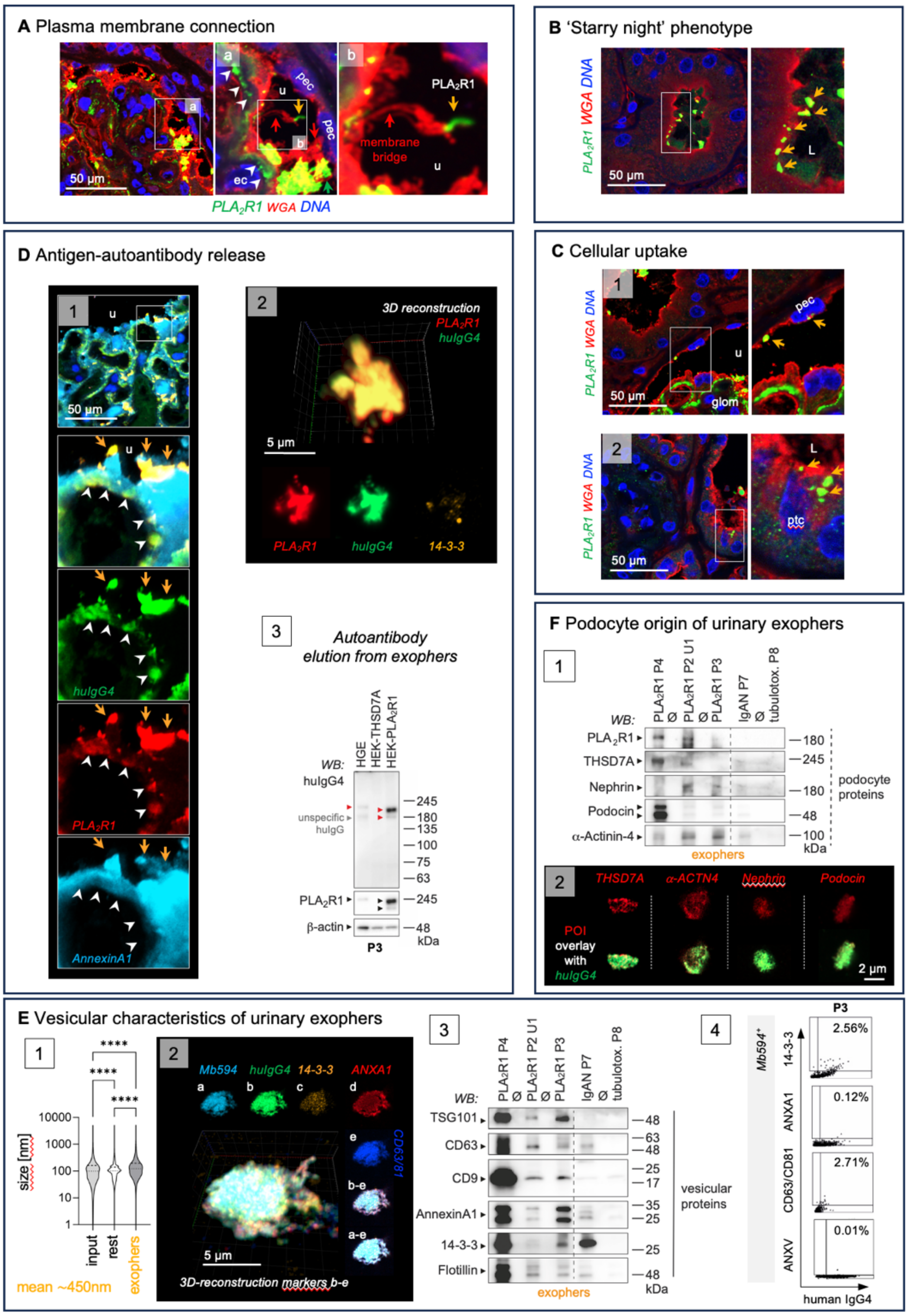

**Exophers are found in nephrotic $PLA_2R1^+$-MN patients P2-P4.** Diagnostic samples of **P2** and **P4** and follow up urine of **P3** $PLA_2R1^+$-MN patients were analyzed for the presence of exophers. If not indicated otherwise, analyses from **P2** urine U1 are shown. **A)** High-resolution confocal micrographs to $PLA_2R1$ (green) and wheat germ agglutinin (red, WGA, binds to N-acetyl-D-glucosamine and sialic acid of the glycocalyx and demarcates plasma membrane), DNA (blue). Glomerular $PLA_2R1$ aggregates localize to the subepithelial space (white arrowheads) and to the urinary (u) space (orange arrows). Urinary space $PLA_2R1$ aggregates exhibit a $WGA^+$ membrane connection (red arrow) to podocytes. **B, C)** High-resolution confocal micrographs to $PLA_2R1$ (green) and WGA (red), DNA (blue) demonstrates $PLA_2R1^+$ aggregates (orange arrows) **B)** within the tubular lumen (L) and **C)** taken up by *panel 1:* parietal epithelial cells (pec) and *panel 2:* proximal tubular cells (ptc). **D)** *Panel 1:* Confocal analyses demonstrate $PLA_2R1$ (red) colocalization with huIgG4 (green) in urinary space aggregates (orange arrows). White arrowheads highlight huIgG4 and PLA$_2$R1 in the subepithelial space. Annexin A1 (light blue) demarcates plasma membrane. *Panel 2:* 3D-reconstructed confocal *z*-stack of an exopher enriched from the patient urine demonstrates the high abundance of PLA$_2$R1 (red) and huIgG4 (green) complexes. Lower row exhibits individual channels in one plane, vesicle marker 14-3-3 (orange). *Panel 3:* Antibodies eluted from **P3** exophers exhibit specific reactivity to PLA$_2$R1 in human glomerular extract (HGE) and in HEK cells transduced with human PLA$_2$R1 but not with human THSD7A. Reprobes to PLA$_2$R1 and β-actin control for molecular weight height and loading. **E**) Vesicular characteristics of PLA$_2$R1$^+$-MN patient **P2** exophers in U1. *Panel 1:* Nanoparticle tracking analyses of vesicular size distribution in total (input) urinary EVs, the exopher-depleted rest fraction and the exopher-enriched fraction. Violin plots indicate median, 25% and 75% percentile. Mean size of PLA$_2$R1$^+$-MN exophers is ~450 nm. *Panel 2:* 3D-reconstructed confocal *z*-stack of an exopher demonstrates expression of the vesicle markers 14-3-3 (orange), Annexin A1 (red), CD63/CD81 (blue). MemBrite594 (light blue) stains the plasma membrane, huIgG4 (green) the bound autoantibody. *a-e:* individual channels in one plane. *Panel 3:* Immunoblot to vesicle markers in enriched exophers isolated from the diagnostic urines of the 3 PLA$_2$R1$^+$-MN patients in comparison to the nephrotic IgAN patient **P7** and the tubulo-toxic kidney injury patient **P8**. *Panel 4:* Image stream analysis of vesicle marker distribution on MemBrite594$^+$/huIgG4$^+$ exophers discerns 14-3-3 as an abundant marker in PLA$_2$R1$^+$-MN exophers collected from the urine of **P3**. Note the lack of Annexin V signal (marker for apoptotic bodies). **F**) Urinary exophers contain podocyte proteins. *Panel 1:* Immunoblot detection of podocyte-specific proteins in enriched exophers isolated from diagnostic urines of the PLA$_2$R1$^+$-MN patients in comparison to the two nephrotic patients diagnosed with IgAN (**P7**) or tubulo-toxic kidney injury (**P8**). *Panel 2:* Confocal analyses demonstrate expression of podocyte-specific proteins (protein of interest (POI) in red) within huIgG4$^+$ (green) exophers of **P2**.

### *Figure S2*

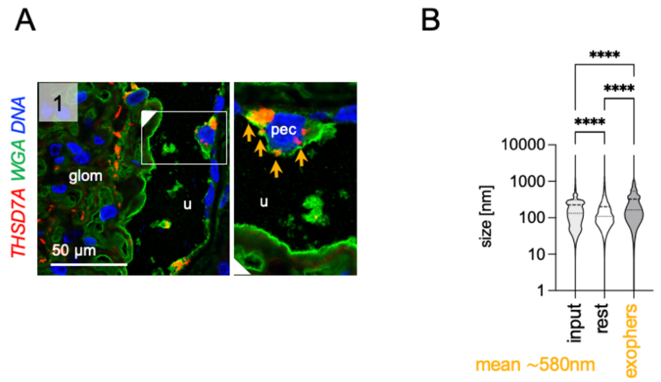

**Exophers are found in a THSD7A$^+$-MN patient P1. A**) High-resolution confocal micrographs to THSD7A (red) and wheat germ agglutinin (green, WGA, binds to N-acetyl-D-glucosamine and sialic acid of the glycocalyx and demarcates plasma membrane), DNA (blue) of the diagnostic biopsy. Glomerular THSD7A aggregates (orange arrows) localize to the cytoplasm of parietal epithelial cells (pec) demonstrating uptake, u = urinary space, glom = glomerular convolute. **B**) Nanoparticle tracking analyses of vesicular size distribution in total (input) urinary EVs, the exopher-depleted rest fraction and the exopher-enriched fraction of the diagnostic urine. Violin plots indicate median, 25% and 75% percentile. Mean size of THSD7A$^+$-MN exophers is ~580 nm. The upper range of the NTA measurements is technically limited to 5000 nm.

### *Figure S3*

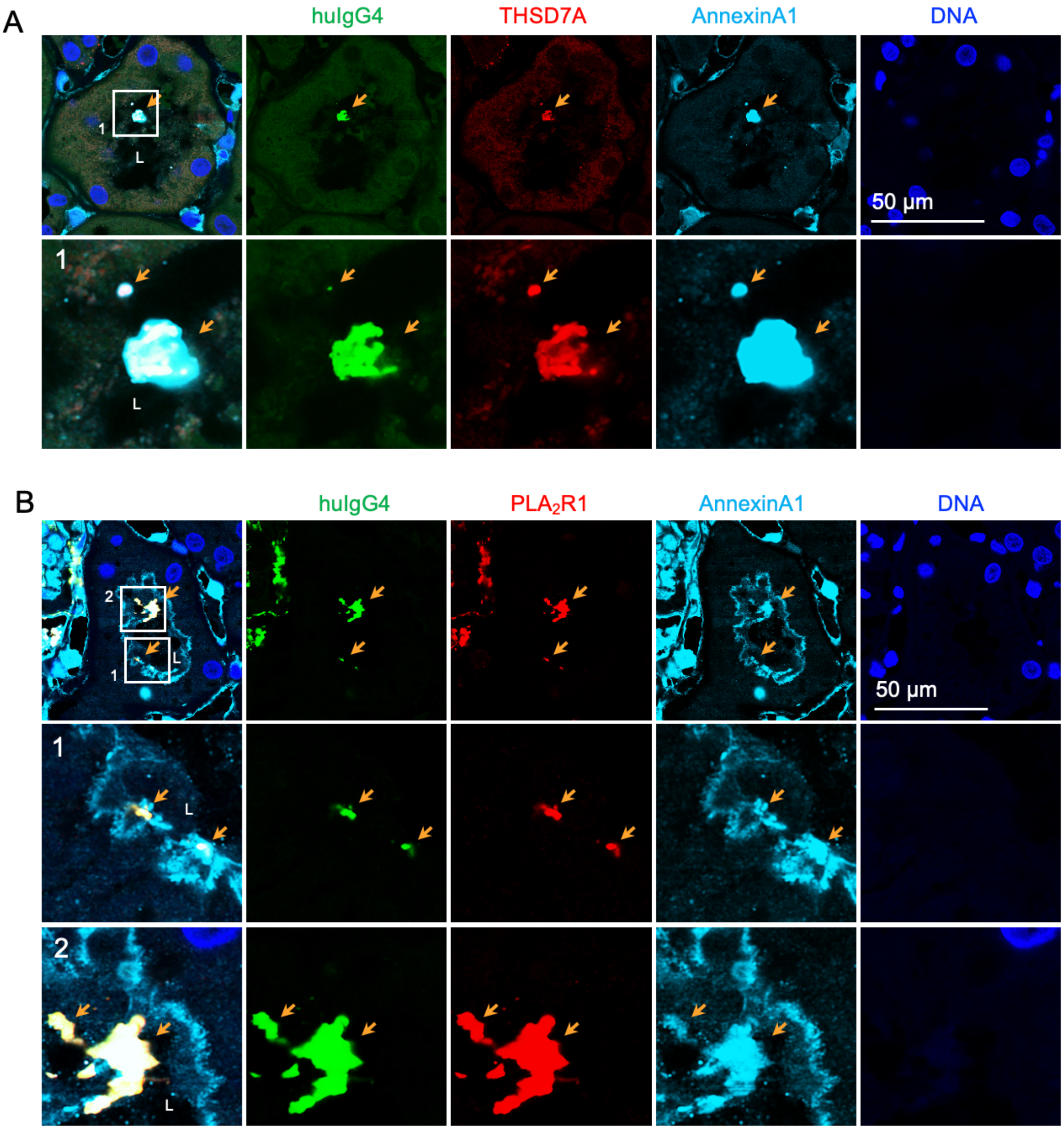

**Antigen/autoantibody complexes are shed to the urine and found in adherence to the tubular apical membrane in membranous nephropathy patients P1 and P2.** Diagnostic biopsies from a A) $THSD7A^+$-MN patient **P1** and B) a $PLA_2R1^+$-MN patient **P2** were stained for human IgG4 (green) to detect the autoantibodies and the respective antigens THSD7A or $PLA_2R1$ (both red). Annexin A1 (light blue) demarcates the vesicular origin of the huIgG4/MN-antigen containing immune complexes (orange arrow), DNA (blue) was stained with Hoechst. L = tubular lumen.

### *Figure S4*

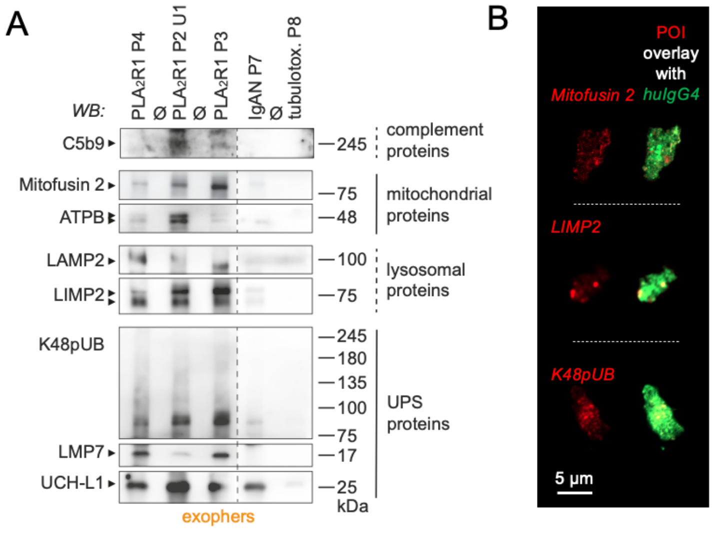

**Urinary exophers enriched from the diagnostic urines of $PLA_2R1^+$-MN patients P2-P4 contain disease-associated proteins.** Urinary exophers enriched from diagnostic urine samples of **P2**, **P3**, and **P4** $PLA_2R1^+$-MN patients were analyzed for the presence of disease-associated proteins. **A)** Immunoblot detection of disease associated proteins belonging to the complement cascade (C5b9, membrane attack complex), mitochondria (membrane proteins Mitofusin 2 and ATP synthase ATPB), lysosomes (membrane proteins LAMP2 and LIMP2), and the ubiquitin proteasome system (UPS: K48-polyubiquitinated proteins, the proteolytic β-subunit LMP7, the deubiquitinating enzyme UCH-L1) in enriched exophers from the diagnostic urines of the 3 $PLA_2R1^+$-MN patients in comparison to the nephrotic IgAN patient **P7** and the tubulo-toxic kidney injury patient **P8**. **B)** Confocal analyses demonstrate expression of Mitofusin 2, LIMP2, or K48pUB (all proteins of interest (POI) in red) within urinary $huIgG4^+$ (green) exophers of U1 of the $PLA_2R1^+$-MN patient **P2**.

### *Figure S5*

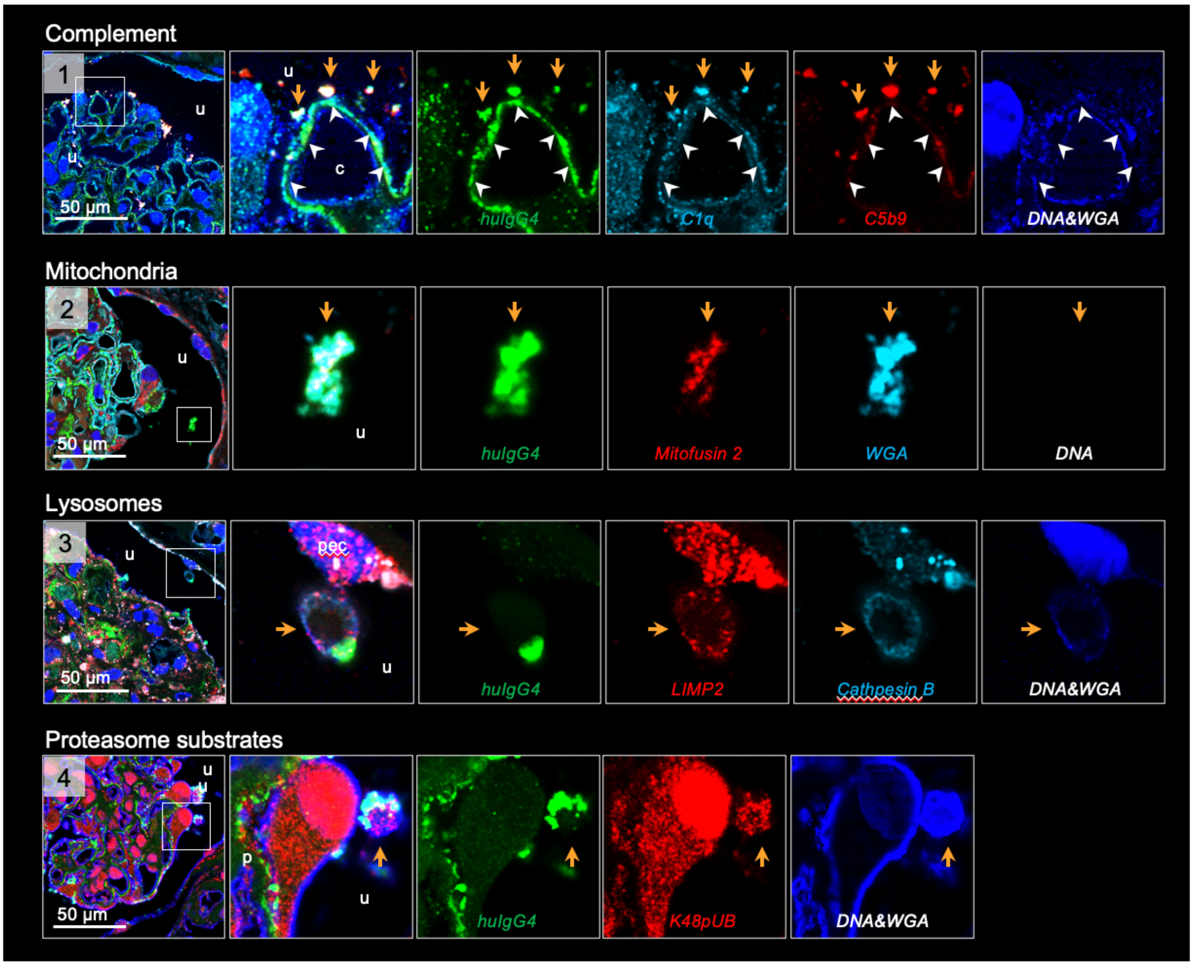

**Glomerular huIgG4$^+$ aggregates within the urinary space contain disease-associated proteins in the diagnostic biopsy of the THSD7A$^+$-MN patient P1.** High-resolution confocal micrographs to huIgG4 (green) of the diagnostic biopsy demarcates glomerular aggregates, urinary space aggregates are highlighted by an orange arrow, the white arrowheads highlight the podocyte subepithelial space. The overview micrographs indicate the urinary space localization of the magnified emboxed glomerular huIgG4$^+$ aggregates. Disease associated proteins were colocalized to huIgG4$^+$ aggregates: *Panel 1:* complement proteins C1q (light blue) and C5b9 (membrane attack complex; red). Blue: DNA and wheat germ agglutinin (WGA) to mark the cell nucleus and plasma membrane. WGA binds to N-acetyl-D-glucosamine and sialic acid of the glycocalyx. *Panel 2:* the mitochondrial membrane protein Mitofusin 2 (red). *Panel 3:* the lysosomal membrane protein LIMP2 (red) and the lysosomal enzyme Cathepsin B (light blue). *Panel 4:* the proteasome substrates K48-polyubiquitinated proteins (red). Note the presence of complement proteins, mitochondrial, lysosomal, and proteasome substrates within huIgG4$^+$ urinary aggregates, classifying them as exophers in this patient.

### *Figure S6*

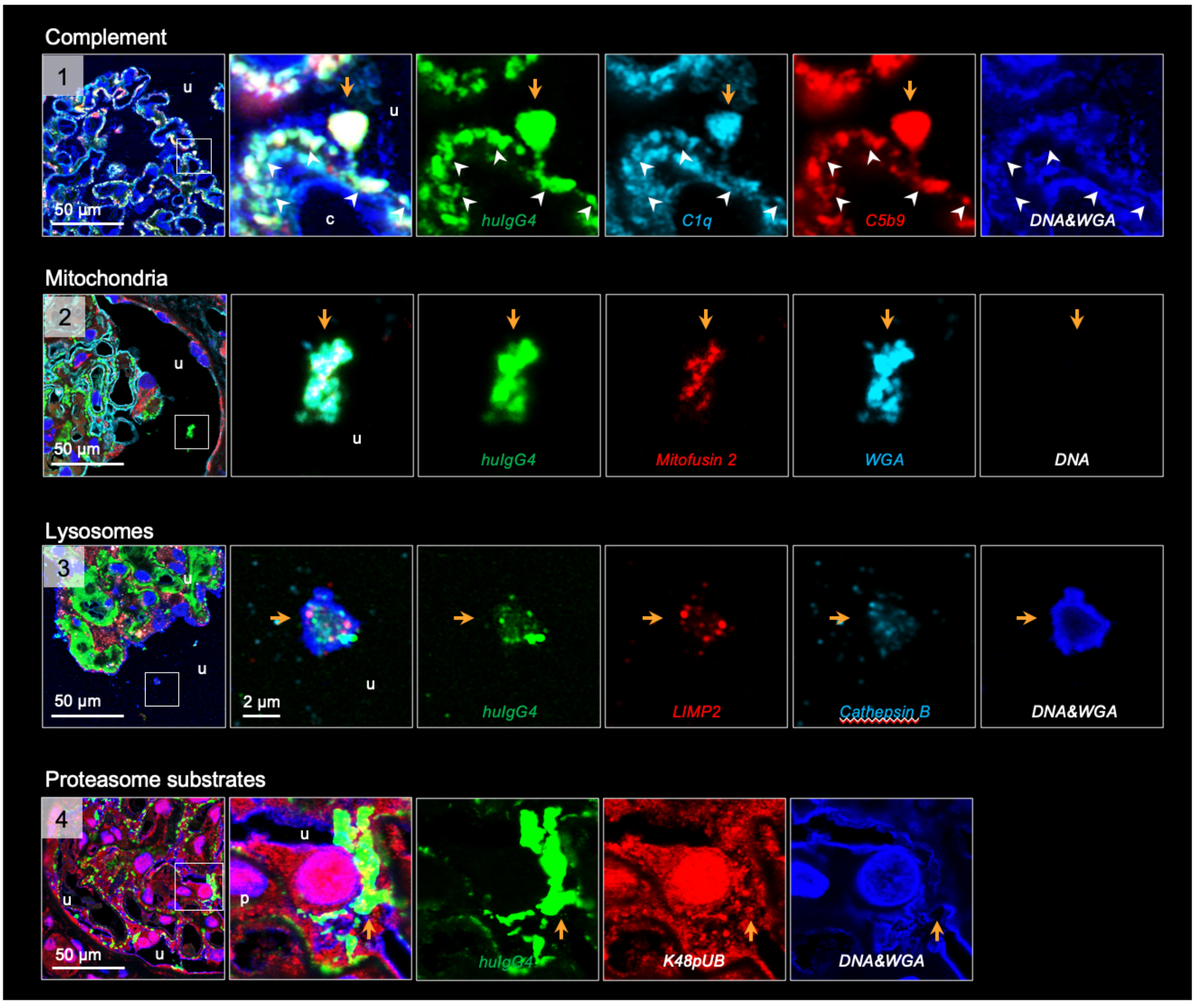

**Glomerular huIgG4⁺ aggregates within the urinary space contain disease-associated proteins in the diagnostic biopsy of the $PLA_2R1^+$-MN patient P2.** High-resolution confocal micrographs to huIgG4 (green) of the diagnostic biopsy demarcates glomerular aggregates, urinary space aggregates are highlighted by an orange arrow, the white arrowheads highlight the podocyte subepithelial space. The overview micrographs indicate the urinary space localization of the magnified emboxed glomerular huIgG4$^+$ aggregates. Disease associated proteins were colocalized to huIgG4+ aggregates: *Panel 1:* complement proteins C1q (light blue) and C5b9 (membrane attack complex; red). Blue: DNA and wheat germ agglutinin (WGA) to mark the cell nucleus and plasma membrane. WGA binds to N-acetyl-D-glucosamine and sialic acid of the glycocalyx. *Panel 2:* the mitochondrial membrane protein Mitofusin 2 (red). *Panel 3:* the lysosomal membrane protein LIMP2 (red) and the lysosomal enzyme Cathepsin B (light blue). *Panel 4:* the proteasome substrates K48-polyubiquitinated proteins (red). Note the presence of complement proteins, mitochondrial, lysosomal, and proteasome substrates within huIgG4$^+$ urinary aggregates, classifying them as exophers in this patient.

### *Figure S7*

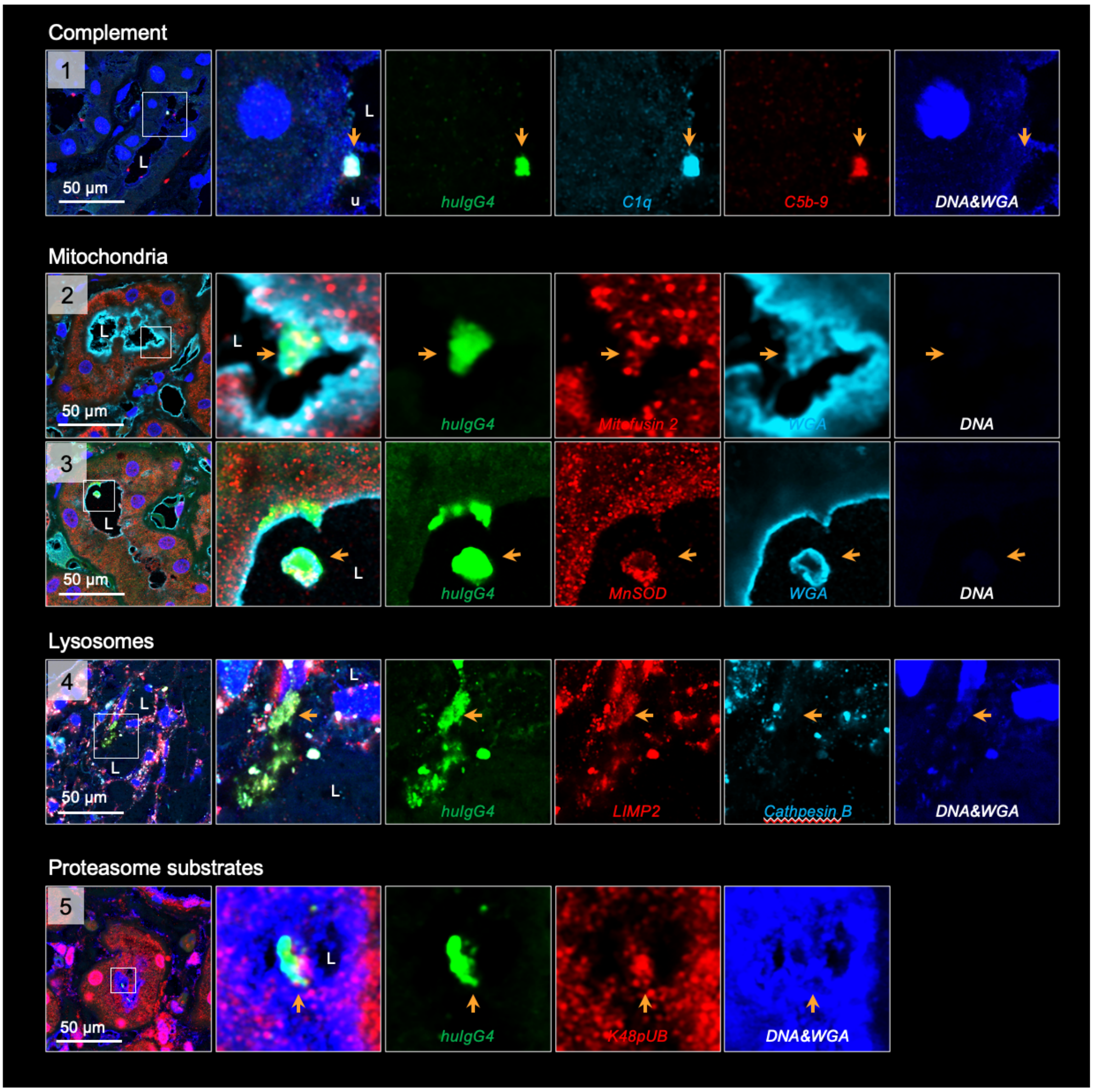

**HuIgG4$^{+}$ aggregates within the tubular lumen contain disease-associated proteins in the diagnostic biopsy of the THSD7A$^{+}$-MN patient P1.** High-resolution confocal micrographs to huIgG4 (green) of the diagnostic biopsy demarcates released glomerular aggregates (highlighted by an orange arrow) present within the tubular lumen (L). The overview micrographs indicate the urinary space localization of the magnified emboxed glomerular huIgG4$^{+}$ aggregates. Disease associated proteins were colocalized to huIgG4$^{+}$ aggregates: *Panel 1:* complement proteins C1q (light blue) and C5b9 (membrane attack complex, red). Blue: DNA and wheat germ agglutinin (WGA) to mark the cell nucleus and plasma membrane. WGA binds to N-acetyl-D-glucosamine and sialic acid of the glycocalyx. *Panels 2, 3:* the mitochondrial membrane protein Mitofusin 2 (red) or MnSOD (red). *Panel 4:* the lysosomal membrane protein LIMP2 (red) and the lysosomal enzyme Cathepsin B (light blue). *Panel 5:* the proteasome substrates K48-polyubiquitinated proteins (red). Note the presence of complement proteins, mitochondrial, lysosomal, and proteasome substrates within huIgG4$^{+}$ urinary aggregates, classifying them as released exophers in this patient.

### *Figure S8*

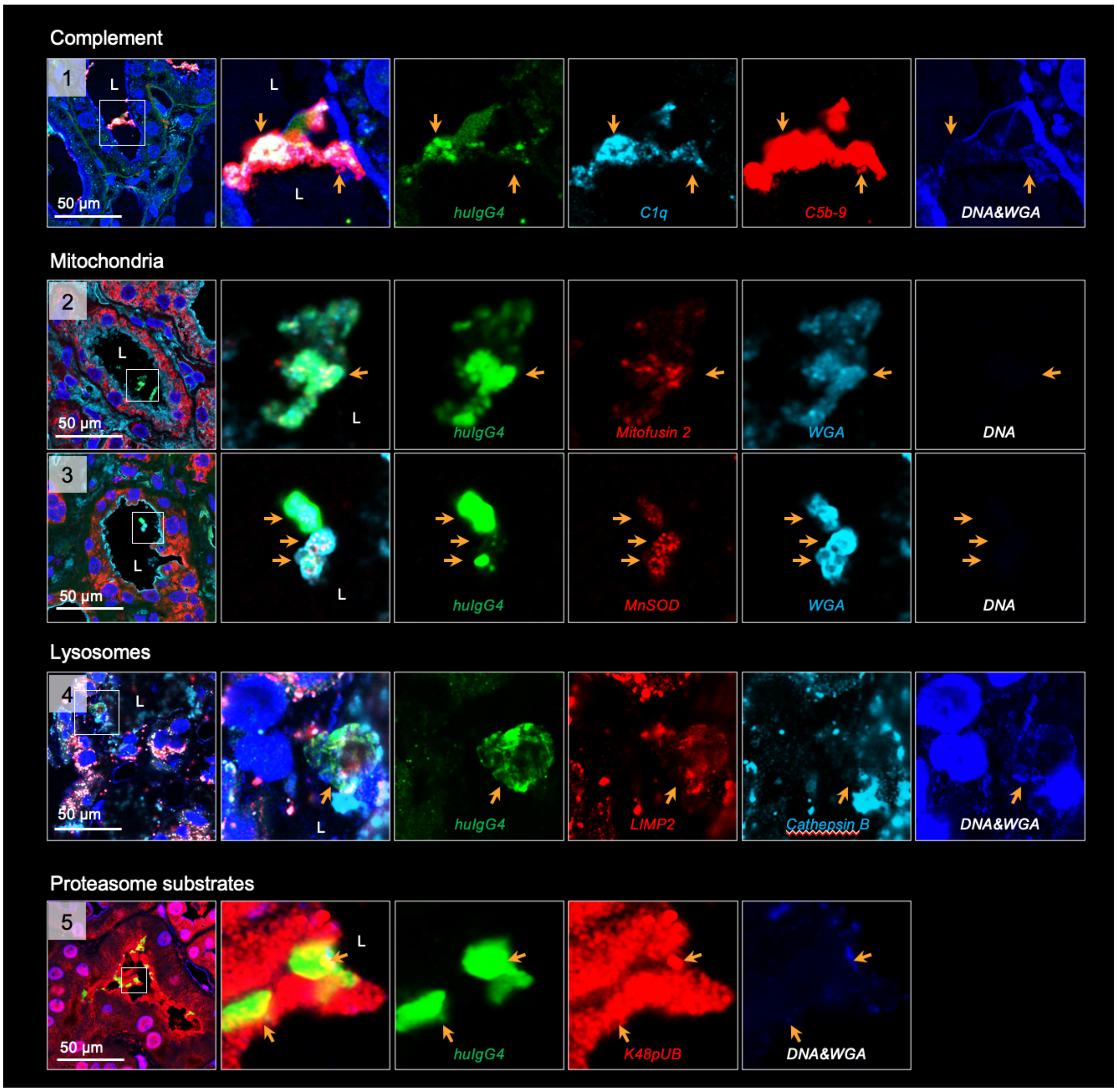

**HuIgG4[+] aggregates within the tubular lumen contain disease-associated proteins in the diagnostic biopsy of the $PLA_2R1^+$-MN patient P2.** High-resolution confocal micrographs to huIgG4 (green) of the diagnostic biopsy demarcates released glomerular aggregates (highlighted by an orange arrow) present within the tubular lumen (L). The overview micrographs indicate the urinary space localization of the magnified emboxed glomerular huIgG4[+] aggregates. Disease associated proteins were colocalized to huIgG4[+] aggregates: *Panel 1:* complement proteins C1q (light blue) and C5b9 (membrane attack complex; red). Blue: DNA and wheat germ agglutinin (WGA) to mark the cell nucleus and plasma membrane. WGA binds to N-acetyl-D-glucosamine and sialic acid of the glycocalyx. *Panels 2, 3:* the mitochondrial membrane protein Mitofusin 2 (red) or MnSOD (red). *Panel 4:* the lysosomal membrane protein LIMP2 (red) and the lysosomal enzyme Cathepsin B (light blue). *Panel 5:* the proteasome substrates K48-polyubiquitinated proteins (red). Note the presence of complement proteins, mitochondrial, lysosomal, and proteasome substrates within huIgG4[+] urinary aggregates, classifying them as released exophers in this patient.

### *Figure S9*

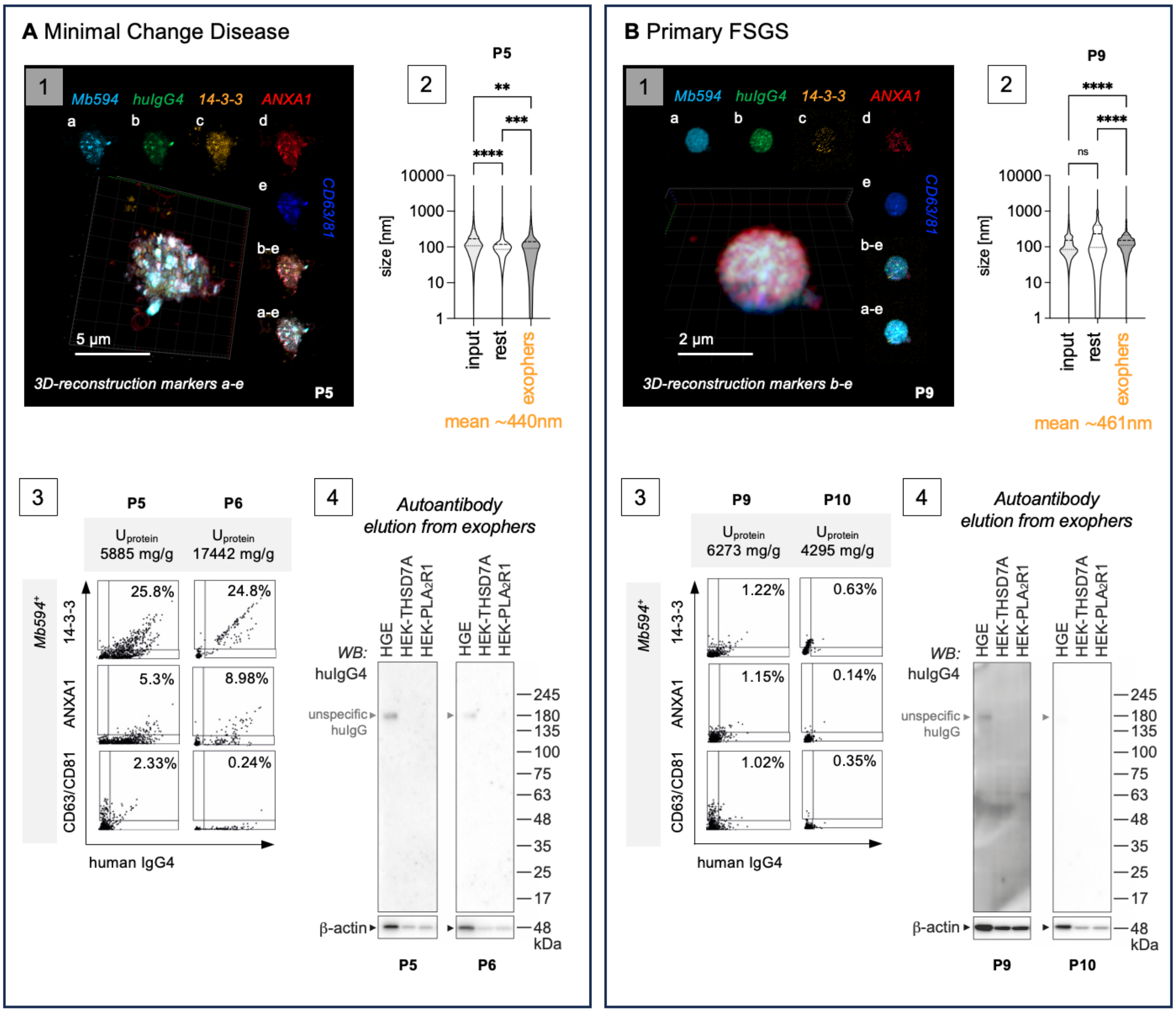

**Characteristics of the huIgG4-enriched urinary EV fractions from MCD patients P5, P6 and primary FSGS patients P9, P10 in diagnostic urines.** Urinary EVs enriched *via* huIgG4-pulldown from **A**) the diagnostic nephrotic urine of minimal change patients **P5** and **P6** or from **B**) nephrotic urine of primary FSGS patients **P9** (diagnostic urine) and **P10** (follow up urine one month after diagnosis) were characterized. *Panels 1:* 3D-reconstructed confocal *z*-stack of an exopher demonstrates expression of the vesicle markers 14-3-3 (orange), Annexin A1 (red), CD63/CD81 (blue). MemBrite594 (light blue) stains the plasma membrane, huIgG4 (green) the bound antibody. *a-e:* individual channels in one plane. *Panels 2:* Nanoparticle tracking analyses of vesicular size distribution in total (input) urinary EVs, the exopher-depleted rest fraction and the exopher-enriched fraction. Violin plots indicate median, 25% and 75% percentile. ***p* <0.01, ****p* <0.001, *****p* <0.0001, One-way ANOVA. Mean size of the huIgG4$^{+}$-enriched EVs is indicated in orange. *Panels 3:* Image stream quantification of vesicle marker distribution on MemBrite594$^{+}$huIgG4$^{+}$ EVs discerns 14-3-3 as an abundant marker in the two MCD patients but not in the two primary FSGS patients. The concomitant levels of proteinuria of the analyzes diagnostic urines are indicated in mg protein / g creatinine. *Panels 4:* Antibodies eluted from patient exophers show no reactivity to proteins present in human glomerular extract (HGE) or in HEK cells transduced with PLA$_2$R1 or with THSD7A.

### *Figure S10*

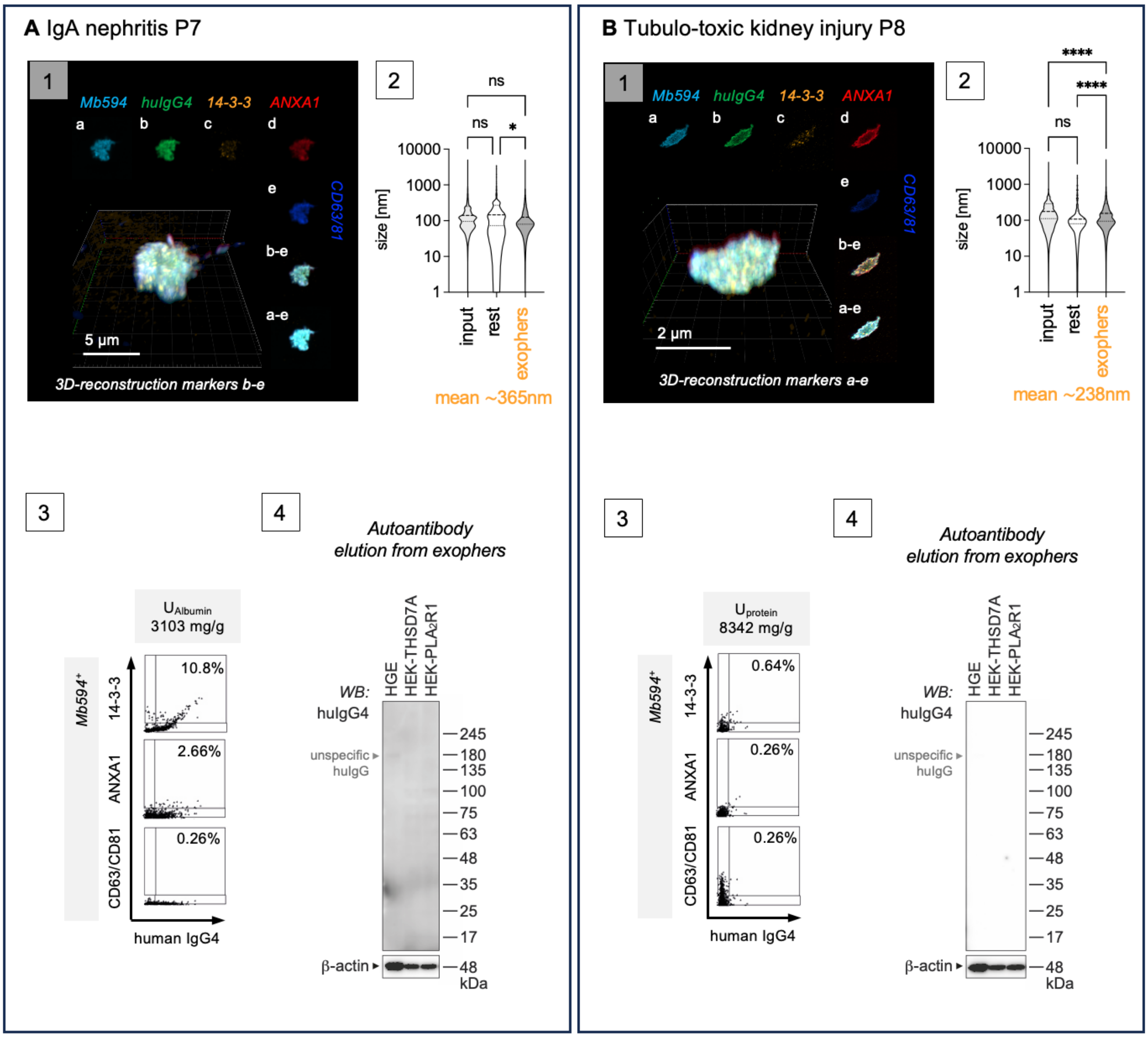

**Characteristics of the huIgG4-enriched urinary EV fractions from nephrotic patients with IgA nephritis P7 or tubulo-toxic kidney injury P8 in diagnostic urines.** Urinary EVs enriched *via* huIgG4-pulldown from the diagnostic nephrotic urine of A) a patient diagnosed with IgA nephritis **P7** or from B) a patient diagnosed with tubulo-toxic kidney injury **P8** were characterized. *Panels 1:* 3D-reconstructed confocal *z*-stack of an exopher demonstrates expression of the vesicle markers 14-3-3 (orange), Annexin A1 (red), CD63/CD81 (blue). MemBrite594 (light blue) stains the plasma membrane, huIgG4 (green) the bound antibody. *a-e:* individual channels in one plane. *Panels 2:* Nanoparticle tracking analyses of vesicular size distribution in total (input) urinary EVs, the exopher-depleted rest fraction and the exophers enriched fraction. Violin plots indicate median, 25% and 75% percentile. **p* <0.05, *****p* <0.0001, ns = not significant, One-way ANOVA. Mean size of the huIgG4$^+$-enriched EVs is indicated in orange. *Panels 3:* Image stream quantification of vesicle marker distribution on MemBrite594$^+$huIgG4$^+$ EVs discerns 14-3-3 as an abundant marker in the IgAN patient. The concomitant levels of proteinuria of the analyzes diagnostic urines are indicated in mg protein / g creatinine. *Panels 4:* Antibodies eluted from patient exophers show no reactivity to proteins present in human glomerular extract (HGE) or in HEK cells transduced with $PLA_2R1$ or with THSD7A.

### *Figure S11*

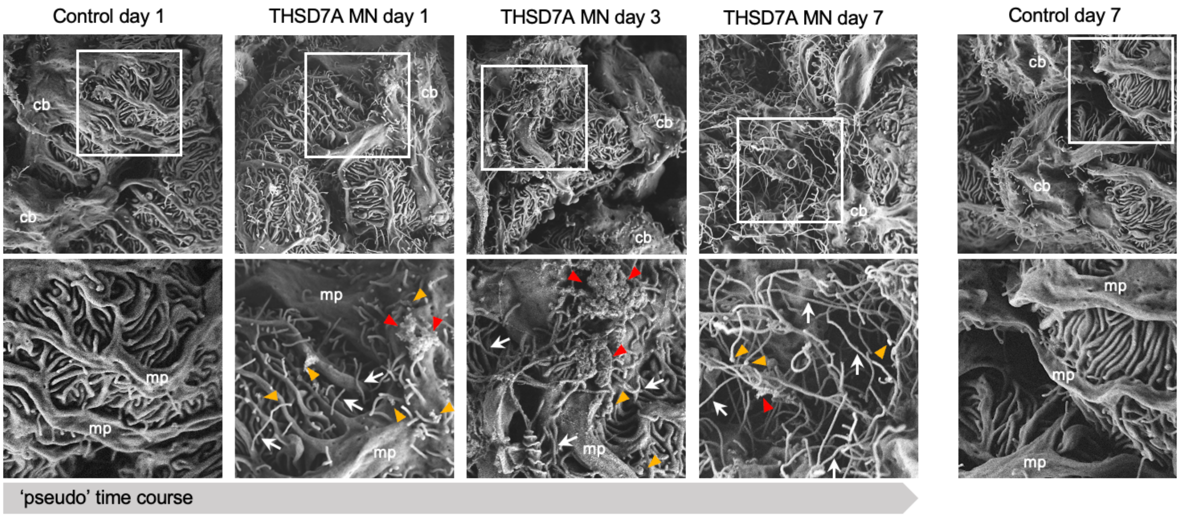

**THSD7A autoantibody binding induces the formation of long membrane extensions with distal bulges from podocytes in experimental THSD7A$^{+}$-MN.** THSD7A$^{+}$-MN was induced in BALB/c mice by injection of rabbit (rb)THSD7A-antibodies (abs) or control rbIgG (ctrl-abs), kidneys were collected on day 1, 3 and 7 fixed by *in vivo* perfusion using an established protocol to "freeze" membrane-processes to enable early and late ultrastructural analyses such as scanning electron microscopic (SEM) evaluations of podocyte morphology. Note the beginning formation of membrane extensions (white arrows) with distal bulges (orange arrowheads) as early as day 1 in the rbTHSD7A-abs treated mouse which originate from foot processes, major processes (mp) as well as the cell body (cb). These extensions and distal bulges are more frequent and longer on day 3 and extensive in day 7 rbTHSD7A-abs treated mouse. Large protrusions with rough surface are focally observed (red arrowheads) at the urinary side of podocyte processes and cell body. Control mice exhibit shorter membrane extensions arising predominantly from the cell body.

### *Figure S12*

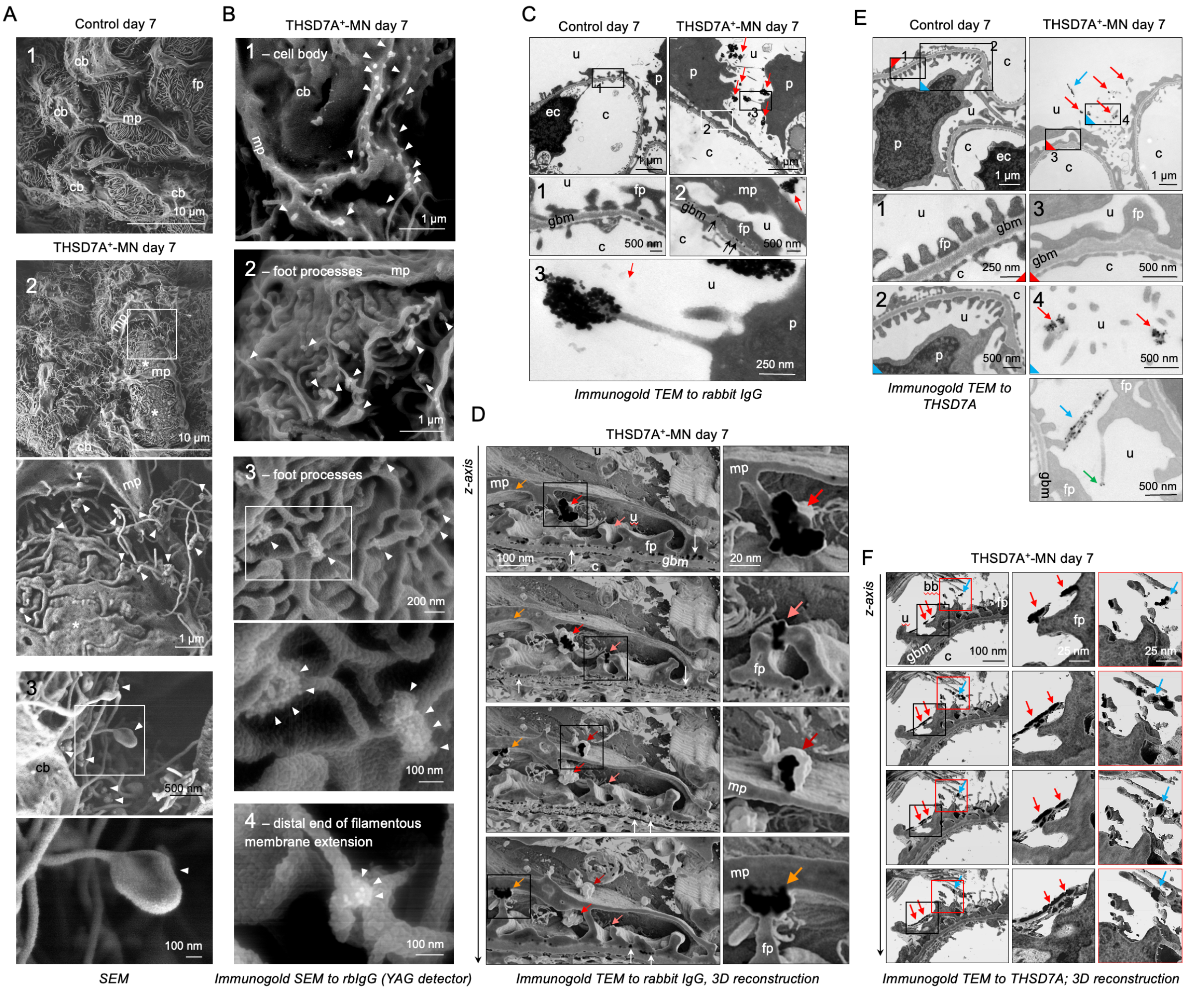

**THSD7A autoantibody binding induces exopher-formation in podocytes experimental THSD7A$^+$-MN.** THSD7A$^+$-MN was induced in BALB/c mice by injection of rbTHSD7A-abs or control rbIgG (ctrl-abs), kidneys were collected on day 1 and on day 7 fixed by *in vivo* perfusion using an established protocol to "freeze" membrane-processes. **A)** Scanning electron microscopic (SEM) (*panel 1*) overview of podocytes from a control glomerulus and (*panel 2*) of a THSD7A$^+$-MN podocyte day 7. Boxed areas depict close-ups; arrowheads point to long membrane extensions with distal bulges, which originate from major (mp) and foot processes (fp) as well as the cell body (cb) of podocytes; asterisks = area of foot process effacement. **B-D)** Pre-embedding immunogold EM evaluation of bound rbTHSD7A-abs was performed with a gold-labeled anti-rabbit IgG. **B)** Gold aggregates (arrowheads) were detected using the YAG laser in SEM. **C)** Transmission EM (TEM) demonstrates gold aggregates to rbIgG within the urinary space connected to the podocyte cell body by long membrane extensions (red arrows); u = urinary space, c = capillary lumen, gbm = glomerular basement membrane, p = podocyte nucleus, ec = endothelial cell nucleus; fp = foot process. Close-ups in boxed areas depict subepithelial gold deposits (black arrows) and urinary gold aggregates (red arrows). **D)** 3-dimensional reconstruction of 48 consecutive TEM micrographs demonstrates the budding-off of gold-labeled structures (colored arrows highlight one individual gold structure from plane to plane) from podocyte foot and major processes. Gold appears black in the plane of cutting. White arrows demarcate subepithelial gold. **E-F)** Pre-embedding immunogold EM evaluations of THSD7A was performed with an affinity-purified patient anti-THSD7A autoantibody as primary and a gold-labeled anti-human IgG as secondary antibody, to circumvent unspecific signals from intrinsic mouse and rabbit IgG. **E)** TEM depicts gold-labeled THSD7A aggregates in the urinary space (red arrows) and as along a filamentous structure (blue arrows) and at the distal end of a long membrane extension (green arrow). **F)** 13 consecutive TEM micrographs were reconstructed 3-dimensionally to demonstrate the budding-off of gold labeled THSD7A structures from podocyte foot processes as well as gold labeled long membrane structures that are derived from foot processes (red arrows). Note the adherence (connection) of gold-labeled THSD7A long membrane structures to the proximal tubular brush-border at the urinary pole of glomeruli (blue arrow).

### *Figure S13*

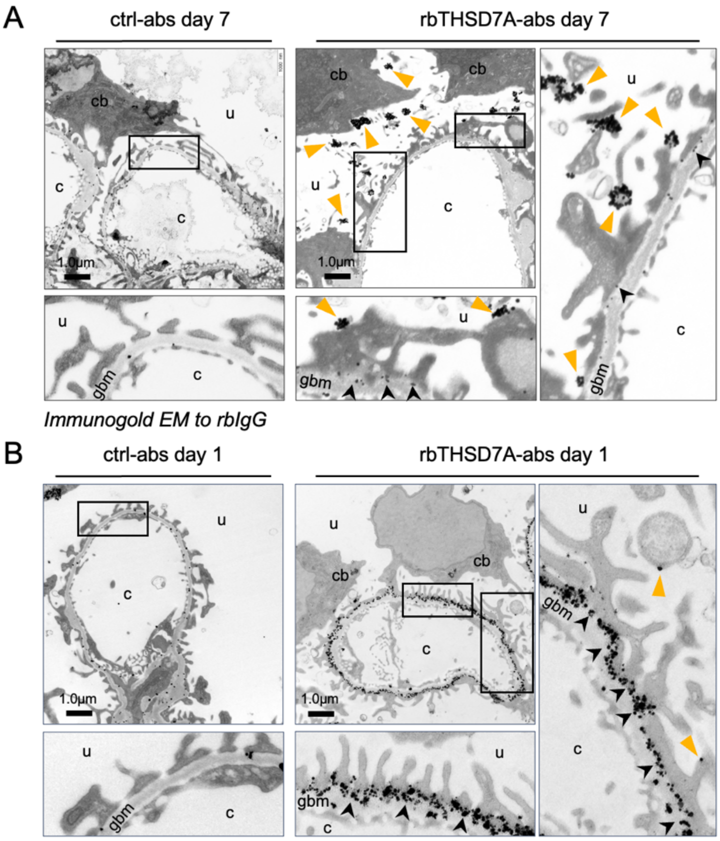

*Immunogold EM to rbIgG*

**Immunogold EM localization of injected rbTHSD7A-abs day 1 and day 7.** THSD7A$^{+}$-MN was induced in BALB/c mice by injection of rabbit (rb)THSD7A-antibodies (abs) or control rbIgG (ctrl-abs), kidneys were collected on **A**) day 7 and on **B**) day 1 fixed by *in vivo* perfusion using an established protocol to "freeze" membrane-processes to enable early and late ultrastructural analyses such as immunogold EM to the bound rabbit THSD7A antibodies. Note the almost exclusive localization of the rabbit IgG in the subepithelial space (black arrowheads) on day 1. No gold is found within podocytes. Some discreet gold particles are found on the apical side of foot processes (orange arrowhead). The control rabbit IgG injected mouse shows unspecifically localized rbIgG at the glomerular basement membrane (GBM). On day 7 rbIgG is sparse in the subepithelial space. In the urinary space, gold accumulations are appreciated at the end of membrane extensions that arise from foot processes, major processes and from the podocyte cell body; cb = cell body; u = urinary space; c = capillary space; gbm = glomerular basement membrane.

### *Figure S14*

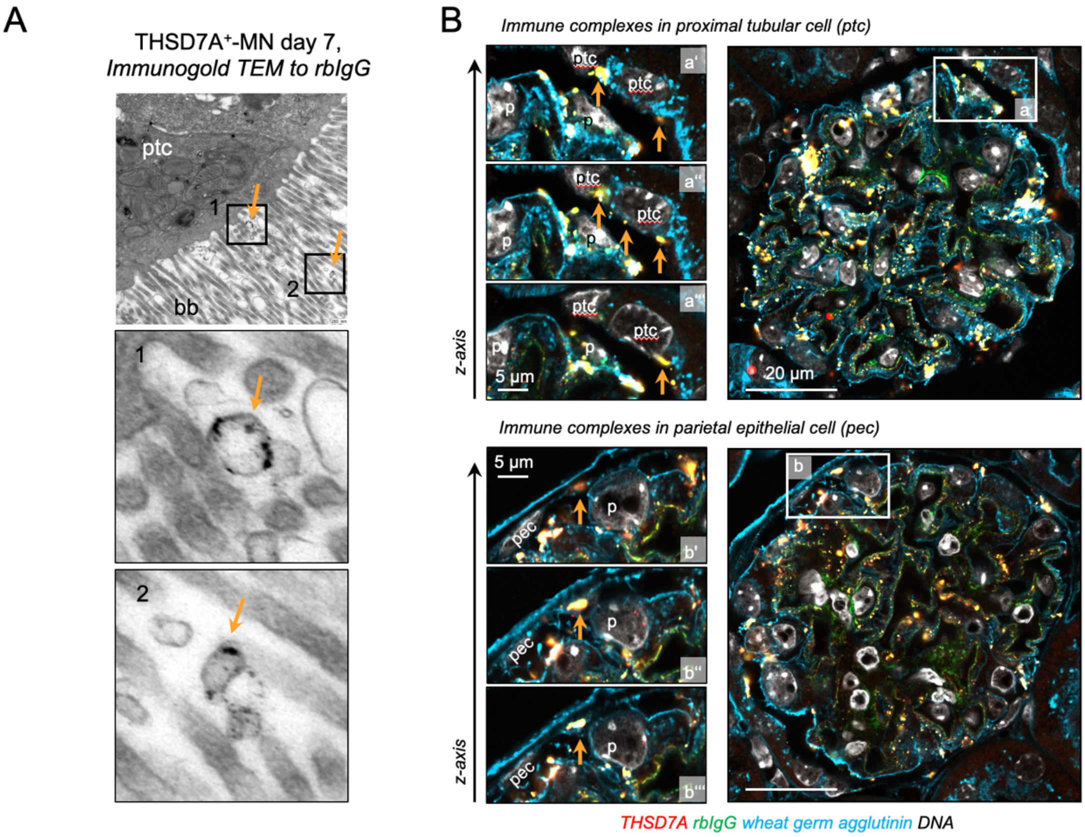

**Urinary exopher-release and uptake in experimental THSD7A$^{+}$-MN.** THSD7A$^{+}$-MN was induced in BALB/c mice by injection of rbTHSD7A-abs or ctrl-abs. **A)** Transmission electron microscopical detection of gold-labeled rbIgG-containing vesicular structures (orange arrows) within the urinary space and at the brush border (bb) of a proximal tubular cell (ptc) of the THSD7A$^{+}$-MN mouse on day 7. **B)** Confocal *z*-stacks of experimental kidneys day 12 stained for THSD7A (red) and rbIgG (green) demonstrate the glomerular accumulation of immune complexes at proximal tubular cells (ptc) of the glomerular urinary pole and in parietal epithelial cells (pec) of Bowman's capsule (orange arrows); p = podocyte. Three different confocal planes in the *z*-axis are shown. Wheat germ agglutinin (WGA, light blue) demarcates the podocyte plasma membrane, as WGA binds to N-acetyl-D-glucosamine and sialic acid of the glycocalyx, Hoechst (DNA, white).

### *Figure S15*

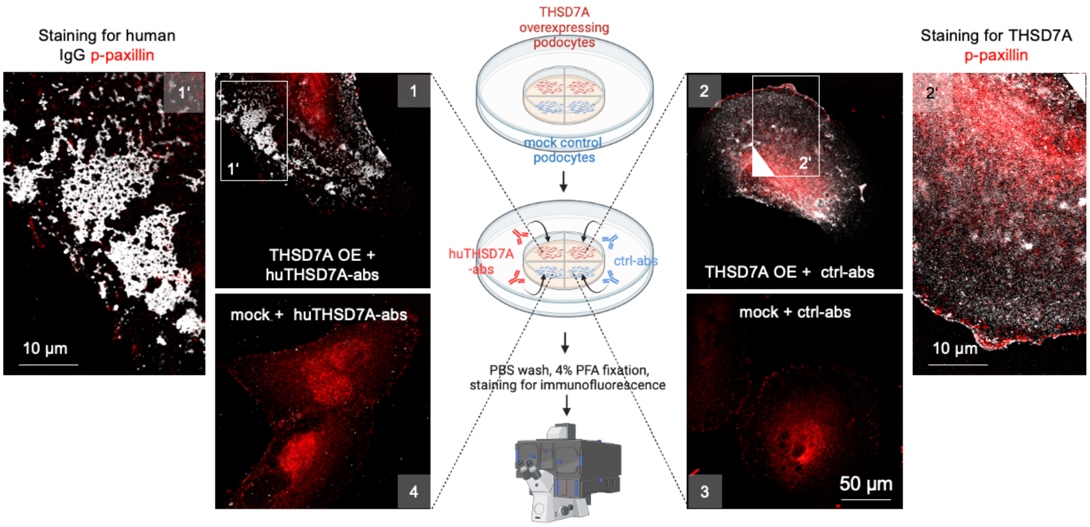

***In vitro* exposure to human THSD7A-abs results in crosslinking of THSD7A.** Human podocytes overexpressing human THSD7A, or mock control cells were treated in culture with affinity-purified patient THSD7A-autoantibodies (huTHSD7A-abs) or unspecific control human IgG (ctrl-abs). *Scheme* in the center summarizes the experimental setup of *in vitro* imaging of all 4 conditions simultaneously. Following *in vitro* exposure to antibodies for 1 hour, cells were washed and fixed. Staining was performed as follows: *Condition 1: in vitro* binding of huTHSD7A-abs to THSD7A was detected by staining with FITC α-huIgG showing changes of THSD7A localization when the autoantibody is bound. *Condition 2:* normal THSD7A expression was visualized by staining with huTHSD7A-abs followed by FITC α-huIgG. As ctrl-abs do not bind to THSD7A, this approach shows the normal distribution of THSD7A without bound autoantibody. *Condition 3:* to check for THSD7A expression in mock cells, cells were stained with huTHSD7A-abs followed by FITC α-huIgG. *Conditions 4:* To check for unspecific binding of huTHSD7A-abs *in vitro*, mock cells were stained with FITC α-huIgG. *Confocal micrographs* demonstrate that exposure to huTHSD7A-abs *in vitro* for 1-hour results in crosslinking of THSD7A (white) at the luminal side of podocytes (*panel 1*). Note the honey-comb pattern of huIgG patches in the huTHSD7A-abs treated podocyte. Phospho-paxillin (p-paxillin, red) was used as counterstain to visualize focal adhesions. *Panel 2* demonstrates unaltered expression of THSD7A at the membrane of THSD7A overexpressing podocytes (THSD7A OE). *Panels 3* and *4* demonstrate specificity of the experimental approach as no THSD7A expression is found in mock podocytes (*panel 3*) and no unspecific binding in of huTHSD7A-abs is present in mock podocytes (*panel 4).*

### *Figure S16*

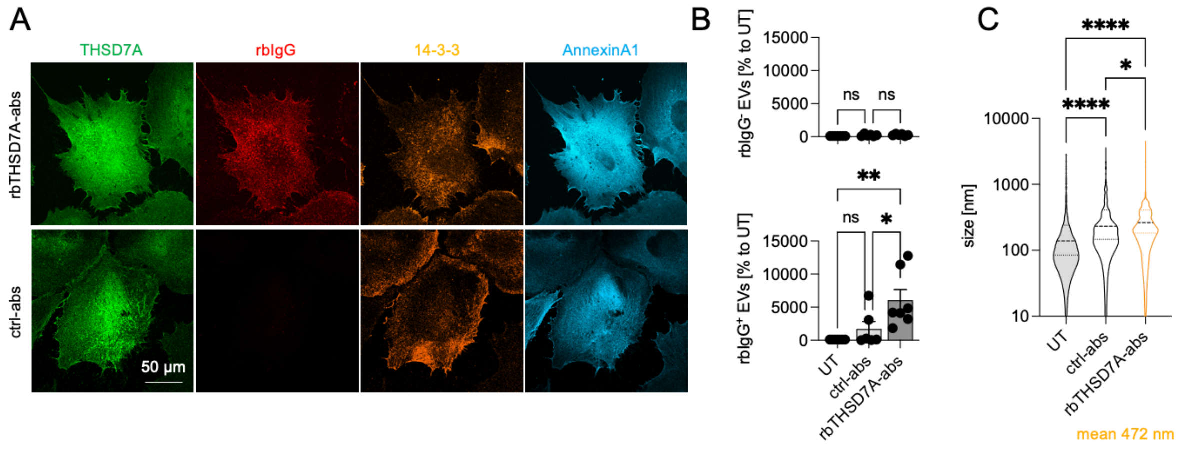

***In vitro* exposure of human podocytes to rbTHSD7A-abs induces release of large rbIgG⁺ EVs.** Human podocytes overexpressing human THSD7A were treated in culture with rbTHSD7A-abs or unspecific control rbIgG (ctrl-abs) for 6h. Cells were fixed, and medium was collected. **A)** Confocal images depicting specific binding of rbTHSD7A-abs (red) to THSD7A (green). Annexin A1 (light blue) and 14-3-3 (orange) were stained to identify areas of exopher-genesis. **B)** Quantification of % EVs without bound rbIgG (upper graph, MemBrite594$^+$/rbIgG$^-$) and with bound rbIgG (lower graph MemBrite594$^+$/rbIgG$^+$, exopher-enriched EVs) released within 6 hours to 3 ml exosome-depleted medium after ctrl-abs or rbTSHD7A-abs exposure in comparison to untreated (UT) hu-podocytes. Graphs depict mean ± SEM, pooled data from 5-6 independent experiments, $N$ = 5-6, **$p \leq 0.01$ to UT, *$p \leq 0.05$ to ctrl-abs, ns = not significant, One-way ANOVA with Dunn's multiple comparison. **C)** Size distribution of EVs enriched from the culture medium of untreated podocytes (UT) or after 6 h of exposure to ctrl-abs or rbTHSD7A-abs. Pooled data from 5 independent experiments with N = 1 per condition, **$p < 0.01$. ****$p < 0.0001$, One way ANOVA. Violin plots indicate median, 25% and 75% percentile, the mean size of EVs released from rbTHSD7A-ab exposed podocytes is indicated.

### *Figure S17*

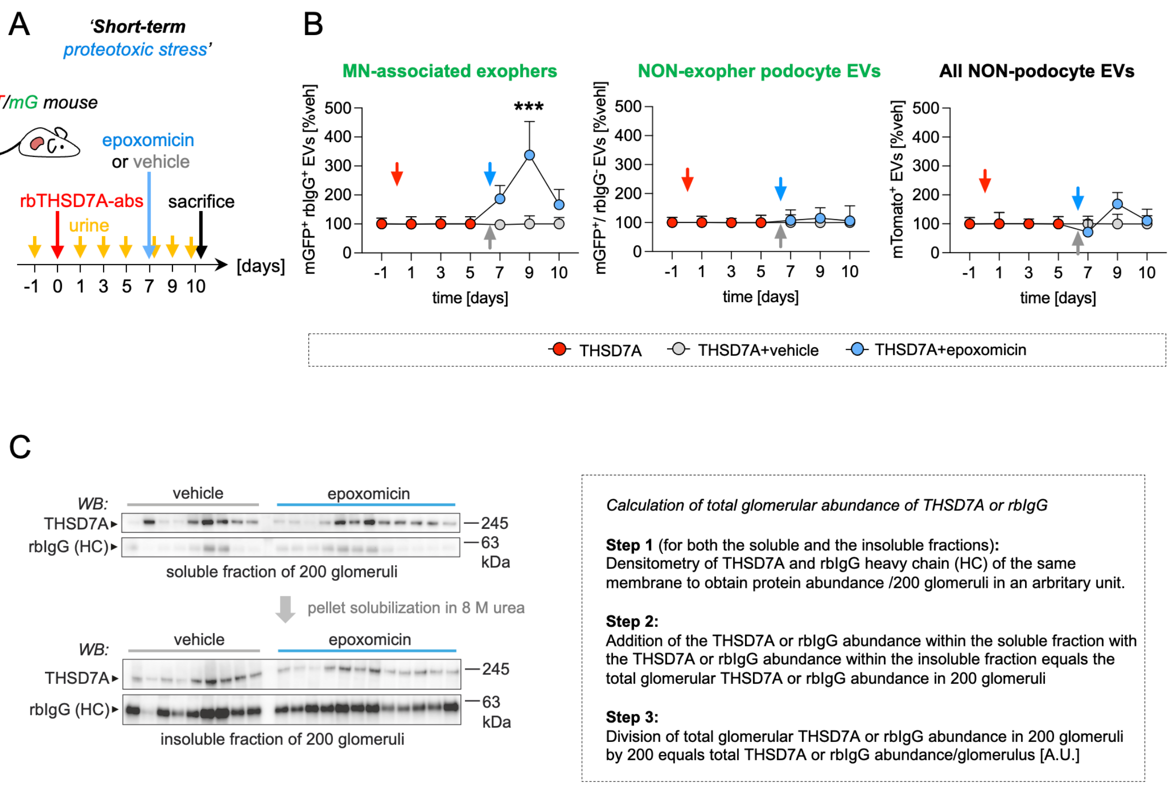

**'Short-term' proteotoxic stress induces specific exopher-release to the urine. A)** Experimental setup. On day 0, THSD7A$^+$-MN was induced in *mT/mG* reporter and BALB/c mice. 'Short-term' proteotoxic stress was achieved by administration of 0.5 µg/g bodyweight epoxomicin solubilized in 125 µl DMSO (light blue arrow) or 125 µl vehicle (DMSO; grey arrow) every day after day 7 for 4 consecutive days. Urine was collected prior to and in the time course of THSD7A$^+$-MN development. Urinary extracellular vesicles (EVs) were isolated from 250 µl urine and $10^9$ total EVs were analyzed by image stream to determine the abundance of rbIgG$^+$-EVs in particles/ml. At the end of the observation period, kidneys were removed and perfused through the renal artery for removal of circulating immunoglobulins and for magnetic bead perfusion for glomeruli isolation. **B)** Quantification of EV-subpopulation release to the urine in response to epoxomicin treatment relative to vehicle-THSD7A$^+$-MN *mT/mG* mice; mean ± SEM, *N* = 7 per group, pooled data from 2 independent experiments, $^{**}p \leq 0.01$, 2-way ANOVA with Tukey`s multiple comparison test. *Left panel:* Relative release of MN-associated exophers from podocytes; *Middle panel:* Relative release of podocyte EVs without bound rbIgG (non-exopher podocyte EVs); *Right panel:* Relative release of non-podocyte EVs (mTomato$^+$ EVs). Note the specific increase in urinary release of MN-associated exophers, whereas the NON-exopher podocyte EVs are not affected in the setting of proteasome inhibition. **C)** Glomeruli were isolated and counted. 200 glomeruli were processed to a soluble and insoluble fraction per mouse and loaded onto separate reducing SDS-PAGEs. THSD7A and rbIgG heavy chain (HC) protein abundance was assessed by immunoblotting of the same membrane; *N* = 9 vehicle treated THSD7A$^+$-MN mice, *N* = 12 epoxomicin treated THSD7A$^+$-MN mice. Total glomerular abundances were calculated as indicated and correlated to the respective urinary release of exophers at the end of the observation period (mean of day 9 and 10 exopher counts per 250 µl urine) of the individual mice.

### *Figure S18*

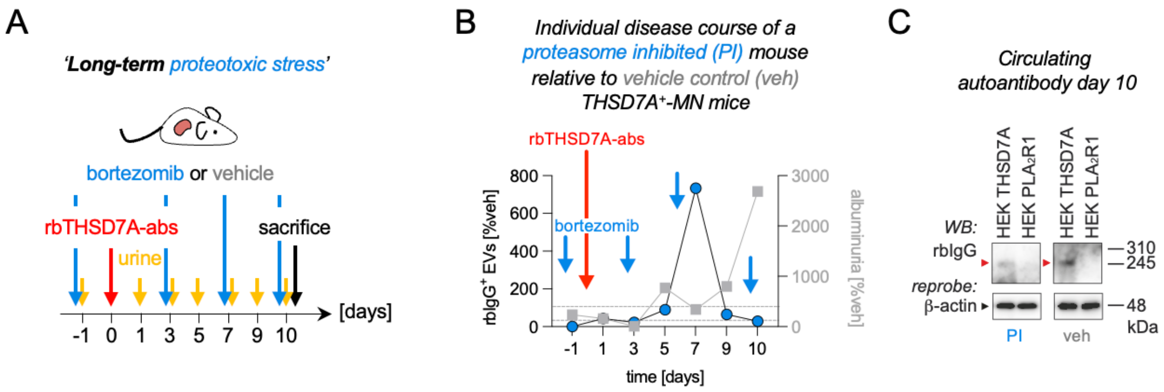

**'Long-term' proteotoxic stress decreases urinary exopher-release despite persistence of rbTHSD7A-abs in the serum of a THSD7A$^+$-MN mouse. A)** Experimental setup: On day 0, THSD7A$^+$-MN was induced in mice. 'Long-term' proteotoxic stress was achieved by administration of bortezomib (0.5 µg/g bodyweight solubilized in 125 µl DMSO; blue arrow) one day prior to THSD7A$^+$-MN induction by intra-peritoneal injection 2 times per week for 10 days. Vehicle control mice received 125 µl DMSO. Urine was collected prior to and in the time course of THSD7A$^+$-MN development. Serum was collected at the end of the observation period. **B)** Graph depicts the quantification of urinary exopher (rbIgG$^+$-EVs) release (left *y* axis, round colored symbols) and corresponding albuminuria (right *y* axis, squared grey symbols) in one individual proteasome inhibited (PI) mouse. Values of this mouse are expressed as relative changes to vehicle treated THSD7A$^+$-MN control mice ($N$ = 4); grey dashed lines indicate the 100% of vehicle treated THSD7A$^+$-MN control mice. Note the decrease in albuminuria upon increased exopher-release and the increase of albuminuria upon decreased exopher-release in the PI mouse. **C)** Circulating rbTHSD7A-abs are still detectable in the serum of the PI mouse and a representative vehicle mouse by using the serum 1:2 diluted as primary antibody on HEK cell lysate containing human THSD7A or human $PLA_2R1$ protein. This demonstrates that the decrease in urinary exopher release in the PI mouse is not the result of an absence of specific autoantibodies in the serum.

### *Figure S19*

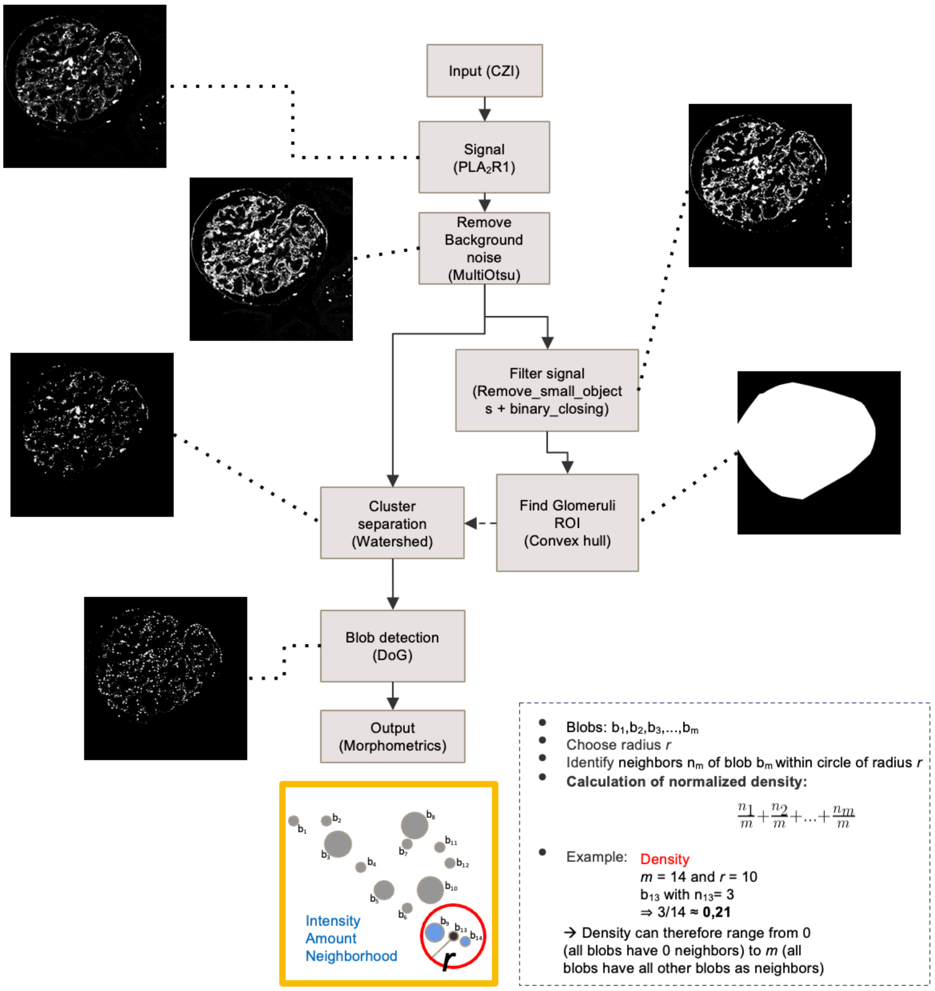

**Computational image analyses of PLA$_2$R1 aggregate density.** The algorithm was implemented in Python 3 (mainly packages Scikit-image and OpenCV) using a Difference of Gaussian blob detection. Algorithm inputs were images saved in Carl Zeiss Image (CZI) Data file format. Using the grayscale PLA$_2$R1 channel, images were first thresholded with a Multi-Otsu approach (Liao, P-S., Chen, T-S. and Chung, P-C., "A fast algorithm for multilevel thresholding", Journal of Information Science and Engineering 17 (5): 713-727, 2001.) separating high signal areas from background noise. The binary image was further processed removing very small noisy pixels and applying the morphological operation of closing. Based on the now cleared binary image a glomerular mask was extracted by calculating the convex hull around the binary signal and therefore roughly following the Bowman's capsules' outline. In the masked area signal clusters (areas of overlapping or touching signal blobs) were separated from each other using a watershed transformation (Peer Neubert & Peter Protzel (2014). Compact Watershed and Preemptive SLIC: On Improving Trade-offs of Superpixel Segmentation Algorithms. ICPR 2014, pp 996-1001). In the last step, the binary output of the watershed separation is used as input for the Difference of Gaussian blob detection. Based on the detected blobs the PLA$_2$R1 density for each blob is calculated. PLA$_2$R1 density is defined as the sum of the number of neighboring blobs (excluding the current blob) within a radius of 50 pixels around each blob divided by total number of blobs in the image. The division normalizes the image to a range from 0 (all blobs have 0 neighbors) to the total number of blobs (all blobs have all blobs as neighbors). The PLA$_2$R1 aggregated density per patient is calculated as the median of all PLA$_2$R1 densities of all images of that patient. All code will be made available via GitHub.

### *Figure S20*

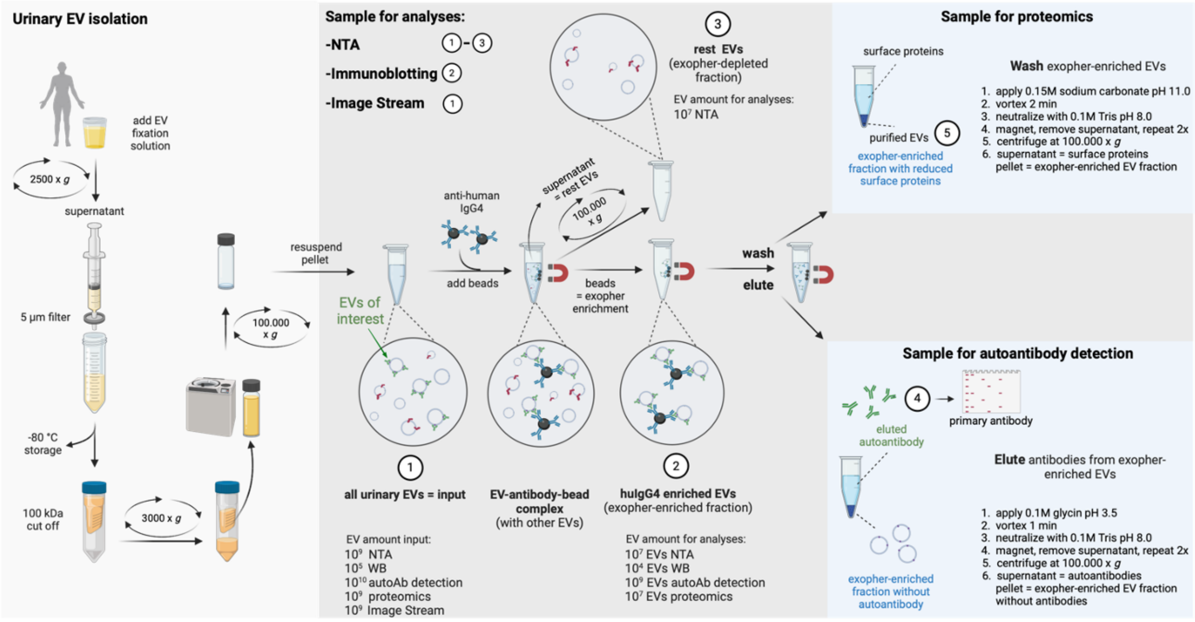

**General workflow of EV sample preparation from patient urine as detailed in the method section.** Illustration was created with Biorender.

### *Table S1*

**Supplementary Table 1. Blood and urine parameters of patients with membranous nephropathy at time points of analyses (U1= diagnostic samples; Ux = follow up samples)**

| | THSD7A⁺-MN -P1 | | | | | | | | $PLA_2R1^+$-MN-P2 | | $PLA_2R1^+$-MN-P3 | $PLA_2R1^+$-MN-P4 |
|---|---|---|---|---|---|---|---|---|---|---|---|---|
| gender | male | | | | | | | | female | | male | male |
| age | 62 | 62 | 63 | 64 | 64 | 64 | 64 | 64 | 19 | 21 | 80 | 68 |
| collection time point | U1 | U2 | U3 | U4 | U5 | U6 | U7 | U8 | U1 | U2 | Ux | U1 |
| GFR (CKD-EPI; ml/min) | >90 | 64 | 16 (-) | 33 (-) | 29 (-) | 29 (-) | 31 (-) | 41 (-) | >90 | >90 | 30 (-) | >90 |
| blood urea nitrogen (mg/dL) | 54 | 71 (+) | 130 (+) | 92 (+) | 81 (+) | 79 (+) | 82 (+) | 65 (+) | 17 | 20 | 113 (+) | 22 |
| serum triglycerides (mg/dL) | 310 (+) | 183 (+) | 218 (+) | 147 | 211 (+) | 195 (+) | 121 | 90 | 226 (+) | | 112 | |
| serum cholesterol (mg/dL) | 442 (+) | 286 (+) | 307 (+) | 173 | 213 (+) | 214 (+) | 185 | 147 | 380 (+) | | 147 | |
| serum albumin (g/L) | 12.4 (-) | 11.0 (-) | 10.6 (-) | 23.1 (-) | 19.3 (-) | 22.4 (-) | 28.9 (-) | 30.8 (-) | 17.3 (-) | 35.9 | 27.9 (-) | 34.1 |
| serum THSD7A titer (IFT) | 1:160 | 1:160 | 1:160 | Neg. | 1:10 | Neg. | Neg. | Neg. | Neg. | | Neg. | Neg. |
| serum $PLA_2R1$-AutoAK (U/ml) | Neg. | 0.8 | | | | | | | 92.5 (+) | Neg. | 935.2 (+) | 72.5 (+) |
| urine creatinine (g/L) | 1.11 | 0.15 (-) | 0.44 | 0.91 | 0.62 | 0.92 | 1.40 | 0.13 (-) | 0.79 | 1.57 (+) | 0.76 | 0.22 (-) |
| urine protein (mg/L) | 14852 (+) | 3172 (+) | 8215 (+) | 12099 (+) | 8674 (+) | 12987 (+) | 11551 (+) | 821 (+) | 10647 (+) | 2582 (+) | 12241 (+) | 202 (+) |
| urine albumin (mg/L) | 10500 (+) | 2252 (+) | 4138 (+) | 9003 (+) | 6853 (+) | 9141 (+) | 8583 (+) | 617 (+) | 7452 (+) | 1760 (+) | 8727 (+) | 145 (+) |
| urine protein-creatinine (mg/g) | 13376 (+) | 20571 (+) | 18670 (+) | 13265 (+) | 13885 (+) | 14186 (+) | 8228 (+) | 6436 (+) | 13474 (+) | 1640 (+) | 16003 (+) | 916.9 (+) |
| urine albumin-creatinine (mg/g) | 9456 (+) | 14604 (+) | 9404 (+) | 9870 (+) | 10970 (+) | 9984 (+) | 6114 (+) | 4839 (+) | 9430 (+) | 1118 (+) | 11409 (+) | 658.8 (+) |
| urine IgG (mg/L) | 473 (+) | | | | | | | | | 57 (+) | | |
| urine α2-macroglobulin (mg/L) | 2.93 | | | | | | | | | <2.50 | | |
| urine spec. weight | 1.105 | | | | | | | | | 1.005 | 1.053 | |
| urine pH | ~6 | | | | | | | | | ~6 | ~5 | |
| urine ketones (mg/dL) | Neg. | | | | | | | | | Neg. | Neg. | |
| urine urobilinogen (mg/dL) | Neg. | | | | | | | | | Neg. | Neg. | |
| urine bilirubin (mg/dL) | Neg. | | | | | | | | | Neg. | Neg. | |
| urine glucose (mg/dL) | Neg. | | | | | | | | | ~100 (+) | 100 (+) | |
| urine nitrite | Neg. | | | | | | | | | Neg. | Neg. | |
| urine erythrocytes (per µL) | ~25 (+) | | | | | | | | | ~10 (+) | ~25 (+) | |
| urine leukocytes (per µL) | Neg. | | | | | | | | | ~100 (+) | Neg. | |

**Abbreviations:** MN = membranous nephropathy; Ux = urine sample; P = patient; Neg. = negative; (+) = elevated; (-) reduced; U = units; ~ = approximately

### Table S2

| **Supplementary Table 2.** Blood and urine parameters of nephrotic control patients at time points of analyses (U1 = diagnostic samples; Ux = follow up sample) | | | | | | |
|---|---|---|---|---|---|---|
| | **MCD-P5** | **MCD-P6** | **IgAN-P7** | **Tubulo-toxic kidney injury-P8** | **pFSGS-P9** | **pFSGS-P10** |
| gender | male | female | female | male | male | male |
| age | 47 | 82 | 58 | 64 | 68 | 54 |
| collection time point | U1 | U1 | U1 | U1 | U1 | Ux |
| GFR (CKD-EPI; ml/min) | >90 | 12 (-) | 25 (-) | 13 (-) | 15 (-) | 28 (-) |
| blood urea nitrogen (mg/dL) | 26 | 258 (+) | 72 (+) | 174 (+) | 150 (+) | 65 (+) |
| serum triglycerides (mg/dL) | 96 | 255 (+) | | | 248 (+) | 93 |
| serum cholesterol (mg/dL) | 209 (+) | 221 (+) | | | | 197 |
| serum albumin (g/L) | 20.7 (-) | 14.8 (-) | 41.7 | 23.4 (-) | 23.5 (-) | 33 (-) |
| serum THSD7A titer (IFT) | | Neg. | Neg. | | | |
| serum $PLA_2R1$-AutoAK (U/ml) | Neg. | Neg. | Neg. | Neg. | Neg. | Neg. |
| urine creatinine (g/L) | 0.16 | 0.78 | | 0.47 | 0.41 | 0.80 |
| urine protein (mg/L) | 918 (+) | 13688 (+) | | 3390 (+) | 2591 (+) | 3432 (+) |
| urine albumin (mg/L) | 780.4 (+) | 9481 (+) | 2565 (+) | 1690 (+) | 1942 (+) | 2684 (+) |
| urine protein-creatinine (mg/g) | 5885 (+) | 17441 (+) | | 7217 (+) | 6272 (+) | 4294 (+) |
| urine albumin-creatinine (mg/g) | 5003 (+) | 12080 (+) | 3103 (+) | 3598 (+) | 4001 (+) | 3358 (+) |
| urine IgG (mg/L) | | | | | | |
| urine $\alpha$2-macroglobulin (mg/L) | | | | | | |
| urine spec. weight | | | | | | |
| urine pH | | | | | | |
| urine ketones (mg/dL) | | | | | | |
| urine urobilinogen (mg/dL) | | | | | | |
| urine bilirubin (mg/dL) | | | | | | |
| urine glucose (mg/dL) | | | | | | |
| urine nitrite | | | | | | |
| urine erythrocytes (per µL) | | | | | | |
| urine leukocytes (per µL) | | | | | | |

**Abbreviations:** MCD = minimal change disease; IgAN = IgA nephritis; pFSGS = primary focal segmental glomerulosclerosis; Ux = urine sample; P = patient; Neg. = negative; (+) = elevated; (-) reduced; U = units; ~ = approximately

### Table S3

| **Supplementary Table 3.** HuIgG4⁺-EV characteristics in the diagnostic urines (U1) of nephrotic patients. | | | | | | | |
|---|---|---|---|---|---|---|---|
| **Characteristic** | | **THSD7A⁺-MN** | **$PLA_2R1^+$-MN** | **MCD** | **pFSGS** | **IgAN** | **Tubulo-toxic kidney injury** |
| huIgG4⁺/14-3-3⁺ EV abundance (%) | *IS* | 35 | 3 | 25 | 1 | 10 | 0.5 |
| vesicular characteristics | | | | | | | |
| mean size (nm) | *NTA* | ~600 | ~500 | ~400 | ~400 | ~350 | ~250 |
| max size (nm) | *IF* | ~8000 | ~8000 | ~5000 | ~2000 | ~5000 | ~2000 |
| marker proteins | *WB, IS* | 14-3-3<br>Annexin A1<br>CD63/CD81<br>CD9<br>Flotillin<br>TSG101 | 14-3-3<br>Annexin A1<br>CD63/CD81<br>CD9<br>Flotillin<br>TSG101 | CD9<br>14-3-3<br>Flotillin<br>TSG101 | TSG101<br>CD9<br>Annexin A1<br>Flotillin<br>CD63 | 14-3-3<br>Annexin A1<br>CD63/CD81<br>CD9<br>Flotillin<br>TSG101 | CD9<br>Annexin A1<br>CD63/CD81 |
| vesicular content | | | | | | | |
| autoantibody | | anti-THSD7A | anti-$PLA_2R1$ | | | | |
| podocyte proteins | *WB* | THSD7A<br>$PLA_2R1$<br>Nephrin<br>$\alpha$-Actinin-4<br>Podocin<br>Synaptopodin | $PLA_2R1$<br>THSD7A<br>Nephrin<br>$\alpha$-Actinin-4<br>Podocin<br>Synaptopodin | Nephrin<br>$\alpha$-Actinin-4<br>Synaptopodin | Podocin<br>Synaptopodin<br>$\alpha$-Actinin-4<br>THSD7A<br>Nephrin | $\alpha$-Actinin-4<br>Podocin<br>Synaptopodin | absent |
| mitochondrial proteins | *WB* | abundant | abundant | present | present | present | absent |
| lysosomal proteins | *WB* | abundant | abundant | present | present | present | present |
| UPS proteins | *WB* | abundant | abundant | present | present | present | present |
| complement proteins | *WB* | abundant | abundant | absent | absent | absent | absent |

**Abbreviations:** *IS* = image stream; *NTA* = nanoparticle tracking analyses; *IF* = immunofluorescence; *WB* = immunoblot; ~ = approximately

### *Table S4*

Proteomic data, please refer to the Excel list.

### *Table S5*

**Supplementary Table 5. Antibodies and dyes used in the study.**

| | clone/catalogue number | company | IF/IHC | WB | IS | iEM | PD | *in vitro* A | *In vivo* A |
|---|---|---|---|---|---|---|---|---|---|
| rb anti-THSD7A | #HPA000923 | Atlas | 1:200, 1:400 | 1:1000 | 1:50 | | | | |
| hu anti-THSD7A | reference[19] | self-made | | 1:1000 | | 1:20 | | 1:100 | |
| rb anti-THSD7A | reference[10] | self-made | | | | | | 1:200 | |
| ms anti-THSD7A mAb | Seifert L. et al KI in press | self-made | 1:50 | | | | | | |
| control rabbit IgG | #I5006 | Sigma | | | | | | | 1:200 |
| ms anti-myc | #sc-41 | Santa Cruz | | 1:1000 | | | | | |
| rb anti-$PLA_2R1$ | #HPA012657 | Thermo Fisher | 1:200 | 1:1000 | 1:50 | | | | |
| gp anti-Nephrin | #GP-N2 | Progen | 1:200 | 1:2000 | 1:50 | | | | |
| rb anti-$\alpha$-Actinin 4 | #0042-05 clone IG-701 | ImmunoGlobe | | 1:1000 | 1:50 | | | | |
| gt anti-Annexin A1 | #AF3770 | R&D Systems | 1:400 | 1:1000 | | | | | |
| rb anti-Annexin A1 | ab214486 | Abcam | | 1:2000 | | | | | |
| APC anti-Annexin A1 | #BLD640912 | Novus Biologicals | | | 1:10 | | | | |
| CoraLight Plus 647 anti-Annexin V | #CL647-66245 | ProteinTech | | | 1:10 | | | | |
| Pacific Blue anti-CD63 | #BLD-353011/353012 | BioLegend | | | 1:50 | | | | |
| Pacific Blue anti-CD81 | #BLD-349516 | BioLegend | | | 1:50 | | | | |
| ms anti-Flotillin | #610821 | BD Bioscience | | 1:1000 | | | | | |
| rb anti-14-3-3 AF546 | #Sc-1657 AF546 pan 14-3-3- H8 | Santa Cruz | | | 1:50 | | | | |
| rb anti-14-3-3 | # Sc-1657 Pan 14-3-3 H8 | Santa Cruz | | 1:1000 | | | | | |
| ms anti-$\beta$-actin | #A5441 clone AC-15 | Sigma | | 1:10.000 | | | | | |
| cy2-dk anti-human IgG | #709-225-149 | Jackson ImmunoResearch Laboratories | 1:100 | | | | | | |
| rb anti-phospho-Paxillin | #44722G | Thermo Fisher | 1:100 | | | | | | |
| cy5 rb anti-rabbit IgG | #711-175-152 | Jackson ImmunoResearch Laboratories | 1:200 | | 1:200 | | | | |
| cy2-dk anti-human IgG | #709-225-149 | Jackson ImmunoResearch Laboratories | 1:200 | | 1:200 | | | | |
| CoraLite® 488 anti-human IgG4 | #CL488-66408 | ProteinTech | 1:100 | | 1:50 | | | | |
| ms anti-human IgG4 | #9190-01 | Southern-Biotech | | | | | | | |
| ms anti-human IgG4 | #9200-09 | Southern-Biotech | 1:4000 | | | | | | |
| gt anti-human IgG | #209-005-088 | Jackson ImmunoResearch Laboratories | 1:7500 | | | | | | |
| 12 nm Colloidal Gold AffiniPure™ Goat Anti-Human IgG (H+L) (EM Grade) | #109-205-088 | Jackson ImmunoResearch | | | | 1:10 | | | |
| 12 nm Colloidal Gold AffiniPure™ Donkey Anti-Rabbit IgG (H+L) (EM Grade | #711-205-152 | Jackson ImmunoResearch | | | | 1:50 | | | |
| rhodamine-wheat germ agglutinin (WGA) | #RL-1022 | Vector | 1:400 | | | | 1:20 | | |
| biotin-WGA | #B-1025-5 | Vector | 1:400 | | | | | | |
| AF647-streptavidin | #S21374 | Molecular Probes | 1:200 | | | | | | |
| rb anti-C1q | A0136 | Dako | 1:200 | | | | | | |
| rb anti-Podocin | #P0372 | Sigma | | 1:1000 | 1:10 | | | | |
| rb anti-Mitofusin-2 | #9482 | Cell Signaling | 1:200 | 1:1000 | 1:50 | | | | |
| rb anti-MnSOD | #06-984 | Millipore | 1:100 | | | | | | |
| ms anti-ATPB | #ab14730 | Abcam | | 1:1000 | | | | | |
| rb anti-C5b9 | #M0777 | Dako | 1:100 | | | | | | |
| rb anti-C5b9 | #ab55811 | Abcam | | 1:1000 | | | | | |
| gt anti-Cathepsin B | #PA5-47975 | Invitrogen | 1:200 | | | | | | |

| | | | | | |
|---|---|---|---|---|---|
| rb anti-LAMP2 | #L0668 | Sigma | | 1:1000 | |
| rb anti-LIMP2 | | Paul Saftig, CAU Kiel, Germany | 1:500 | 1:1000 | 1:50 |
| ms anti-α3 | #sc-166205 | Santa Cruz | | 1:1000 | |
| rb anti-CD63 | #ab216130 | Abcam | | 1:1000 | |
| rb anti-CD9 | #ab92726 | Abcam | | 1:1000 | |
| rb anti-k48pUB | #05-1308 | Millipore | | 1:1000 | 1.50 |
| rb anti-k48pUB | #ab140601 | Abcam | 1:300 | | |
| rb anti-LMP7 | #ab3329 | Abcam | | 1:1000 | |
| rat anti-UCH-L1 mAb | reference[32] | self-made | | 1:250 | |
| MemBrite® Fix Cell Surface dye 594/615 | #30096 | Biotium | | | 1x |
| MitoSOX™ | #M36008 | Invitrogen | 10 µM | | 10 µM |
| LysoView™ 488 | #77067 | Biotium | 1:20 | | 1:20 |
| Hoechst 33342 | #H3570 | Thermo Fisher | 1:1000 | | |
| rat anti-UCH-L1 mAb | reference[32] | self-made | | 1:250 | |
| MemBrite® Fix Cell Surface dye 594/615 | #30096 | Biotium | | | 1x |
| MitoSOX™ | #M36008 | Invitrogen | 10µM | | 10 µM |
| LysoView™ 488 | #77067 | Biotium | 1:20 | | 1:20 |
| Hoechst 33342 | #H3570 | Thermo Fisher | 1:1000 | | |

All secondary antibodies used were either biotinylated, HRP- gold-, or fluorescent dye-conjugated affinity purified donkey antibodies (Jackson ImmunoResearch).

**Abbreviations:** *IS* = image stream; *IF* = immunofluorescence; *WB* = immunoblot; *iEM* = Immunogold EM; *PD* = Pulldown; *A* = application; *rb* = rabbit, *ms* = mouse; *gt* = goat; *ha* = hamster; *gp* = guinea-pig; *dk* = donkey, *mAb* = monoclonal antibody
